# 53BP1 deficiency leads to hyperrecombination using break-induced replication (BIR)

**DOI:** 10.1101/2024.09.11.612483

**Authors:** Sameer Bikram Shah, Youhang Li, Shibo Li, Qing Hu, Tong Wu, Yanmeng Shi, Tran Nguyen, Isaac Ive, Linda Shi, Hailong Wang, Xiaohua Wu

## Abstract

Break-induced replication (BIR) is mutagenic, and thus its use requires tight regulation, yet the underlying mechanisms remain elusive. Here we uncover an important role of 53BP1 in suppressing BIR after end resection at double strand breaks (DSBs), distinct from its end protection activity, providing insight into the mechanisms governing BIR regulation and DSB repair pathway selection. We demonstrate that loss of 53BP1 induces BIR-like hyperrecombination, in a manner dependent on Polα-primase-mediated end fill-in DNA synthesis on single-stranded DNA (ssDNA) overhangs at DSBs, leading to PCNA ubiquitination and PIF1 recruitment to activate BIR. On broken replication forks, where BIR is required for repairing single-ended DSBs (seDSBs), SMARCAD1 displaces 53BP1 to facilitate the localization of ubiquitinated PCNA and PIF1 to DSBs for BIR activation. Hyper BIR associated with 53BP1 deficiency manifests template switching and large deletions, underscoring another aspect of 53BP1 in suppressing genome instability. The synthetic lethal interaction between the 53BP1 and BIR pathways provides opportunities for targeted cancer treatment.

## Introduction

DNA double strand breaks (DSBs) present a major threat to genome stability, and improper repair of DSBs can result in chromosomal rearrangements and susceptibility to cancer ^1,2^. Multiple DSB repair pathways can be used to repair DSBs, which are classified into two main groups: end joining and homology-directed recombination (HDR). While KU-dependent canonical nonhomologous end joining (NHEJ) serves as the major end joining pathway, microhomology-mediated end joining (MMEJ) emerges as an alternative pathway, potentially more relevant to cancer etiology, as evidenced by the frequent occurrence of microhomology sequences at cancer breakpoints ^3–5^. Homologous recombination (HR), often referred to as short tract gene conversion (STGC), is mainly carried out through synthesis-dependent strand annealing (SDSA) in mitotic cells ^6–8^. HR is considered as the most precise DSB repair mechanism due to its utilization of homologous templates. The other two HDR repair pathways, break-induced replication (BIR) and single-strand annealing (SSA), are error-prone and mutagenic ^9–12^.

BIR is utilized when one DSB end successfully invades the homologous template site but has difficulty in catching the other end after new DNA synthesis, which typically occurs when DSBs are single-ended or the second end is situated too far away ^9–11,13^. BIR plays an important role in repairing broken replication forks and eroding telomeres, where single-ended DSBs (seDSBs) are often generated ^13,14^. In mammalian cells, BIR is also shown to be involved in mitotic DNA synthesis [MiDAS ^13,15,16^]. In BIR, after strand invasion, a migrating replication bubble is established and BIR DNA synthesis is driven by branch migration, resulting in the conservative inheritance of newly synthesized DNA ^9^. While studies in yeast revealed that BIR can proceed for a long distance (∼100 kb) to the end of the chromosomes ^17,18^, BIR tract length in mammalian cells appears to be short (< 4kb) at double-ended DSB (deDSBs) generated by endonucleases ^19^. Different from the general HR (STGC), BIR requires Pol32/POLD3 and helicase PIF1 ^19–25^. Compared to S-phase DNA replication, BIR exhibits a 100- to 1000-fold increase in mutation rate ^25,26^. BIR also promotes template switching ^27^, which could cause gross chromosomal rearrangements that are frequently observed in the cancer genome ^28,29^. Therefore, while serving as an important DSB repair mechanism, particularly to deal with challenging situations such as replication stress and telomere damage, BIR activity should be restricted unless its use is necessary. However, the mechanism to regulate the use of BIR remains elusive.

53BP1 is a chromatin-binding protein that protects DNA ends from excessive end resection through the recruitment of its downstream effectors RIF1 and the shieldin complex (SHLD1/SHLD2/SHLD3/REV7) ^30–32^, thereby promoting NHEJ and limiting HR ^33^. The role of 53BP in NHEJ is supported by its importance for NHEJ-mediated immunoglobulin class switch ^34,35^ and the fusion of dysfunctional telomeres ^36^. 53BP1 and BRCA1 compete for DSB end binding to regulate the usage of NHEJ and HR ^37–40^. Importantly, the loss of 53BP1 or its downstream effectors induces PARP inhibitor (PARPi) resistance in BRCA1-deficent tumors, at least in part by restoring HR activity ^33,38–52^. It has also been shown that 53BP1 deficiency shifts HR repair towards SSA for DSB repair, leading to the proposal that SSA could be used as one mechanism to rescue the HR defect in BRCA1-deficient cells ^53^.

In our previous study, we showed that HR (STGC) is preferentially used at deDSBs, while BIR is activated and predominantly utilized at seDSBs on broken forks ^19^. However, the regulatory mechanism that suppresses BIR at deDSBs remains unclear. In this study, we detect a hyperrecombination activity at deDSBs when 53BP1 is deficient, using the BIR mechanism in a manner dependent on the BIR key players POLD3 and PIF1. We propose that loss of 53BP1 not only impairs NHEJ at deDSBs, but also leads to BIR-like hyperrecombination that is critical for rescuing the HR defect in BRCA1-deficient cells. Induction of BIR-like hyperrecombination in 53BP1-deficient cells is not primarily governed by hyper end resection but instead promoted by the overloading of ubiquitinated PCNA (PCNA-Ub) and PIF1 at deDSBs, which depends on Polα-primase-mediated end fill-in DNA synthesis on ssDNA overhangs. These studies have unveiled mechanisms underlying the control of BIR at deDSBs and have suggested a role for 53BP1 in controlling DSB repair pathway selection after end resection. We also show that upon fork breakage, the chromatin remodeling protein SMARCAD1 antagonizes 53BP1 binding to DSBs on the broken forks to facilitate BIR. Furthermore, cells deficient in 53BP1 exhibit reliance on BIR for survival, irrespective of BRCA1 status, establishing a therapeutic strategy by targeting the BIR key player PIF1 to treat not only PARPi-resistant BRCA1-deficient tumors resulting from a compromised 53BP1 pathway, but also tumors with TIRR overexpression leading to 53BP1 inactivation.

## Results

### Loss of 53BP1 results in a substantial increase of BIR

It has been proposed that 53BP1 protects DSB ends, thereby facilitating NHEJ while inhibiting HR ^30,32,54^. As expected, HR/STGC is substantially increased in U2OS (EGFP-HR/STGC) reporter cell line when 53BP1 or RIF1 is depleted or when 53BP1 is knocked out (Fig. 1a and S1a). To test whether 53BP1 deficiency also modulates BIR, we depleted 53BP1 in our established U2OS (EGFP-BIR/LTGC) reporter cell line [Fig. 1b ^19^] and found that the percentage of EGFP-positive cells is significantly increased after I-SceI cleavage (Fig. 1c left and S1b left). Similar results were obtained in *53BP1*-KO U2OS (EGFP-BIR/LTGC) cells (Fig. 1c middle and S1b middle). Increased percentage of EGFP-positive cells observed in *53BP1*-KO or 53BP1-depleted cells shows dependence on POLD3, PIF1, BRCA1 and RAD51, resembling the pattern seen in wildtype (WT) U2OS (EGFP-BIR/LTGC) cells (Fig. 1d, S1c and S16), confirming that BIR is substantially increased in 53BP1-deficient cells. However, BIR/LTGC scored by our reporter does not require RAD52 in both WT and *53BP1*-KO cells (Fig. 1e and S14b). Depleting the 53BP1 downstream effectors RIF1 and SHLD1 also causes an increase in BIR (Fig. 1c right, S1b right and S1d).

**Fig. 1.**
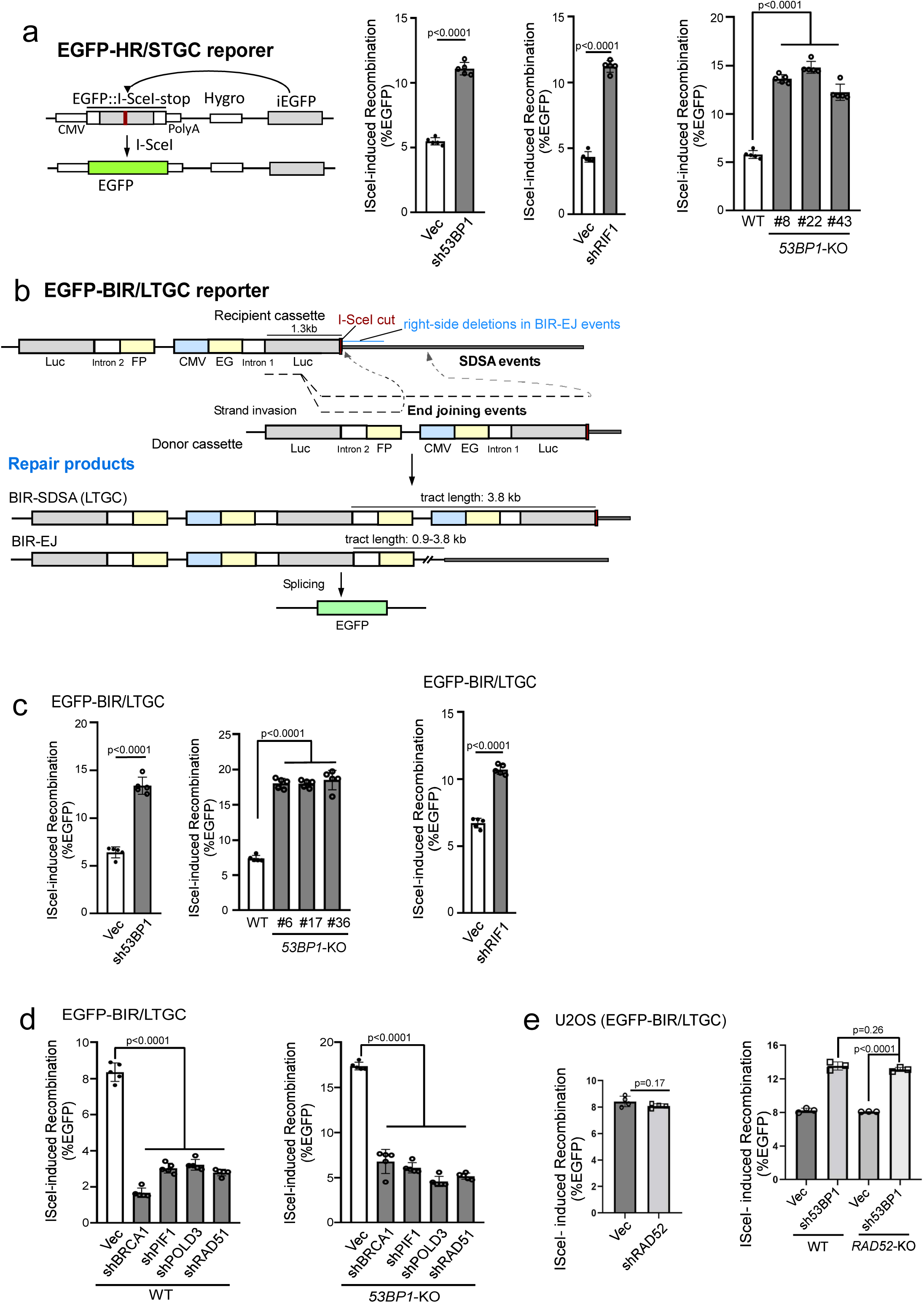
Inactivation of the 53BP1 pathway leads to hyperrecombination for both HR/STGC and BIR/LTGC. (a). Schematic drawing of the EGFP-HR/STGC reporter and the repair product after endonuclease I-SceI cleavage (left). The HR frequency was determined in U2OS (EGFP-HR/STGC) cells expressing shRNAs for 53BP1 and RIF1 (middle), or with *53BP1*-KO (three KO clones), following infection with lentiviruses encoding I-SceI endonuclease (right). The recombination frequency was determined by FACS analysis, 5 days post-infection. Western blot analysis was conducted to confirm 53BP1 and RIF1 knockdown and *53BP1* KO (Supplementary Figure 1a). (n=5 replicates) (b). Schematic drawing of the EGFP-BIR/LTGC reporter and the repair products using synthesis-dependent stand annealing (SDSA) or end joining (EJ) to complete BIR, termed as BIR-SDSA and BIR-EJ, respectively. The tract length for BIR-SDSA is 3.8 kb, and for BIR-EJ, it ranges from 0.9 kb to 3.8 kb. (c). U2OS (EGFP-BIR/LTGC) cells, infected with lentiviruses encoding shRNAs for 53BP1 (left) and RIF1 (right) with a vector control, or harboring *53BP1*-KO (middle), were assayed for BIR frequency by determining the percentage of EGFP positive cells with FACS analysis 5 days post-infection of I-SceI lentiviruses. The expression of 53BP1 and RIF1 is shown by Western blotting and qPCR, respectively (Supplementary Figure 1b). (n=5 replicates) (d and e). U2OS (EGFP-BIR/LTGC) WT or *53BP1*-KO cells stably expressing shRNAs targeting BRCA1, PIF1, POLD3 or RAD51 (d), or RAD52 (e, left), as well as U2OS (EGFP-BIR/LTGC) WT or *RAD52*-KO cells expressing 53BP1 shRNAs (e, right), were infected with lentiviruses encoding I-SceI, and the percentage of EGFP-positive cells was determined by FACS, 5 days post-infection. The expression of indicated proteins was examined by qPCR (Supplementary Figure 1c and S14b). (d n=5 replicates, e n=4 replicates)

We showed previously that at endonuclease-generated DSBs, BIR can be completed by SDSA (BIR-SDSA) or end joining (BIR-EJ) [Fig. 1b ^19^]. To initiate BIR after I-SceI cleavage in the EG-Luc cassette, the Luc sequence invades the homologous Luc sequence in the Luc-FP cassette on its sister chromatid, and if DNA synthesis on the invading strand can proceed for 3.8 kb to reach the right-side homology outside of the reporter, BIR is completed by SDSA (Fig. 1b, BIR-SDSA). However, if the replicating strand is prematurely disassociated from its template, the newly synthesized DNA end may be ligated to the other end of the original DSB and BIR is completed by end joining (Fig. 1b, BIR-EJ). In this case, if the invading strand has completed 0.9 kb DNA synthesis to reach the end of the intron-FP fragment, EGFP-positive cells can also be produced. Therefore, in our EGFP-BIR/LTGC reporter system, the BIR tract length for BIR-SDSA is 3.8 kb, whereas for BIR-EJ, it ranges from 0.9 kb to 3.8 kb.

To examine the mechanistic details of BIR in 53BP1-deficient cells, we analyzed the BIR products recovered from single EGFP-positive clones of WT and *53BP1*-KO U2OS (EGFP-BIR/LTGC) reporter cell lines after I-SceI cleavage. In agreement with the previous findings ^19^, the majority of BIR events at deDSBs are completed by BIR-EJ (80.0%) in WT cells, with most events containing microhomology (MH) at the end joining junctions (73.0%), suggesting that MMEJ is the major mechanism for end joining to complete BIR-EJ as assayed by our reporter (Fig. 2a and Supplementary Figure 2a). In *53BP1*-KO cells, the ratio of BIR-EJ versus BIR-SDSA and the percentage of BIR-EJ associated with MMEJ remain at similar levels as that in WT (Fig. 2a and Supplementary Figure 2a), suggesting that the loss of 53BP1 does not affect the use of SDSA or EJ to complete BIR. The BIR replication tract length also remains at similar levels in *53BP1*-KO cells compared to WT cells (Fig. 2a and 2b left; Fig. 1b, Repair products and Supplementary Figure 2a). However, the size of deletions at the right side of the I-SceI site in the recipient cassette of the EGFP-HR/BIR/LTGC reporter is significantly increased (Fig. 2a and 2b right, Fig. 1b top), consistent with more extensive end resection occurring in 53BP1-deficient cells, resulting in larger deletions associated with MMEJ to complete BIR. In yeast, BIR is associated with template switching ^27^, and microhomology-mediated jumping was also observed in the absence of Pif1 helicase ^55^. With our reporter system, we observed local jumping/template switching in the BIR-EJ events in WT U2OS cells (14.6%), with a notable, albeit not significant, increase in *53BP1*-KO cells (22.9%) (Fig. 2c top: table and Supplementary Figure 2b), and most jumping/template switching events contain MH sequences at the jumping sites in both WT and *53BP1*-KO cells (WT: 72.7%; *53BP1*-KO: 85.7%) (Fig. 2c), suggesting that MH sequences are often used to mediate template switching. Several examples of BIR jumping/template switching events in the context of the EGFP-BIR/LTGC reporter are shown in Fig. 2c, bottom panel.

**Fig. 2.**
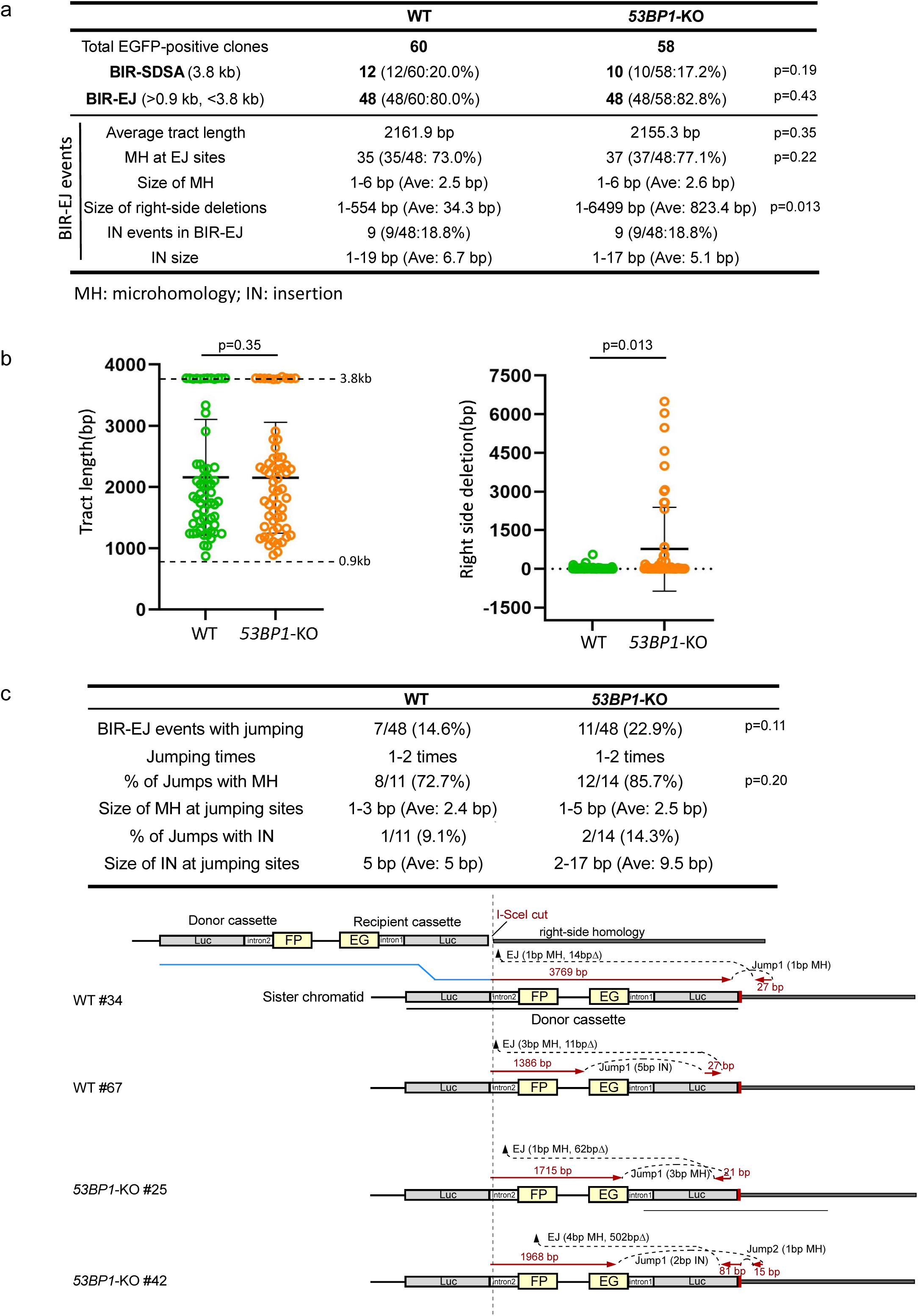
Analysis of BIR events in WT and *53BP1*-KO U2OS (EGFP-BIR/LTGC) cells. (a). A summary of the data from two sets of experiments displaying the features of BIR repair products from single EGFP-positive clones derived from WT and *53BP1*-KO (EGFP-BIR/LTGC) reporter cell lines after I-SceI cleavage. Genomic DNA extracted from single EGFP-positive clones was characterized by Sanger sequencing following PCR to determine the repair products and the repair junctions. The result of each set of experiments are shown in Supplementary Figure 2a. (b). The BIR tract length (left) and the size of the right-side deletions in BIR-EJ events (right) were analyzed in EGFP-positive single clones derived from U2OS (EGFP-BIR/LTGC) WT and *53BP1*-KO reporter cell lines after I-SceI cleavage. Group means are shown, and error bars represent ± SD. Dashed lines (3.8 kb and 0.9 kb) in the left panel indicate the upper and lower limits of the BIR tract length, respectively, that can be scored by the EGFP-BIR/LTGC reporter (see Fig. 1b). (WT n=60 clones, *53BP1*-KO n=58 clones) (c). A summary of the data from two sets of experiments on the BIR-EJ events with jumping/template switching, assayed in U2OS (EGFP-BIR/LTGC) WT and *53BP1*-KO reporter cell lines after I-SceI cleavage is show at the top. The result of each set of experiments are shown in Supplementary Figure 2b. Illustrations depicting examples of BIR-EJ events with jumping/template switching are presented at the bottom. In each example, the solid red lines represent the DNA synthesis tract copying the donor sequence, with the DNA synthesis length indicated. The red arrows represent the direction of DNA synthesis. The size of microhomology (MH) and insertion (IN) at each template jumping site, and at each end joining (EJ) site of the newly synthesized DNA ligated with the right end of the original DSB, is marked.

Collectively, our study reveals an important role of 53BP1 in constraining the use of BIR at deDSBs. Increased use of BIR in 53BP1-deficient cells would not only escalate the overall mutagenic outcome naturally associated with BIR, such as template switching, but also increase the risk of generating large deletions when BIR-EJ mechanism is used.

### Hyper HR/STGC observed in 53BP1-deficient cells exhibits dependence on PIF1 and POLD3

One unique feature of BIR/LTGC is its dependence on POLD3 and PIF1, whereas both factors are dispensable for HR/STGC at deDSBs ^19^. Consistently, when we depleted POLD3 and PIF1 in U2OS (EGFP-HR/STGC) reporter cells, HR/STGC remains unchanged after I-SceI cleavage (Fig. 3a left and S3a left). Interestingly, however, in *53BP1*-KO U2OS (EGFP-HR/STGC) cells, elevated HR at deDSBs is substantially reduced upon depletion of POLD3 and PIF1 (Fig. 3a right and S3a right), suggesting that hyperrecombination at deDSBs resulting from 53BP1 deficiency has changed the repair pathway, becoming dependent on PIF1 and POLD3 and using the BIR-like mechanism.

**Fig. 3.**
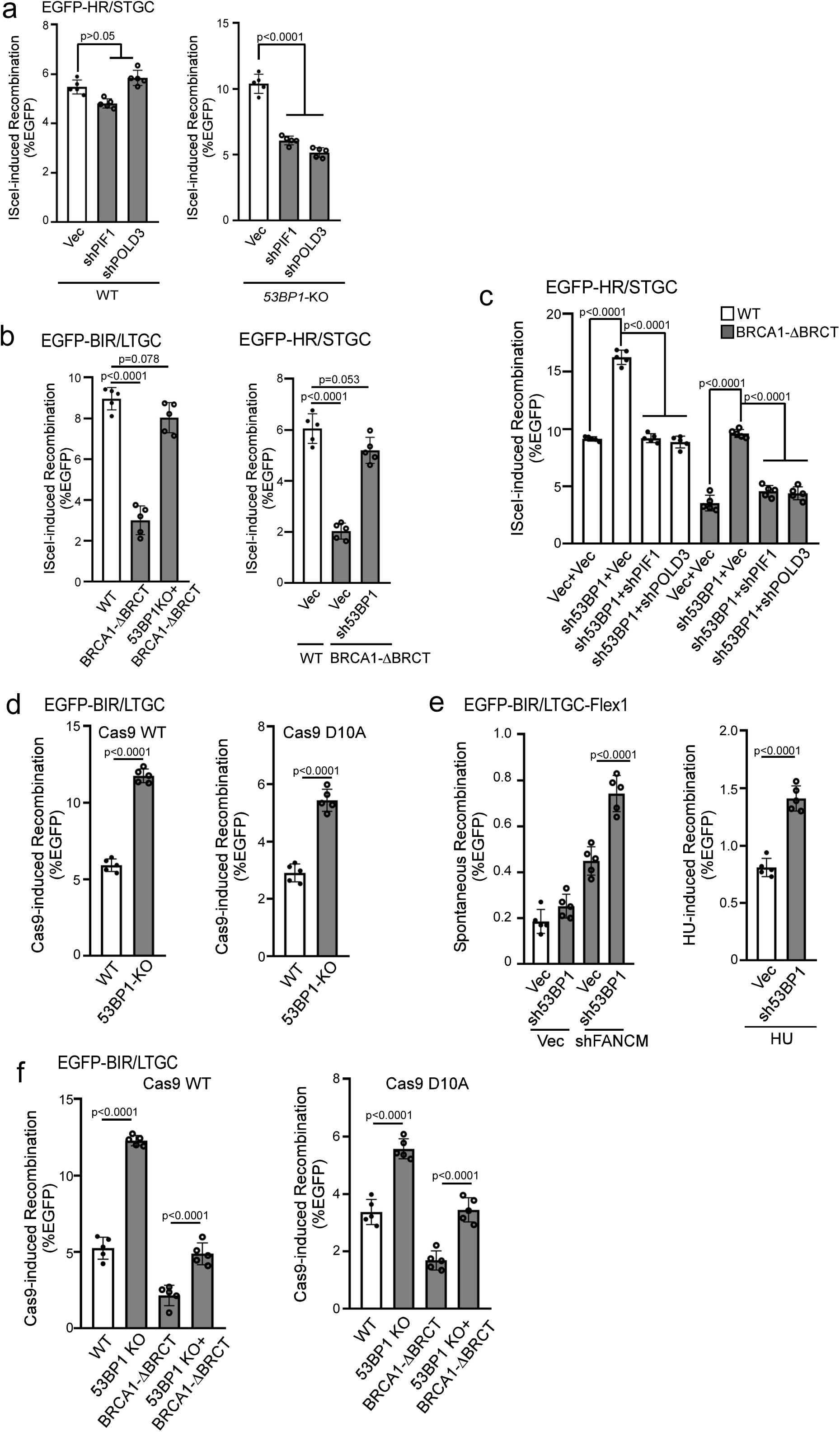
Hyperrecombination induced by 53BP1 deficiency uses the BIR mechanism. (a). U2OS (EGFP-HR/STGC) WT or *53BP1*-KO cells stably expressing shRNAs targeting PIF1 or POLD3 were infected with lentiviruses encoding I-SceI, and the percentage of EGFP-positive cells was analyzed by FACS, 5 days post-infection. The expression of PIF1 and POLD3 was examined by qPCR (Supplementary Figure 3a). (n=5 replicates) (b). BIR frequency in U2OS (EGFP-BIR/LTGC) WT, BRCA1-ΔBRCT and *53BP1*-KO/BRCA1-ΔBRCT cells (left), and HR frequency in U2OS (EGFP-HR/STGC) WT cells expressing vector, or in BRCA1-ΔBRCT cells expressing 53BP1 shRNA or vector (right), were determined by FACS analysis of EGFP positive cells, 5 days after I-SceI lentiviral infection. (n=5 replicates) (c). U2OS (EGFP-HR/STGC) WT and BRCA1-ΔBRCT cells were infected with lentiviruses expressing 53BP1 shRNA or vector, followed by a second round of lentiviral infection with shRNAs targeting PIF1 and POLD3 using vector as a control. HR frequency was determined by FACS analysis of EGFP positive cells, 5 days after I-SceI lentiviral infection. The expression of indicated proteins was examined by qPCR (Supplementary Figure 3c). (n=5 replicates) (d). BIR frequency was determined in U2OS (EGFP-BIR/LTGC) WT and *53BP1*-KO cells by FACS analysis 5 days after lentiviral infection of gRNA/Cas9^WT^ (left) or gRNA/Cas9^D10A^ (right). (n=5 replicates) (e). U2OS (EGFP-BIR/LTGC-Flex1) cells expressing 53BP1 shRNA with vector as a control were infected with lentiviruses encoding FANCM shRNA or vector (left), or synchronized to S-phase using double thymidine block followed by HU treatment (2 mM, 24 h, right). BIR frequency was determined by FACS analysis of EGFP positive cells 6 days after. The expression of FANCM was examined by qPCR (Supplementary Figure 3d). (n=5 replicates) (f). BIR frequency was determined by FACS analysis of EGFP positive cells in U2OS (EGFP-BIR/LTGC) WT, *53BP1*-KO, BRCA1-ΔBRCT and *53BP1*-KO/BRCA1-ΔBRCT cell lines, 5 days after lentiviral infection of gRNA/Cas9^WT^ (left) or gRNA/Cas9^D10A^ (right). (n=5 replicates)

BIR is engaged to repair seDSBs occurring on broken forks ^13,14^. We previously showed that at nick-induced seDSBs, BIR/LTGC is used more frequently and both BIR/LTGC and HR/STGC rely on PIF1 and POLD3, which differs from the repair at deDSBs generated by endonucleases, where PIF1- and POLD3-independent HR/STGC is predominantly utilized over BIR/LTGC at deDSBs ^19^. We postulated that BIR replisomes containing PIF1 and POLD3 are readily assembled upon fork breakage as BIR is essential for seDSB repair (Supplementary Figure 4a left), whereas since BIR is not necessarily required in most cases for deDSB repair, BIR replisomes are assembled only when BIR is in need (Supplementary Figure 4a right top) ^19^. In our current study, we showed that deficiency in 53BP1 leads to uncontrolled assembly of BIR-like replisomes at deDSBs, driving hyperrecombination using the BIR-like mechanism for both BIR/LTGC and HR/STGC (Supplementary Figure 4a right bottom), analogous to the mechanism utilized at seDSBs on forks (Supplementary Figure 4a left). We propose that 53BP1 plays a regulatory role in restricting BIR at deDSBs, balancing the use of HR and BIR.

### Hyperrecombination due to 53BP1 loss rescues the defect of HR and BIR in BRCA1-deficient cells

BRCA1 is not only required for HR/STGC but also for BIR/LTGC (Fig. 1d) ^19^. It is well established that 53BP1 loss alleviates the reliance on BRCA1 for end resection and HR/STGC ^33,52^. We asked whether loss of 53BP1 would also rescue the BIR defect in BRCA1-deficient cells. Using CRISPR, we generated homozygous deletions of the two BRCT domains (BRCA1-ΔBRCT) at the C-terminus of BRCA1 after the coiled coil domain that binds to PALB2 [Supplementary Figure 3b ^56–58^] in U2OS (EGFP-BIR/LTGC) and U2OS (EGFP-HR/STGC) cell lines. Both BIR and HR are defective in BRCA1-ΔBRCT cells, which can be restored after inactivating 53BP1 (Fig. 3b). Like that in 53BP1-deficient cells (Fig. 3a right, 3c left four lanes and S3c), restored HR/STGC at deDSBs upon 53BP1 depletion in BRCA1-ΔBRCT cells after I-SceI cleavage is also reliant on PIF1 and POLD3 (Fig. 3c right four lanes and S3c). This is consistent with our model that loss of 53BP1 leads to rewiring of the recombination machinery using the BIR-like mechanism, involving a role associated with PIF1 and POLD3, to promote hyper HR and BIR at deDSBs (Supplementary Figure 4a right bottom), and this gained hyperrecombination activity rescues the HR/BIR defect in BRCA1-deficient cells.

To monitor BIR on broken replication forks, we used Cas9 nickase, Cas9^D10A^, to generate nicks on the EGFP-BIR/LTGC reporter, where seDSBS are produced when replication encounters the nicks (Supplementary Figure 4b). Like at deDSBs generated by Cas9^WT^, BIR is also substantially increased in *53BP1*-KO cells at seDSBs induced by Cas9^D10A^ (Fig. 3d). We also examined BIR on the replication forks induced at the structure-prone Flex1 sequence derived from common fragile sites (CFSs) upon replication stress using the EGFP-BIR/LTGC-Flex1 reporter ^19^. We showed that BIR induced at Flex1 by FANCM depletion or HU treatment, which causes fork stalling and fork breakage at Flex1 due to DNA secondary structures ^59^, is substantially increased when 53BP1 is depleted (Fig. 3e and S3d). These data suggest that 53BP1 also limits the BIR activity at seDSBs on broken forks. We further showed that BIR is defective at seDSBs generated by Cas9^D10A^ in BRCA1-ΔBRCT cells, and 53BP1 inactivation rescues the BIR defect in BRCA1-ΔBRCT cells not only at deDSBs after Cas9^WT^ cleavage, but also at seDSBs on broken forks after Cas9^D10A^ cleavage (Fig. 3f). We conclude that BIR defect at both seDSBs and deDSBs in BRCA1-deficient cells can be rescued by the loss of 53BP1.

### Loss of 53BP1 leads to accumulation of PIF1 at DSB ends after IR in a manner dependent on PCNA and Polα-primase

To investigate how BIR-like activity is stimulated in *53BP1*-deficient cells resulting in hyperrecombination, we monitored PIF1 recruitment to DSB ends upon laser microirradiation. The longer in *53BP1*-KO cells (Fig. 4a and S5a). We also performed an *in situ* proximity ligation assay (PLA) of stably expressed Flag-PIF1 and γH2AX after IR (4 Gy). The expression of Flag-PIF1, assayed by qPCR, is estimated to be ∼1.8 fold as that of endogenous PIF1 (S5b). While only a limited amount of PIF1 is present at DSB ends marked by γH2AX in WT cells after IR, a significant recruitment of PIF1 occurs at IR-induced DSBs in *53BP1*-KO cells (Fig. 4b and S5c). Similarly, depletion of 53BP1 or RIF1 results in PIF1 accumulation at IR-induced DSBs (Fig. 4c and S5d). In addition, PCNA recruitment to IR-induced DSB ends is also significantly increased in *53BP1*-KO cells (Fig. 4d and S5e) or when 53BP1 is depleted (Supplementary Figure 5f). Immunostaining experiments also showed more colocalization of Flag-PIF1 and PCNA with γH2AX after IR (4 Gy) in *53BP1*-KO cells compared to WT cells (Supplementary Figure 6a and S6b). Furthermore, similar to PIF1 recruitment to stalled replication forks, which relies on PCNA ^19^, we demonstrated that PIF1 recruitment to IR-induced DSB ends in 53BP1-deficient cells is also dependent on PCNA as revealed by PLA (Fig. 4e and S5g). 53BP1 plays an important role in promoting KU-dependent NHEJ ^32,60^. To exclude the possibility that PIF1 and PCNA accumulation at γH2AX sites in 53BP1-deficient cells is caused by an increased DSB formation due to a defect in NHEJ, we performed PLA of PIF1 with γH2AX after depleting KU70 and XRCC4. PLA signals of PIF1 with γH2AX are only increased after depletion of 53BP1, but not KU70 and XRCC4 (Supplementary Figure 7a), suggesting a specific role of 53BP1 in restricting PIF1 accumulation to DSBs after IR. We propose that restricted recruitment of PIF1 to deDSBs in 53BP1-proficient WT cells may prevent the overuse of BIR, while the substantial increase of PIF1 recruitment to IR-induced deDSBs in 53BP1-deficient cells in a manner dependent on PCNA facilitates the establishment of BIR-like hyperrecombination.

**Fig. 4.**
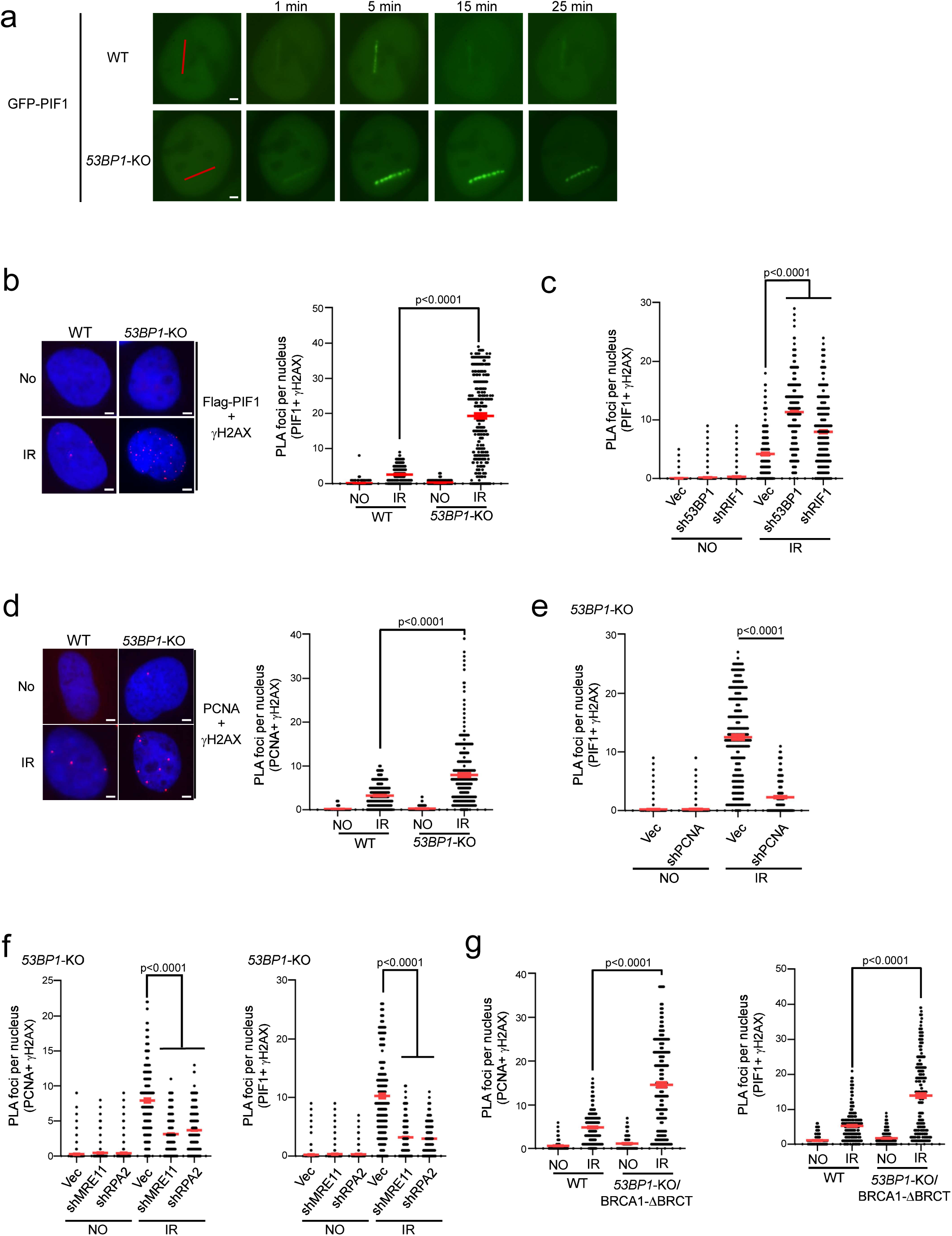
PIF1 is accumulated at DSBs after IR when the 53BP1 pathway is compromised. (a). Time-lapse live cell imaging of GFP-PIF1 in U2OS WT and *53BP1*-KO cells was performed before and after laser-induced microirradiation. Red lines: laser-induced damage region. Scale bar =2 μm. Also see Supplementary Figure 5a. (b and c). Recruitment of Flag-PIF1 to γH2AX sites was analyzed by PLA in U2OS WT and *53BP1*-KO cells (b), or in U2OS cells expressing shRNAs targeting 53BP1 or RIF1 and a vector control (c) before and after IR (4 Gy) treatment. Left: representative PLA images. Right: quantification of PLA foci per nucleus. Also see Supplementary Figure 5c for b and Supplementary Figure 5d for c. Scale bar =2 μm. (n=300 cells) (d). Recruitment of PCNA to γH2AX sites was analyzed by PLA in U2OS WT and *53BP1*-KO cells treated with or without IR (4 Gy). Left: representative PLA images. Right: quantification of PLA foci per nucleus. Also see Supplementary Figure 5e. Scale bar =2 μm. (n=298 cells) (e). Recruitment of PIF1 to γH2AX sites was analyzed by PLA in U2OS cells expressing PCNA shRNA with vector as a control after treatment with IR (4 Gy). Quantification of PLA foci per nucleus is displayed. Also see Supplementary Figure 5g. (n=304 cells) (f). Recruitment of PCNA (left) and PIF1 (right) to γH2AX sites was analyzed by PLA in U2OS *53BP1*-KO cells expressing shRNAs targeting MRE11 or RPA2 with vector as a control after treatment with IR (4 Gy). Quantification of PLA foci per nucleus is displayed. Also see Supplementary Figure 5h. (n=300 cells) (g). Recruitment of PCNA (left) and PIF1 (right) to γH2AX sites was analyzed by PLA in U2OS WT and *53BP1*-KO/BRCA1-ΔBRCT cells with or without treatment of IR (4 Gy). Three experiments were performed with ∼100 nuclei analyzed in each experiment. Quantification of PLA foci per nucleus is displayed. Also see Supplementary Figure 5i. (n=300 cells) b to g: Three experiments were performed with ∼100 nuclei analyzed in each experiment. Quantification of PLA foci per nucleus from a total of ∼300 nuclei are displayed.

Since the loss of 53BP1 yields hyper-resection ^33^, we asked whether elongated ssDNA overhangs play a role in PCNA and PIF1 recruitment to IR-induced DSBs. Inactivation of MRE11 and RPA2 significantly reduces the PLA signals of PCNA and PIF1 with γH2AX after IR in *53BP1*-KO cells (Fig. 4f and S5h), suggesting that generating ssDNA overhangs by end resection and subsequent RPA binding are needed for PCNA and PIF1 recruitment to DSBs. However, we observed that the recruitment of PCNA and PIF1 to IR-induced DSBs in *53BP1*-KO/BRCA1-ΔBRCT cells is also significantly more compared to WT cells (Fig. 4g and S5i). Given that 53BP1 loss rescues the end resection defect in BRCA1-deficient cells to the extent comparable to WT cells ^61^ (Supplementary Figure 5j), this data suggests that while generating ssDNA overhangs is necessary, their length is not the key determinant in triggering the overloading of PCNA and PIF1 to IR-induced DSBs in the absence of 53BP1.

Since PCNA recruitment to DSBs after IR is increased in 53BP1-deficient cells, we asked whether PCNA may be ubiquitinated at DSB ends when 53BP1 is compromised. We performed PLA of PCNA with γH2AX using the PCNA Ub antibody that specifically recognizes PCNA ubiquitination at K164 ^62^. While ubiquitinated PCNA (PCNA-Ub) is readily localized to γH2AX sites in WT cells after HU treatment, with a slight increase in *53BP1*-KO cells, we did not detect above-background PLA signals of PCNA-Ub with γH2AX in WT cells after IR, but the PLA signals are strongly induced by IR in *53BP1*-KO cells (Fig. 5a and S8a). This suggests that in contrast to HU, PCNA ubiquitination is triggered at deDSBs by IR only when 53BP1 is deficient. We speculate that certain types of DNA replication/synthesis occur but are stalled at DSB ends in 53BP1-deficient cells, thereby inducing PCNA ubiquitination.

**Fig. 5.**
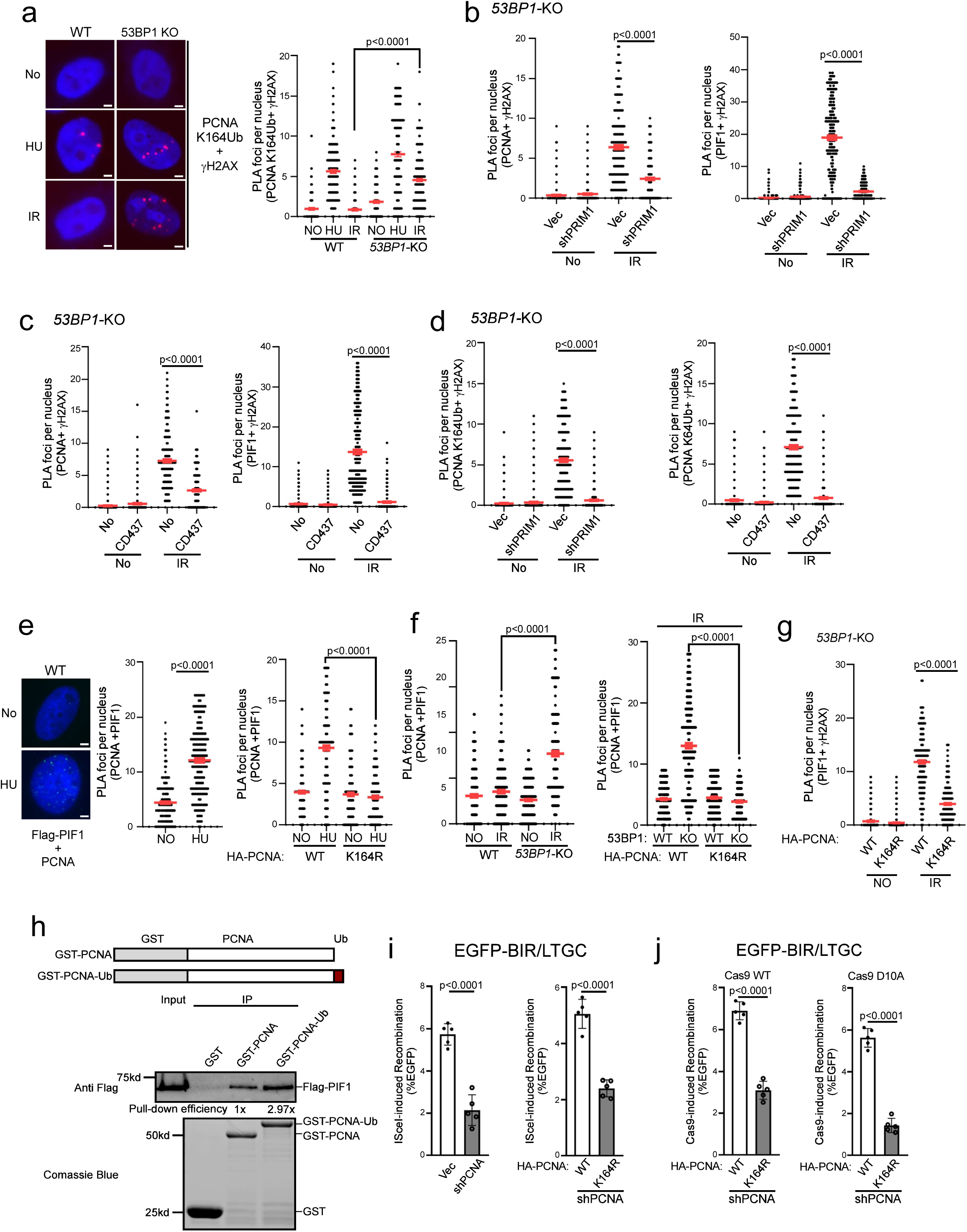
Polα-primase activity is important for PCNA ubiquitination and PIF1 loading to deDSBs after IR to promote BIR. (a). PCNA ubiquitination (K164) at γH2AX sites was analyzed by PLA in U2OS WT and *53BP1*-KO cells before or after treatment with HU (1 mM, 24h) or IR (4 Gy, 2h after for PLA). Left: representative PLA images. Right: quantification of PLA foci per nucleus. Also see Supplementary Figure 8a. Scale bar =2 μm. (n=300 cells) (b). Recruitment of PCNA (left) and PIF1 (right) to γH2AX sites was analyzed by PLA in U2OS *53BP1*-KO cells expressing shRNA targeting PRIM1 upon IR (4 Gy, 2h after for PLA). Quantification of PLA foci per nucleus is displayed. Also see Supplementary Figure 8b. (n=310 cells) (c). Recruitment of PCNA (left) and PIF1 (right) to γH2AX sites was analyzed by PLA in U2OS *53BP1*-KO cells in the presence of Polα inhibitor CD437 (10 μM) upon IR (4 Gy, 2h after for PLA). Quantification of PLA foci per nucleus is displayed. Also see Supplementary Figure 8c. (n=300 cells) (d). PCNA ubiquitination (K164) at γH2AX sites was analyzed by PLA in U2OS *53BP1*-KO cells expressing shRNAs targeting PRIM1 (left) or in the presence of Polα inhibitor CD437 (10 μM, right) upon IR (4 Gy, 2h after for PLA). Quantification of PLA foci per nucleus is displayed. Also see Supplementary Figure 8d. (n=306 cells) (e). PLA was performed to assay for the interactions of stably expressing Flag-PIF1 with endogenous PCNA in U2OS cells (left and middle) or with expressed HA-PCNA-WT and HA-PCNA-K164R (right) with or without HU treatment (1 mM, 24h). Left: representative PLA images. Right: quantification of PLA foci per nucleus. Also see Supplementary Figure 8e. Scale bar =2 μm. (n=300 cells) (f). PLA was performed to assay for the interactions of stably expressing Flag-PIF1 with endogenous PCNA in U2OS WT and *53BP1*-KO cells (left) or with expressed HA-PCNA-WT and HA-PCNA-K164R (right) with or without IR (4 Gy, 2h after for PLA). Quantification of PLA foci per nucleus is displayed. Also see Supplementary Figure 8f. (n=300 cells) (g). Recruitment of PIF1 to γH2AX sites was analyzed by PLA in U2OS cells expressing HA-PCNA-WT or K164R with endogenous PCNA depleted by shRNA upon IR (4 Gy, 2h after for PLA). Quantification of PLA foci per nucleus is displayed. Also see Supplementary Figure 8g. (n=296 cells) a to g: Three experiments were performed with ∼100 nuclei analyzed in each experiment. Quantification of PLA foci per nucleus from a total of ∼300 nuclei are displayed. (h). A schematic drawing depicts GST-PCNA and GST-PCNA-Ub (top). Pull-down experiments were performed using GST-PCNA, GST-PCNA-Ub or GST and Flag-PIF1 expressing in 293T cells. Anti-Flag Western blotting and Coomassie blue staining for GST proteins are shown (bottom). The relative fold of Flag-PIF1 Western signals over GST-PCNA or GST-PCNA-Ub is indicated as pull-down efficiency. (i and j). BIR frequency was determined in U2OS (EGFP-BIR/LTGC) cells expressing PCNA shRNA with vector as a control 5 days after I-Sce1 expression (i, left), or in cells expressing HA-PCNA-WT or HA-PCNA-K164R with endogenous PCNA depleted by shRNA, 5 days after I-Sce1 expression (i, right) or Cas9^WT^ (j, left) and Cas9^D10A^ (j, right) expression. (n=5 replicates)

It has been shown that DNA synthesis primed by Polα-primase occurs at ssDNA overhangs of deDSBs to counteract end resection ^47,48^. Although shieldin-dependent loading of the CST (CTC1, STN1 and TEN1) complex contributes to promoting Polα-primase recruitment ^47,48^, Polα activity for end fill-in DNA synthesis on the ssDNA overhangs is sustained in 53BP1-deficient cells ^63^. We hypothesize that when the 53BP1 pathway is compromised, end fill-in DNA synthesis on ssDNA overhangs, directed by shieldin-independent Polα-primase activity, is often stalled, leading to PCNA ubiquitination. We depleted PRIM1, a subunit of primase by shRNA or inhibited Polα activity by its specific inhibitor CD437 ^64^, and performed PLA of PCNA and PIF1 with γH2AX after IR in *53BP1*-KO cells. We found that the elevated recruitment of PCNA and PIF1 to the γH2AX sites in *53BP1*-KO cells after IR, as well as PCNA ubiquitination at γH2AX sites, is strongly dependent on PRIM1 (Fig. 5b, 5d left, S8b and S8d left) and on Polα activity (Fig. 5c, 5d right, S8c and S8d right). In addition to end fill-in DNA synthesis on ssDNA overhangs at deDSBs, it remains possible that PCNA ubiquitination is induced during BIR D-loop migration DNA synthesis. Since BIR DNA synthesis occurs after RAD51-medaited strand invasion, we performed PLA of PCNA-Ub with γH2AX upon IR in 53BP1 deficient cells after depleting RAD51. While inhibiting primase activity by the inhibitor CD437 strongly inhibits PCNA-Ub accumulation at DSBs, RAD51 depletion fails to do so (Supplementary Figure 7b), suggesting that the signals triggering PCNA ubiquitination at DSBs are generated prior to strand invasion and D-loop formation during BIR. These data support the model that when the 53BP1 pathway is deficient, Polα-primase-directed end fill-in DNA synthesis is stalled on ssDNA overhangs, which leads to PCNA overloading and ubiquitination, subsequently facilitating PIF1 recruitment to deDSBs to promote hyperrecombination using the BIR mechanism (Supplementary Figure 13b).

### PCNA ubiquitination enhances the interaction between PCNA and PIF1

It has been shown that yeast Pif1 interacts with PCNA ^65^. By performing PLA of stably expressed Flag-PIF1 with endogenous PCNA, we found that human PIF1 interacts with PCNA in U2OS cells without treatment, and this interaction is enhanced upon HU treatment (Fig. 5e left and middle, and Supplementary Figure 8e left). We also performed co-immunoprecipitation (co-IP), and showed that PIF1 interacts with PCNA, and this interaction is increased after HU treatment (Supplementary Figure 7c). Mutating the PCNA ubiquitination site K164 significantly reduces the PLA signals of PIF1 and PCNA after HU (Fig. 5e right and Supplementary Figure 8e right), suggesting that HU-induced PCNA ubiquitination stimulates the interaction of PIF1 and PCNA. In contrast to HU treatment, we only detected a very minor increase of PIF1 and PCNA interaction by PLA in WT cells after IR, whereas this interaction is significantly induced by IR in *53BP1*-KO cells (Fig. 5f left and Supplementary Figure 8f left). The elevated interaction of PIF1 and PCNA after IR due to 53BP1 loss is abolished in the PCNA-K164R mutant (Fig. 5f right and Supplementary Figure 8f middle). By PLA, we further showed that the recruitment of PIF1 to IR-induced γH2AX sites in 53BP1-deficient cells is dependent on K164 of PCNA (Fig. 5g and Supplementary Figure 8g). Thus, not only the increased PCNA and PIF1 interaction but also PIF1 recruitment to deDSBs in 53BP1-deficient cells after IR is reliant on PCNA ubiquitination at K164.

To further investigate the interaction of PIF1 with PCNA, we purified GST-PCNA and GST-PCNA-Ub, with Ub fused at the PCNA C-terminus to mimic K164 ubiquitination ^66^. While GST-PCNA can readily pull-down Flag-PIF1 expressed in U2OS cells, GST-PCNA-Ub exhibits a stronger interaction with PIF1 (Fig. 5h), supporting the conclusion that human PIF1 interacts with PCNA, and PCNA ubiquitination further enhances their interaction.

BIR is strongly dependent on PCNA [^19^, Fig. 5i left]. To examine whether PCNA ubiquitination at K164 is important for BIR, we expressed Flag-PCNA-WT and Flag-PCNA-K164R in U2OS (EGFP-BIR/LTGC) reporter cell line with endogenous PCNA depleted by shRNA. We showed that BIR after I-SceI cleavage is defective in cells expressing the PCNA-K164R mutant compared to the Flag-PCNA-WT allele with endogenous PCNA depleted by shRNA (Fig. 5i right). We also demonstrated that both Cas9^WT^- and Cas9^D10A^-induced BIR is impaired in the PCNA-K164R mutant cells (Fig. 5j), suggesting that PCNA ubiquitination at K164 is important for BIR at both deDSBs and seDSBs.

### SMARCAD1 displaces 53BP1 at broken forks to antagonize the role of 53BP1 in BIR suppression

53BP1 is enriched on stalled replication forks and plays an important role in protecting nascent DNA on stalled forks ^67–70^. We showed that in addition to elevated BIR at deDSBs, 53BP1 loss also results in increased BIR at broken forks induced by Cas9^D10A^ nicking and Flex1 (Fig. 3d and 3e), suggesting that 53BP1 also has a role in suppressing BIR at seDSBs on broken forks. However, since BIR is preferentially established on broken forks to repair seDSBs in 53BP1-proficient WT cells ^19^, the mechanism of 53BP1 in BIR suppression on broken forks must be different from that on deDSBs, where BIR is restricted in WT cells.

As described above, upon HU treatment, PCNA ubiquitination is readily induced in 53BP1-proficient WT cells, in sharp contrast to minimal PCNA ubiquitination detected at deDSBs after IR unless the 53BP1 pathway is compromised (Fig. 5a). Upon replication stress, forks often stall prior to breakage, which could result in PCNA ubiquitination. We anticipate that at deDSBs, a mechanism is required to induce PCNA ubiquitination (such as inactivating the 53BP1 pathway), serving as a prerequisite step for BIR activation, while this step, however, is already accomplished on stalled forks prior to breakage (See Discussion).

It has been shown that 53BP1 plays a role in antagonizing PCNA loading at replication restart sites, and SMARCAD1 acts to displace 53BP1 to allow sufficient PCNA loading ^71^. Interestingly, while we observed similar levels of RAD51 recruitment to DSBs after releasing cells from HU (1 mM, 24 h), the condition inducing fork breakage (Supplementary Figure 9g), and IR treatment (Fig. 6a left and middle, and Supplementary Figure 9a left), 53BP1 localization to γH2AX sites is significantly lower at seDSBs after HU compared to that at deDSBs after IR (Fig. 6a left and right, and Supplementary Figure 9a right). This observation implies that 53BP1 may be actively displaced from DSBs at broken forks preparing for replication restart. In addition, depleting SMARCAD1 substantially increases 53BP1 binding to γH2AX sites after releasing from HU, accompanied with a reduction of PCNA and PIF1 loading to γH2AX sites (Fig. 6b and Supplementary Figure 9b). This suggests that SMARCAD1 plays an active role in removing 53BP1 from DSBs on broken forks, enabling sufficient loading of PCNA and PIF1 for replication restart. Moreover, consistent with a role of 53BP1 in constraining PCNA loading, we observed an increased PLA signals of PCNA and PIF1 with γH2AX in *53BP1*-KO cells after releasing from HU (Fig. 6c and Supplementary Figure 9c). Collectively, we propose that at seDSBs on broken forks, 53BP1 does not directly influence PCNA ubiquitination but rather exerts an activity to inhibit PCNA loading onto DSB ends. Conversely, SMACAD1 plays a role in displacing 53BP1 from DSBs on broken forks, thereby introducing a regulatory mechanism for BIR activation to facilitate fork restart via modulating PCNA and PIF1 loading.

**Fig. 6.**
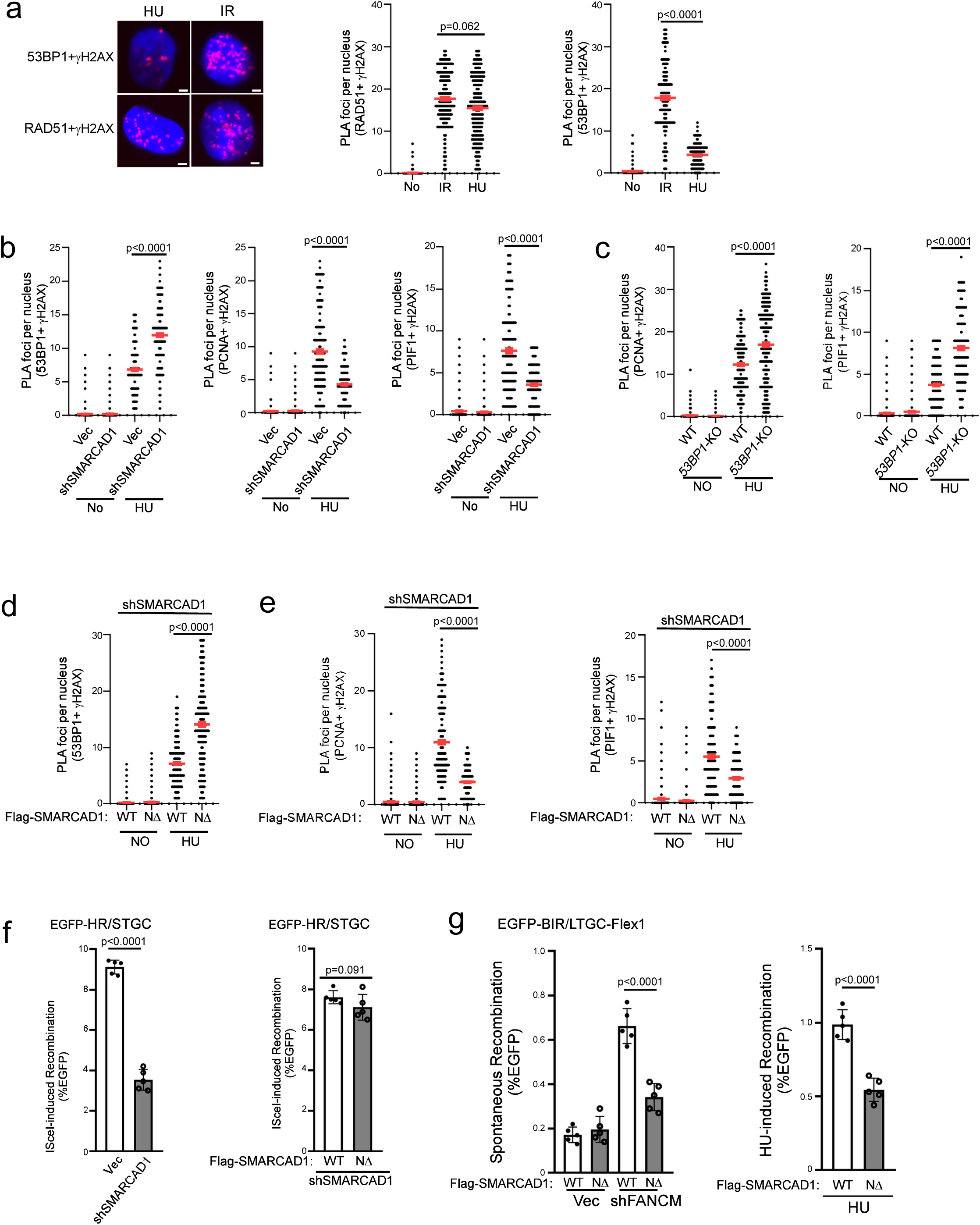
SMARCAD1 displaces 53BP1 at γH2AX sites on broken forks to promote BIR. (a). Recruitment of RAD51 (left and middle) and 53BP1 (left and right) to γH2AX sites was analyzed by PLA in U2OS cells treated with IR (4 Gy, 2h after for PLA) or HU (1 mM, 24h, 30 min after release for PLA). Left: representative PLA images. Right: quantification of PLA foci per nucleus. Also see Supplementary Figure 9a. Scale bar =2 μm. (n=300 cells) (b). Recruitment of 53BP1 (left), PCNA (middle) and PIF1 (right) to γH2AX sites was analyzed by PLA in U2OS cells expressing SMARCAD1 shRNAs with vector as a control after HU treatment (1 mM, 24h). Quantification of PLA foci per nucleus is displayed. Also see Supplementary Figure 9b. (n=300 cells) (c). Recruitment of PCNA (left) and PIF1 (right) to γH2AX sites was analyzed by PLA in U2OS WT and *53BP1*-KO cells after HU treatment (1 mM, 24h). Quantification of PLA foci per nucleus is displayed. Also see Supplementary Figure 9c. (n=300 cells) (d and e). Recruitment of 53BP1 (d), PCNA (e, left) and PIF1 (e, right) to γH2AX sites was analyzed by PLA in U2OS cells expressing Flag-SMACRAD1-WT or Flag-NΔ-SMACRAD1 with endogenous SMACRAD1 depleted by shRNA after HU treatment (1 mM, 24h). Quantification of PLA foci per nucleus is displayed. Also see Supplementary Figure 9d for d and Supplementary Figure 9e for e. (n=300 cells) a to e: Three experiments were performed with ∼100 nuclei analyzed in each experiment. Quantification of PLA foci per nucleus from a total of ∼300 nuclei are displayed. (f). HR frequency was determined in U2OS (EGFP-HR/STGC) cells expressing SMACRAD1 shRNA with a vector control (left) or in cells expressing Flag-SMACRAD1-WT or Flag-NΔ-SMACRAD1 with endogenous SMACRAD1 depleted by shRNAs, 5 days after I-Sce1 lentiviral infection. Also see Supplementary Figure 9d. (n=5 replicates) (g). U2OS (EGFP-BIR/LTGC-Flex1) cells expressing Flag-SMACRAD1-WT or Flag-NΔ-SMACRAD1 with endogenous SMACRAD1 depleted by shRNA were further infected with lentiviruses encoding FANCM shRNA (left) or synchronized to S-phase using double thymidine block followed by HU treatment (1 mM, 24h, right). BIR frequency was assessed by FACS 6 days after. Also see S9f. (n=5 replicates)

At deDSBs, SMARCAD1 has been shown to act downstream of BRCA1, competing with 53BP1 to promote end resection and HR ^72–74^. To distinguish the SMARCAD1 activity in displacing 53BP1 to facilitate replication restart and its role in HR at deDSBs, we used a separation-of-function mutant NΔ-SMARCAD1 that is defective in PCNA binding and 53BP1 displacement but proficient in HR to repair deDSBs generated by endonucleases ^71^. After depleting endogenous SMARCAD1 by shRNAs, the PLA signals of 53BP1 with γH2AX after HU treatment in NΔ-SMARCAD1 mutant cells is significantly increased compared to SMARCAD1-WT cells (Fig. 6d and Supplementary Figure 9d), confirming a defect of this mutant in 53BP1 displacement at DSBs on broken forks. Significantly, PCNA and PIF1 loading at DSBs on broken forks after HU treatment is much reduced in NΔ-SMARCAD1 mutant cells than that in SMARCAD1-WT cells (Fig. 6e and Supplementary Figure 9e). This suggests that SMARCAD1-mediated 53BP1 displacement is important for PCNA and PIF1 loading to DSB ends on broken forks.

Furthermore, while depleting SMARCAD1 leads to a reduction of HR (Fig. 6f left and Supplementary Figure 9b right), expressing NΔ-SMARCAD1 mutant with endogenous SMARCAD1 depleted does not results in HR defect at deDSBs after I-SceI cleavage compared to cells expressing SMARCAD1-WT allele (Fig. 6f right and Supplementary Figure 9d right), in agreement with the previous findings ^71^. However, Flex1-induced BIR on broken forks, either after FANCM depletion or HU treatment, is deficient in U2OS (EGFP-BIR/LTGC-Flex1) cells expressing the NΔ-SMARCAD1 mutant but not the SMARCAD1-WT allele (Fig. 6g and Supplementary Figure 9f). Thus, the activity of SMARCAD1 in fork-specific 53BP1 displacement is important for promoting BIR upon fork breakage.

### Cells deficient in the 53BP1 pathway rely more on PIF1 for survival

While 53BP1 loss impairs NHEJ ^32,60^, our study demonstrated that hyperrecombination resulting from 53BP1 deficiency is PIF1-and POLD3-dependent. Therefore, cells deficient in the 53BP1 pathway may rely more on BIR for DSB repair to survive. We depleted PIF1 and POLD3 by shRNAs in U2OS WT and *53BP1*-KO cells and observed a significant increase in cell death of *53BP1*-KO cells (Fig. 7a, Supplementary Figure 10a and Supplementary Figure 10b). We also expressed shRNAs for 53BP1 and RIF1 in U2OS WT and *PIF1*-KO cells (Fig. 7b and Supplementary Figure 10c), 53BP1 shRNA in RPE WT and *PIF1*-KO cells (Supplementary Figure 10d) and POLD3 shRNA in RPE WT and *53BP1*-KO cells (Supplementary Figure 10e). We found that combined inactivation of 53BP1 or RIF1 with PIF1 or POLD3 significantly reduces cell viability, suggesting a synthetic interaction of the BIR pathway with the

**Fig. 7.**
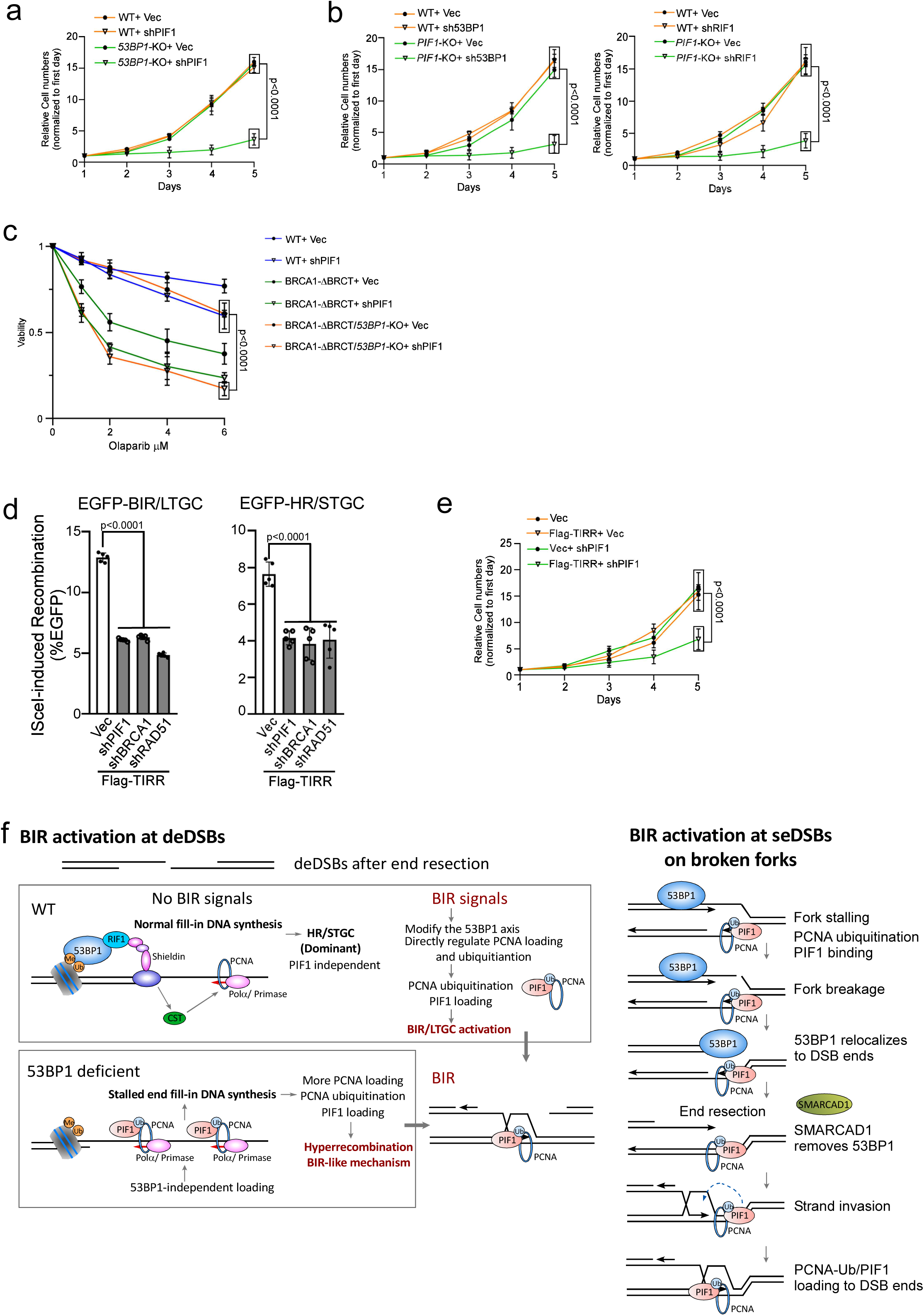
Inactivation of BIR by targeting PIF1 causes cell death when the 53BP1 pathway is compromised. (a). The growth curves of U2OS WT and *53BP1*-KO cells were plotted after infection with lentiviruses expressing PIF1 shRNA with vector as a control. The expression of PIF1 was examined by qPCR (Supplementary Figure 10a). (n=3 replicates) (b). The growth curves of U2OS WT and *PIF1*-KO cells were plotted after infection with lentiviruses expressing shRNAs targeting 53BP1 (left) or RIF1 (right) with vector as a control. The expression of 53BP1 and RIF1 was examined by qPCR (Supplementary Figure 10c). (n=3 replicates) (c). Cell viability was determined in U2OS WT, BRCA1-ΔBRCT and *53BP1*-KO/BRCA1-ΔBRCT cells expressing PIF1 shRNA with a vector control after treatment with the indicated concentrations of Olaparib for 72 hours. The expression of PIF1 was determined by qPCR (Supplementary Figure 11a). (n=3 replicates) (d) U2OS (EGFP-BIR/LTGC) cells (left) and U2OS (EGFP-HR/STGC) cells (right) overexpressing Flag-TIRR were infected with shRNAs targeting PIF1, BRCA1 or RAD51 with vector as a control, followed by infection with lentiviruses encoding I-SceI. The percentage of EGFP-positive cells was determined by FACS, 5 days post-infection. The expression of Flag-TIRR was examined by Western blot analysis (Supplementary Figure 12a). The expression of PIF1, BRCA1 and RAD51 was determined by qPCR (Supplementary Figure 12b). (n=5 replicates) (e). The growth curves of U2OS cells with or without overexpressing Flag-TIRR were plotted after infection with lentiviruses expressing PIF1 shRNA with vector as a control. The expression of PIF1 was determined by qPCR (Supplementary Figure 12c). (n=3 replicates) (f). Working models depicting the involvement of 53BP1 in limiting BIR at deDSBs (left) and at seDSBs on broken forks (right). See details in the main context.

Deficiency in the 53BP1 pathway renders BRCA1-deficient cells resistant to PARPi ^33^. We showed that inhibition of PIF1 sensitizes *53BP1*-KO/BRCA1-ΔBRCT cells to Olaparib treatment (Fig. 7c and Supplementary Figure 11a), supporting the notion that the onset of BIR-like hyperrecombination resulting from the loss of 53BP1 rescues the HR/BIR defect in BRCA1-deficient cells. We also used BRCA1-deficient ovarian cancer cell line UWB1 ^75,76^ and showed that acquired Olaparib resistance of UWB1 cells upon 53BP1 depletion can be reverted by inhibiting PIF1 (Supplementary Figure 11b).

TIRR, which interacts with 53BP1, inhibits the binding of the 53BP1 Tudor domain to H4K20me2 on chromatin ^77–80^. Overexpression of TIRR abolishes 53BP1 foci formation and confers PARPi resistance of BRCA1-deficient cells ^77^. We showed that similar to 53BP1 deficiency, TIRR overexpression induces hyperrecombination for both HR and BIR, exhibiting dependence on PIF1 (Fig. 7d, Supplementary Figure 12a and Supplementary Figure 12b). In addition, depleting PIF1 leads to more cell death in TIRR-overexpressing cells compared to normal cells (Fig. 7e and Supplementary Figure 12c).

## Discussion

### 53BP1 suppresses BIR-like hyperrecombination at deDSBs

DSBs can be repaired by multiple pathways, and end resection has been thought to be a critical determinant in repair pathway selection ^81^. It is well established that the role of 53BP1 in end protection suppresses end resection and promotes the selection of NHEJ over HR ^30–32^. Here we identified another role of 53BP1 in suppressing BIR-like hyperrecombination at deDSBs after end resection to favor the use of HR over BIR (Supplementary Figure 13a).

At deDSBs, BIR is restricted and HR is used predominately ^19^. Using the reporter systems, we showed that loss of the 53BP1 control not only leads to hyperrecombination at deDSBs, but also triggers a change in the recombination mechanism, involving the participation of PIF1 and POLD3 for both HR/STGC and BIR/LTGC at deDSBs (Supplementary Figure 4a right bottom), mirroring the mechanism utilized at broken forks, where the BIR mechanism is established for both HR/STGC and BIR/LTGC (Supplementary Figure 4a left) ^19^. We speculate that promoting the assembly of BIR-like replisomes is probably the mechanism leading to hyperrecombination at deDSBs in cells deficient for the 53BP1 pathway. BIR is mutagenic, causing a high mutation rate and template switching-mediated chromosomal rearrangements ^25–27^. Therefore, it is important to restrict BIR activity at deDSBs unless its utilization is necessary (Supplementary Figure 4a right top and S13a bottom). We propose that 53BP1 is involved in the regulation of BIR at deDSBs by preventing the establishment of the BIR mechanism, thereby allowing predominant utilization of HR/STGC at deDSBs (Supplementary Figure 4a right).

Loss of 53BP1 rescues the HR defect in BRCA1-deficient cells ^33^. The current model is that 53BP1 deficiency enables more extensive end resection, thereby suppressing NHEJ at deDSBs and consequently allowing for more efficient HR. Based on our findings, we propose that restoration of HR activity upon 53BP1 loss in BRCA1-deficient cells is not merely a simple passive acquisition of HR due to compromised NHEJ at over-resected deDSB ends, but also involves an active mechanism to induce BIR-like hyperrecombination. 53BP1 binds to a large chromosomal region around DSBs, spanning thousands of kbs, while the association of BRCA1 is more localized to DSB ends, typically within 1-2 kbs ^82,83^, and thus the competition between BRCA1 and 53BP1 for chromatin loading is likely restricted to the DSB proximal sites. In normal 53BP1-proficient cells, BRCA1 competes away 53BP1 at DSB ends to facilitate extensive end resection and HR (Supplementary Figure 13a top). We propose that after end resection when HDR is already committed, 53BP1 chromatin binding internal to the ssDNA overhangs has another role in suppression of the use of BIR, thereby ensuring efficient onset of HR (Supplementary Figure 13a bottom). This aligns with the ssDNA binding activity of the shieldin complex ^43,44,84^, arguing that the 53BP1-shieldin axis has roles after end resection. Taken together, we propose that while 53BP1 antagonizes the extensive end resection to promote the selection of NHEJ over HR, once end resection is achieved, 53BP1 suppresses BIR activation to facilitate the use of HR at deDSBs. Balancing the choice of HR versus BIR at deDSBs is important for achieving not only sufficient repair but also DSB repair with high fidelity.

### The mechanism underlying the activation of BIR-like hyperrecombination at deDSBs upon loss of the 53BP1 control

Significant PCNA ubiquitination and subsequent PIF1 recruitment to deDSBs after IR are observed only in 53BP1-deficient cells, not in 53BP1-proficient cells (Fig. 5a and Fig. 4b). Considering the requirement of PCNA ubiquitination at K164 for PIF1 recruitment and BIR (Fig. 5g and 5i), we propose that inducing PCNA ubiquitination and PIF1 recruitment represents a critical step in establishing the BIR mechanism at deDSBs. Given that PCNA ubiquitination is typically instigated by replication stalling ^85^, we speculate that PCNA ubiquitination may be triggered by the stalling of end fill-in DNA synthesis on ssDNA overhangs at deDSBs. Indeed, IR-induced PCNA ubiquitination in 53BP1-deficient cells is dependent on Polα-primase activity (Fig. 5b, 5c and 5d), which is required for initiating the end fill-in DNA synthesis at deDSBs. We propose that 53BP1 deficiency disrupts the balance of end fill-in DNA synthesis on ssDNA overhangs by interfering with the coordination of Polα-primase-directed priming and the subsequent elongation of DNA synthesis (Fig. 7f, left and Supplementary Figure 13b). This disruption induces fork stalling on ssDNA overhangs, subsequently leading to increased PCNA loading, PCNA ubiquitination and PIF1 loading at the vicinity of DSBs. We anticipate that following strand invasion, PIF1, in association with PCNA-Ub enriched around DSBs, could be recruited promptly to the 3’ of the invading strands to support Polδ for BIR DNA synthesis and to facilitate D-loop migration, thereby establishing the BIR mechanism for recombination repair (Supplementary Figure 13b).

Localized DNA synthesis directed by Polα-primase on ssDNA overhangs at DSBs has been detected genome-wide ^63^. 53BP1 downstream effector CST has been shown to promote Polα-primase loading to ssDNA overhangs to counteract end resection ^47,48^, yet this regulation mainly occurs in G0/G1 cells ^63^. Moreover, in the absence of 53BP1, Polα activity on ssDNA overhangs is sustained for local DNA synthesis, suggesting that Polα-primase can be recruited to ssDNA overhangs independently of CST ^63^. We speculate that Polα-primase loading to ssDNA overhangs in 53BP1 deficient cells could be facilitated by the direct interactions of Polα-primase with RPA ^86,87^.

We anticipate that the 53BP1 axis establishes a context, in which the loading of Polα-primase onto ssDNA overhangs is coordinated with efficient DNA synthesis from the priming sites, ensuring smooth DNA synthesis without stalling (Fig. 7f, left top). When the 53BP1 pathway is impaired, 53BP1/CST-independent loading of Polα-primase cannot be sufficiently coupled with DNA synthesis, resulting in fork stalling and PCNA ubiquitination, which in turn leads to PIF1 loading to trigger the establishment of BIR-like mechanism for repairing deDSBs. In this respect, CST has been shown to interact with the replication accessary protein AND1, which is required for optimal replication fork progression ^88,89^. In the 53BP1 network, CST-mediated loading of Polα-primase may be coupled with CST replication activity to support efficient DNA synthesis at established priming sites without eliciting DNA synthesis stalling. Additionally, RIF1 is linked to DNA replication by controlling replication timing ^90,91^, and it remains possible that RIF1 has a role in supporting efficient DNA synthesis from Polα-primase priming sites on ssDNA overhangs.

### A working model for BIR activation at deDSBs

Despite most deDSBs are channeled to HR/STGC for repair, BIR is still utilized at deDSBs in normal cells, albeit in a restrained manner. Based on the observation that BIR-like hyperrecombination is established in 53BP1-deficient cells, we propose that in normal cells, a regulatory mechanism under the control of the 53BP1 axis is employed to suppress BIR at deDSBs unless its use is necessary (Fig. 7f, left top). The activation of BIR at deDSBs requires BIR signals, which currently are still elusive but possibly stem from certain types of DNA ends, cell cycle status or stressed cell conditions. The BIR signals may induce the displacement of 53BP1 or its effectors from deDSBs, or modify the function of these proteins, resulting in PCNA ubiquitination and PIF1 recruitment. Alternatively, the BIR signals may directly target the pathways that modulate PCNA ubiquitination to induce PCNA-Ub accumulation and subsequent PIF1 recruitment, leading to BIR activation at deDSBs.

PIF1 recruitment to deDSBs by PCNA-Ub is likely a key step to activate BIR or BIR-like mechanisms at deDSBs. We showed that PCNA interacts with PIF1 in undamaged cells and this interaction is enhanced upon PCNA ubiquitination. PIF1 family helicases are involved in lagging-strand synthesis and promoting replication through hard-to-replicate sites such as G-quadruplex DNA ^92,93^. The constitutive interaction of PIF1 with PCNA could be important for PIF1 recruitment to forks during normal replication and/or when replication is transiently paused at hard-to-replicate sites. PCNA ubiquitination, on the other hand, is often induced upon fork stalling, which could recruit DNA translesion polymerases for damage bypass repair ^85^. In our study, based on the findings from 53BP1-deficient cells, if PCNA ubiquitination is triggered at deDSBs, PCNA-Ub induces BIR onset through PIF1 recruitment, possibly facilitating the assembly of BIR replisomes at deDSBs. Studies in yeast showed that Pif1 facilitates Polδ recruitment to D-loops and possibly is also involved in unwinding the template DNA ahead of the BIR bubble as well as newly synthesized DNA behind the BIR bubbles ^9,23^. The increased interactions between PCNA and PIF1 following PCNA ubiquitination could result from enhanced affinity of PIF1 for PCNA-Ub or from the presence of a bridging protein that binds to PCNA-Ub, indirectly promoting the interaction between PIF1 and PCNA-Ub.

We anticipate that the establishment of BIR-like replisomes, incorporating the activities of PIF1 and POLD3, underlies the BIR activation at deDSBs. However, despite observing hyper BIR in *53BP1*-KO cells, the BIR tract length remains similar to that in 53BP1-proficient cells. This differs from the onset of BIR on broken forks, where not only BIR is preferentially used, but the repair synthesis tract length is also much longer ^19^. Therefore, while assembling BIR-like replisomes at deDSBs, including PIF1 and POLD3, is important for initiating BIR, processive DNA synthesis during BIR may require additional factors to be incorporated into the BIR replisomes, which may only occur on broken forks but not at replication-independent deDSBs. Although BIR can be established upon receiving BIR signals at replication-independent deDSBs, the operating mechanisms may still not be identical to those on broken forks.

### BIR activation at seDSB upon fork breakage and the role of 53BP1 and SMARCAD1

Unlike at deDSBs, BIR is readily promoted on broken forks as the primary mechanism to repair seDSBs, even for STGC events in normal 53BP1-proficient cells ^19^. However, both the onset of BIR on broken forks and at deDSBs share a common dependence on PCNA ubiquitination at K164 and subsequent recruitment of PIF1. Two notable differences may account for the distinct responses of the immediate BIR activation upon fork breakage and limited BIR onset at deDSBs despite both involving PCNA ubiquitination. First, both PCNA and PIF1 bind to replication forks ^85,92^, providing easy access of PCNA and PIF1 to seDSBs generated upon fork breakage. Second, PCNA ubiquitination is often induced under replication stress due to fork stalling prior to fork breakage, which could “license” BIR for repairing seDSBs on broken forks. Upon fork breakage, PCNA-Ub, in association with PIF1, which are already on forks, could quickly relocate to seDSBs, to assemble BIR replisomes for BIR activation and replication restart. However, PCNA-Ub at deDSBs is not induced upon IR unless the 53BP1 pathway is compromised. Hence, BIR activation at replication-independent deDSBs requires a mechanism to induce PCNA ubiquitination (e.g. loss of 53BP1 control), while the signals bearing with PCNA ubiquitination for BIR activation are already present on stalled forks prior to fork breakage (Fig. 7f right).

Although BIR can be readily activated upon fork breakage, it is still suppressed by 53BP1. We showed that BIR induced by nicks or Flex1 on forks is also increased when 53BP1 is deficient (Fig. 3d and 3e). However, the mechanisms by which 53BP1 suppresses BIR at seDSBs on broken forks and at deDSBs after IR do not appear the same but could be interconnected. We propose that at deDSBs, the 53BP1 axis acts to prevent PCNA ubiquitination on ssDNA overhangs unless signals calling for BIR are received to break this barrier to activate BIR. On broken forks, 53BP1 may not have a direct role in modulating PCNA ubiquitination, as it is likely already induced due to replication stress-induced fork stalling, but instead, 53BP1 antagonizes the localization of PCNA/PCNA-Ub from forks to the DSB ends (Fig. 7f right).

It has been shown that 53BP1 is enriched on stalled forks, where it is important for protecting nascent DNA in a cell type-dependent manner ^67–70,94,95^. Upon releasing from replication stress, SMARCAD1 displaces 53BP1 from replication active sites to facilitate replication restart ^71^. We demonstrated that SMARCAD1 is involved in removing 53BP1 from DSB ends on broken forks. At deDSBs, SMARCAD1 functions in the BRCA1 pathway, removing or repositioning 53BP1 on chromatin to promote end resection and HR ^72^. By using the separation-of-function NΔ-SMARCAD1 mutant ^71^, we demonstrated that a fork-specific and HR-independent SMARCAD1 activity in displacing 53BP1 from seDSB ends on broken forks is important for PCNA ubiquitination and PIF enrichment to promote BIR. Consistent with the role of 53BP1 in preventing PCNA loading to seDSBs on broken forks, 53BP1 has been shown to interact with PCNA-unloader ATAD5 to disassociate PCNA from forks and antagonize the role of SMARCAD1 in PCNA loading ^71^. We propose that the function of 53BP1 in preventing PCNA/PCNA-Ub from relocating to seDSBs ends on broken forks is likely the underlying mechanism for the BIR suppression activity of 53BP1 on broken forks.

Discovering the antagonizing roles of 53BP1 and SMARCAD1 in modulating BIR activity on broken forks has highlighted the involvement of regulatory mechanisms for BIR activation upon fork breakage. The presence of PCNA and PIF1 on active forks ^85,92^, along with the enrichment of PCNA-Ub and PIF1 on stalled forks, qualify the potential use of BIR on broken forks, but the launch of BIR and the balance of BIR with other repair mechanisms still require additional regulations. We anticipate that displacing 53BP1 by SMARCAD1 from DSB ends on broken forks is one of the mechanisms in response to fork breakage to modulate BIR activity (Fig. 7f right). At this stage, however, the signals that trigger SMARCAD1 to displace 53BP1 from seDSB ends to facilitate BIR upon fork breakage, and how this process is coordinated with other regulatory mechanisms for BIR activation, remain unclear. Further investigations are necessary to unravel the intricacies of the regulatory network for BIR activation upon fork breakage.

### 53BP1 in suppression of genome instability and implication for targeted cancer therapy

53BP1 protects DSB ends and suppresses extensive end resection, which allows quick and efficient repair of DSBs by NHEJ, particularly in G1 when HR is not available ^30–32^. Such activity is important for preventing genome instability caused by unrepaired DSBs. In this study, we uncovered another role of 53BP1 in suppressing genome instability by preventing hyperrecombination using the BIR mechanism. An elevated BIR in 53BP1-deficient cells would cause increased overall template switching, often with microhomology sequences present at the switching junctions (Fig. 2c). Related to this observation, microhomology-mediated templated insertion exhibiting the features of BIR template switching has been shown to accumulate in various cancers ^96,97^. Furthermore, we found that BIR in 53BP1-deficient cells is often accompanied by large deletions at BIR-repair junctions. Taken together, while hyperrecombination resulting from 53BP1 loss would compensate for the defective NHEJ due to 53BP1 deficiency, hyperrecombination using the BIR-like mechanism could also lead to genome instability. Similarly, although the loss of 53BP1 rescues HR in BRCA1-deficient cells, thereby preventing NHEJ-mediated genome instability ^33^, chromosomal rearrangements resulting from using the BIR mechanism would still be anticipated.

BIR has been proposed as a major mechanism for MiDAS, which is used to complete the duplication of under-replicated genomic regions in mitosis, especially when cells are under replication stress ^16^. MiDAS is dependent on POLD3, PIF1, RAD52, but is independent of RAD51 ^19,98,99^ (Supplementary Figure 14a). With our BIR assay using the EGFP-BIR/LTGC reporter, we demonstrated that in cycling cells, BIR is dependent on POLD3, PIF1 and RAD51, but not RAD52 (Fig. 1d left, Fig. 1e and S14b) ^19^. On the other hand, when BIR assay is performed in mitotic cells, BIR exhibits dependence on RAD52, but not on RAD51 ^19^, similar to the dependence for MiDAS. The different requirement of RAD51 and RAD52 for BIR in interphase cells and in mitotic cells is largely due to suppression of the recruitment of BRCA1, RNF8 and RNF168 to DSBs ^100–102^, resulting in abrogation of RAD51 filament formation and inhibition of RAD51-mediated HR in mitosis ^103–107^. In cycling cells, elevated BIR due to 53BP1 deficiency remains to be RAD51-dependent and RAD52 independent (Fig. 1d right, Fig. 1e and S14b). Consistently, by performing immunostaining, we observed a moderate increase of RAD51 colocalized with γH2AX, but a substantial increase of RAD51 colocalized with PCNA and PIF1 in *53BP1*-KO cells compared to WT cells after IR (4 Gy) (Supplementary Figure 15)

It has been also shown that 53BP1 recruitment to DSB ends is attenuated in mitotic cells, likely due to CDK1 and PLK1-mediated phosphorylation ^101,102^, suggesting that 53BP1 is not engaged in regulating DSB repair in mitosis. Along this line, inactivation of 53BP1 does not lead to a defect in MiDAS^108^. Based on our findings that 53BP1 inhibits BIR at DSBs in interphase cells, we propose that suppression of 53BP1 recruitment to DSB ends in mitotic cells could be an important mechanism to facilitate MiDAS. Therefore, although BIR underlies the mechanism of MiDAS, the genetic requirement for MiDAS (RAD52 dependent, RAD51 and 53BP1 independent) and for BIR in interphase cells (RAD51 dependent and RAD52 independent with suppression by 53BP1) could be quite different due to a special regulation of DSB repair in mitotic cells. We also speculate that active MiDAS would lead to substantial jumping/template switching events associated with the BIR mechanism, contributing to genome instability.

Loss of 53BP1 impairs NHEJ ^30–32^, rendering cells more reliant on HDR mechanism for survival. In addition to hyperrecombination, inactivation of 53BP1 also shifts the recombination repair to the BIR-like mechanism, requiring PIF1. Consistently, we showed that PIF1 exhibits synthetic lethal interactions with the 53BP1 pathway, and inactivating PIF1 induces cell death in cells with compromised 53BP1 axis, including those with TIRR overexpression, which inactivates 53BP1 function ^77^. TIRR amplification is frequently observed in a wide range of cancers ^77,109,110^, with the highest frequencies found in breast invasive carcinoma (4.06%, cholangiocarcinoma (2.78%) and diffuse large B-cell lymphoma (2.08%), according to cancer genomics data sets from cBioPortal (Supplementary Figure 12d). Thus, targeting PIF1 presents an attractive therapeutic strategy for treating tumors with TIRR amplification. Furthermore, since restored HR upon loss of 53BP1 activity in BRCA1-deficient cells is dependent on PIF1, depleting PIF1 overcomes PARPi resistance in cells deficient for both BRCA1 and 53BP1. Therefore, targeting the BIR pathway also provides a treatment avenue to overcome PARPi resistance in BRCA1-deficient cells resulting from the disruption of the 53BP1 pathway.

## Methods

### Cell cultures and lentiviral production

U2OS (human osteosarcoma) and HEK293T cells were obtained from the ATTC cell repository. RPE-1 WT cells were received from Dr. Stephen P. Jackson’s lab. UWB1 and UWB1 reconstituted with BRCA1 cells were received from Dr. Lee Zou’s lab. Cells were cultured in Dulbecco’s modified Eagle’s medium (DMEM; Gibco) supplemented with 10% fetal bovine serum (FBS; GeminiBio.), 2mM L-glutamine (Sigma-Aldrich), and 1% penicillin-streptomycin containing glutamine (Gibco) at 37°C in a humid atmosphere containing 5% CO_2_.

For lentivirus production, HEK293T cells were co-transfected with a lentivirus based PLKO.1 vector and packaging vectors using the standard calcium chloride protocol.

### Plasmids construction and generation of reporter cell lines

The EGFP-PIF1 expression construct was generated by placing the EGFP tag on the N-terminus of human PIF1 cDNA and subcloned into the lentiviral vector pCDH-CMV-MCS-EF1-PURO (System Biosciences) at the NotI and NheI cloning sites. A stable cell line expressing EGFP-PIF1 was generated in the wild-type (WT) and *53BP1*-KO cells by lentiviral infection followed by puromycin (2μg/ml, 2 days) selection.

The HA-tagged PCNA-WT allele and the PCNA-K164R mutant allele were subcloned into pCDH-CMV-HA-MCS-EF1-PURO, with the shRNA target site (5’-TGGAGAACTTGGAAATGGAA) in PCNA disrupted by site-directed mutagenesis using the Quick Change system (Stratagene). Stable cell lines expressing HA-PCNA-WT or HA-PCNA-K164R were generated by lentiviral infection, followed by puromycin (2μg/ml, 2 days) selection, and confirmed by Western blotting.

To generate U2OS cell lines expressing 3xFlag-SMARCAD1-WT or 3xFlag-NΔ-SMARCAD1 deficient in the interaction with PCNA ^71^, the cDNA of *SMARCAD1*-WT or NΔ-*SMARCAD1* lacking the first 137 amino acids was subcloned into pCDH-CMV-3xFlag-EF1-Neo vector. The shRNA target site (CCAGCACCTTATGACAATTAA) was disrupted by site-directed mutagenesis (Stratagene) following the user protocol. Stable cell lines were generated by lentiviral infection of 3xFlag-SMARCAD1-WT or NΔ-SMARCAD1, followed by G418 (400µg/ml, 4 days) selection. The endogenous SMARCAD1 is depleted with shRNA, which targets the endogenous gene but not exogenously expressed alleles.

The lentiviral vector carrying 3xFlag-TIRR was constructed by subcloning *TIRR* cDNA into the pCDH-CMV-3xFlag-PURO vector. Stable U2OS (EGFP-BIR/LTGC) and U2OS (EGFP-HR/STGC) cell lines expressing 3xFlag-TIRR were generated by lentiviral expression.

The U2OS (EGFP-BIR/LTGC) and U2OS (EGFP-Flex1-BIR) reporter cell lines were described previously ^19^. To generate the EGFP-HR/STGC reporter, a single I-SceI site along with two in-frame stop codons was inserted in the middle of the EGFP open reading frame, 306 bp away from the starting codon that was under the control of the CMV promoter (the recipient cassette), resulting in a disruption of the expression of EGFP. A donor template (650 bp in size) carrying the internal part of EGFP (iEGFP), corresponding to the 314 bp left and 315 bp right sequences of the I-SceI site in the recipient cassette, was placed 2.3 kb downstream of the recipient cassette. The U2OS (EGFP-HR/STGC) reporter cell line was generated by transfection of the EGFP-HR/STGC reporter into U2OS cells with polyethylenimine (PEI) using the standard protocol, followed by hygromycin B (100µg/ml) selection.

### Generation of knock out (KO) cell line by CRISPR

The mCherry marker was inserted into the Cas9 plasmid for sgRNA insertion (Addgene, #62988). A pair of sgRNAs (5’-ACCTTCTCAATAAAGTTGAT and 5’-TCCAATCCTGAACAAACAGC) targeting the *53BP1* intronic region and exon3 respectively, were individually sub-cloned into the mCherry-Cas9 all-in-one plasmid. The combination of two Cas9/gRNAs would cause a frameshift in the open reading frame of the 53BP1 protein. For generating BRCA1-ΔBRCT in U2OS cells with deletion of the two BRCT domains at the C-terminus of BRCA1 (Supplementary Figure 3b), a sgRNA (5’-TTCAGAGGGAACCCCTTACC) targeting the region upstream of the BRCT domains of *BRCA1* was integrated into mCherry-Cas9 all-in-one plasmid. U2OS *PIF1*-KO cells were described previously ^19^. For RPE-1 cells, the *TP53* gene was knocked-out using a pair of sgRNAs (5’-GCTATCTGAGCAGCGCTCA and 5’-AGACCTCAGGCGGCTCATA) sub-cloned into the mCherry-Cas9 all-in-one vector. Subsequently, we generated *PIF1*-KO in RPE-1 *TP53-*KO cells using a pair of sgRNAs (5’-CACTCACAGGCATCGGCTC and 5’-GGTCATTGACGAGATCTCAA) targeting PIF1 Exon 5 leading to an open reading frame shift, knocking out the PIF1 catalytic site. *RAD52-*KO in U2OS (EGFP-BIR/LTGC) cells was generated using a pair of sgRNAs (5’-TCCAGAAGGCCCTGAGGCAG and 5’-AGTAGCCGCATGGCTGGCGG) targeting *RAD52* exon 3 ^111^.

To generate KO cells, U2OS (EGFP-BIR/LTGC), U2OS (EGFP-HR/STGC) cells and RPE-1 cells were seeded into a 100 mm petri dish to achieve 80% confluency at the time of transfection. Transfection was performed using PEI following the standard protocol. 48 hours post-transfection, mCherry positive cells were sorted by flow cytometry. Single clones were isolated and screened by Sanger sequencing for identifying KO clones, followed by confirmation with Western blotting. For creating *53BP1*-KO/BRCA1-ΔBRCT cell lines, *53BP1*-KO was generated into the BRCA1-ΔBRCT cell lines similarly as described above.

### shRNA interference

Endogenous gene silencing was achieved via lentiviral infection using the pLKO.1-blast vector (Addgene #26655) to express corresponding shRNAs. The shRNA sequences targeting different genes are listed in the Supplementary Table1.

Lentiviruses for the indicated genes were harvested from 293T cells. They were concentrated with a lentivirus concentrator solution (40% PEG-8000 (W/V), 1.2M NaCl). Lentiviral infection was followed by blasticidin (10 μg/ml, 2 days) selection, and the knockdown efficiency of the targeted gene was verified by Western blotting and RT-qPCR.

### Immunoblotting

Cells were lysed in NETN buffer (20 mM Tris-HCl, pH 8.0, 100mM NaCl, 0.5 mM EDTA, 0.5% NP-40) containing aprotinin (2µg/ml) and PMSF (20µg/ml). 2xSDS loading buffer was added, and samples were boiled for 5min and separated on 6-15% SDS-PAGE. Antibodies used were: 53BP1 (NB100-305, Novus Biologicals), FLAG (F1804, Sigma-Aldrich), PCNA (SC-56, Santa Cruz), HA (E10176EF, Covance), PRIM1 (10773-1-AP, Proteintech), RPA2 (NA19L, Calbiochem), SMARCAD1 (A5850, ABclonal), and KU70 (E-5, SC-17789, Santa Cruz Biotechnology). MRE11 antibody was described previously ^112^.

### GST-Immunoprecipitation

GST-fused PCNA or PCNA-Ub [C-terminal Ub, ^66^] was expressed in Rosetta and affinity-purified with glutathione-Sepharose 4B (GE Healthcare). SDS-PAGE electrophoresis and Coomassie blue staining were used to determine the purity and concentration of the protein bound to the glutathione-Sepharose beads. 293T cells transfected with pCDH-CMV-3xFlag-PIF1-neo plasmid were lysed in NETN [150 mM NaCl, 1 mM EDTA, 20 mM Tris-HCl (pH 8.0), 0.5% NP-40] containing protease inhibitors, aprotinin (2µg/ml) and PMSF (20µg/ml) and then incubated with GST-PCNA-bound Sepharose beads for 3 hours at 4°C. After extensive wash, pull-down samples were subject to Western blotting and Coomassie blue analysis.

### Growth curve and cell viability assay

Cell proliferation was determined by growth curves ^113^. Briefly, U2OS and RPE1 cells as well as their derived cell lines were seeded at a density of 1 × 10^4^ cells in 6 well cell culture dishes. Cell proliferation was assessed by counting trypsinized cells using Countess™ II FL automated cell counter (Thermo Fisher) every 24 hours. The cell number was normalized to that on day 1.

For the cell viability assay, cells were seeded in 96-well plates at a density of 2000 cells per well and treated with indicated concentrations of Olaparib for 72 hours. Subsequently, 100 μl of cell medium from each well was mixed with 20 μl Cell Counting Kit-8 (CCK-8, Dojindo) and incubated at 37°C for at least 2 hours. Cell viability was determined by measuring the emission at 490 nm using 800TS Microplate Reader (BioTek).

### Fluorescence-activated cell sorting (FACS) and tract length analysis

U2OS cells harboring EGFP-BIR/LTGC, EGFP-HR/STGC and EGFP-BIR-Flex1 reporters were infected with concentrated lentiviruses expressing either an empty vector or vectors expressing indicated shRNAs. 24 hours post infection, cells were treated with blasticidin (10 μg/ml, 2 days). To induce DSBs, U2OS (EGFP-HR/STGC) cells were infected with lentiviruses encoding I-SceI (pCDH-CMV-I-SceI-EF1-PURO), and U2OS (EGFP-LTGC/BIR) cells with lentiviruses of I-SceI or Cas9^WT^ or Cas9^D10A^ along with gRNA (5’-GTAGGAATTCAGTTACGCT) from the Cas9^WT^ vector derived from lentiCRISPR V2 (Addgene, #52961) or the Cas9^D10A^ vector ^19^. 96 hours after infection, cells were analyzed for EGFP positive events using a BD Accuri C6 flow cytometer.

To monitor BIR induced by replication stress in U2OS (EGFP-BIR-Flex1) reporter cell line, double thymidine block (2 mM, two cycles of 16h in drug with a 12h interval of drug-free medium in between) was performed to enrich cells in S-phase, followed by treatment with 2 mM HU for 24 hours. Alternatively, U2OS (EGFP-BIR-Flex1) cells were infected with lentiviruses encoding FANCM shRNA. Six days after HU treatment or FANCM shRNA lentiviral infection, EGFP-positive events were quantified by FACS analysis using a BD Accuri C6 flow cytometer.

To analyze BIR repair events, U2OS (EGFP-BIR/LTGC) WT or *53BP1*-KO cells were infected with lentiviruses expressing I-SceI, followed by puromycin selection (2µg/ml, 2 days). After 3 days of infection, EGFP positive cells were collected by FACS sorting, and spread on a 100 mm petri plate for 2-3 weeks to form single clones. Single clones were then transferred to 24 well plates, expanded, and their repair junction sequences analyzed using Sanger sequencing. The forward primer was fixed at the end of the FP region “GGCATGGACGAGCTGTACAAGTAA”, while the reverse primers were placed at 1kb intervals to the right side of the I-SceI cut site (Fig. 2b). The sequencing results were aligned to the predicted BIR/SDSA repair product (Fig. 2b) to determine the repair tract length, indels, and template jumping events.

### In situ proximity ligation assay (PLA)

Cells were seeded in a 6-well plate at 70% confluency. Cells were treated with 1 mM HU for 24 hours, or irradiated with 4 Gy and analyzed 2 hours after. For Polα inhibition, IR was performed immediately after initiating CD437 (10 μM) treatment. Cells were prepared by washing three times with PBS, followed by fixation with 2% paraformaldehyde for 20 min. Fixed samples were then permeabilized with 0.5% Triton X-100 for 10 min, followed by blocking with 3% BSA for 30 min. Blocked samples were incubated with primary antibodies overnight at 4°C. PLA was performed using the Duolink PLA technology (Sigma-Aldrich) following the manufacturer’s protocol. Coverslips were affixed onto glass slides using ProLong Gold antifade mountant (Invitrogen) with DAPI. The PLA signals were visualized as distinct fluorescence spots, and images were captured with an Olympus confocal microscope using a 60X objective. The number of PLA signals was quantified using CellProfiler 4.2.6 software.

Antibodies used for PLA include: FLAG (F1804, Sigma-Aldrich), FLAG (AE004, ABclonal), HA (E10176EF, Covance), PCNA (SC-56, Santa Cruz), PCNA (10205, Proteintech), PCNA K164Ub (13439, Cell Signaling Technology), γH2AX (05636, Upstate), γH2AX (07164, Upstate), RAD51 (05-530-I, Santa Cruz).

### RT–qPCR

Total RNA was extracted from the cell lines using the RNeasyMini Kit (Qiagen), following the manufacturer’s instructions. cDNA was synthesized by reverse transcription using the iScript cDNA synthesis kit (Bio-Rad). SYBR qPCR mix (Vazyme) was used to perform RT-qPCR on a Bio-Rad IQ5 real-time PCR system. The primer sequences are listed in the Supplementary Table 2.

### Laser microirradiation and live-cell imaging

The recruitment of EGFP-PIF1 to DSB sites in live cell nuclei was monitored following laser-induced microirradiation in WT and *53BP1*-KO cells ^114,115^. Briefly, a laser stripe was created using a femtosecond pulsed laser at 780 nm, coupled to a Zeiss Axiovert fluorescence microscope ^116,117^. After laser ablation, time-lapse live cell images were captured at various time intervals to measure the recruitment of EGFP-PIF1 to the laser-induced damaged sites. ImageJ software (National Institutes of Health) was used to quantify the fluorescence intensities at the microirradiated sites. The absolute intensity at the indicated time points was calculated by subtracting the background fluorescence from the fluorescence intensity generated at the damaged site. Each data point is an average of at least 5 independent measurements, with error bars showing the standard deviation (SD).

### Detection of EdU incorporation in mitosis

MiDAS was performed according to standard protocols^113,118^. Briefly, U2OS cells were treated with APH (0.4 μM) and RO-3306 (7 μM) for 16 hours to synchronize cells to the late G2 phase. Then cells were washed with cold PBS, released into fresh DMEM medium at 37°C within 5 minutes, followed by replacing the medium containing 20 μM EdU and 0.1 μg/ml Colcemid for incubation at 37°C for 60 minutes. Cells were subsequently shaken-off and resuspended in 75 mM KCl for 20 min at 37°C. Swollen mitotic cells were collected and fixed in a fixative solution (3:1 ratio of methanol and acetic acid) at room temperature for at least 30 minutes, then dropped onto pre-cooled slides from a height. Fixed cells were left overnight at room temperature. EdU incorporation was detected using the Click-IT EdU Alexa Fluor 488 Imaging Kit (Invitrogen). The Click-iT reaction was terminated with blocking buffer (3% BSA in PBS), and chromosome staining was performed with DAPI. Images were taken on Nikon ECLIPSE Ni-L microscope.

### Immunostaining

For PCNA-related fluorescent staining, cells fixed in acetone–methanol (1:1 v/v) and extracted with PBS containing 0.1% Tween-20. For other staining, cells were pre-extracted with 0.5% Trion X-100 in cold PBS for 5 mins, and then fixed with 1% formaldehyde for 20 minutes at room temperature. Fixed cells were blocked with 5% BSA in PBS for 30 minutes and then incubated overnight with indicated primary antibodies diluted in 5% BSA at 4°C. On the next day, cells were washed with 0.1% PBST and further incubated with secondary antibodies in the presence of DAPI. Finally, slides were sealed with mounting media (P36930, Invitrogen) to prevent quenching. Images were taken on Nikon ECLIPSE Ni-L microscope.

Antibodies used were as followed: FLAG (AE004, Abclonal, 1:1000), PCNA (SC-56, Santa Cruz, 1:500), γH2AX (05636, Upstate, 1:500), γH2AX (07164, Upstate, 1:500), RAD51 (05-530-I, Santa Cruz, 1:500), RAD51 (ab46981-100, Abcam, 1:500)

### End resection assay

End resection assay was performed in U2OS (EGFP-BIR/LTGC) cells after I-SceI cleavage as described ^119^ with modifications. Cells were infected with lentivirus expressing I-SceI to induce DSBs in the EGFP-BIR/LTGC reporter. Two days after infection, cells were harvested for genomic DNA extraction using Puregene Kits (QIAGEN). Genomic DNA (1 ug) was digested with restriction enzyme BsiHKAI or SacII (control) overnight, and digested genomic DNA was used as template for qPCR with indicated primer listed below.

0.25kb:

ER_L1F GCTCCAACACCCCAACATCTTCGAC
ER_L1R CGGTACTTCGTCCACAAACACAACTCC

### Statistical analysis

Statistical analysis was performed using GraphPad Prism9 and Microsoft Excel. In all experiments, error bars represent standard error of the mean (SEM) of at least three independent experiments. Significant differences between the two groups were determined by unpaired Student’s t-test (sample sizes ≤30) or Mann-Whitney U test (sample sizes greater than 30). P values are indicated in the figures.

### Data availability statement

All data supporting the findings of this study are available in figshare with the identifier (doi.org/10.6084/m9.figshare.26341645) within the paper and its Supplementary Information. Source data are provided with this paper.

## Acknowledgements

We would like to thank Dr. Cyril Sanders (University of Sheffield, UK) for providing cDNA of human PIF1, Dr. Jacqueline Mermoud (Babraham Institute, UK), Dr. Zihua Gong (Cleveland Clinic Lerner Research Institute) for providing the Flag-SMARCAD1 expression plasmid, Dr. Stephen P. Jackson for providing RPE-1 cells, and Dr. Lee Zou for UWB1 and UWB1 reconstituted with BRCA1 cell lines. Plasmids pLKO.1-blast vector (#26655), lentiCRISPR v2 (#52961), pSpCas9(BB)-2A-Puro (PX459) V2.0 (#62988) are from Addgene. This study is funded by NIH grants CA244912, GM141868 and CA187052 to X.W.

## Author contributions

S.B.S and Y.L. designed and performed experiments and analyzed the data; S.L. designed some experiments and established reporter analysis systems; Q.H., T.W., Y.S., T.N., I.I. and L.S. conducted some experiments. H.W. contributed to student supervision. X.W. is responsible for the overall project’s planning and experimental design. X.W., S.B.S and Y.L. wrote the manuscript.

## References

1 Aguilera, A. & Gomez-Gonzalez, B. Genome instability: a mechanistic view of its causes and consequences. Nat Rev Genet 9, 204–217, doi:nrg2268 [pii] 10.1038/nrg2268 (2008).

2 Negrini, S., Gorgoulis, V. G. & Halazonetis, T. D. Genomic instability--an evolving hallmark of cancer. Nat Rev Mol Cell Biol 11, 220–228, doi:10.1038/nrm2858 (2010).

3 Nik-Zainal, S. et al. Mutational processes molding the genomes of 21 breast cancers. Cell 149, 979–993, doi:10.1016/j.cell.2012.04.024 (2012).

4 Chiarle, R. et al. Genome-wide translocation sequencing reveals mechanisms of chromosome breaks and rearrangements in B cells. Cell 147, 107–119, doi:10.1016/j.cell.2011.07.049 (2011).

5 Stephens, P. J. et al. Complex landscapes of somatic rearrangement in human breast cancer genomes. Nature 462, 1005–1010, doi:10.1038/nature08645 (2009).

6 Heyer, W. D. Regulation of recombination and genomic maintenance. Cold Spring Harbor perspectives in biology 7, a016501, doi:10.1101/cshperspect.a016501 (2015).

7 Paques, F. & Haber, J. E. Multiple pathways of recombination induced by double-strand breaks in Saccharomyces cerevisiae. Microbiol Mol Biol Rev 63, 349–404 (1999).

8 Jasin, M. & Rothstein, R. Repair of strand breaks by homologous recombination. Cold Spring Harbor perspectives in biology 5, doi:10.1101/cshperspect.a012740 (2013).

9 Liu, L. & Malkova, A. Break-induced replication: unraveling each step. Trends Genet 38, 752–765, doi:10.1016/j.tig.2022.03.011 (2022).

10 Anand, R. P., Lovett, S. T. & Haber, J. E. Break-induced DNA replication. Cold Spring Harbor perspectives in biology 5, a010397, doi:10.1101/cshperspect.a010397 (2013).

11 Llorente, B., Smith, C. E. & Symington, L. S. Break-induced replication: what is it and what is it for? Cell Cycle 7, 859–864 (2008).

12 Bhargava, R., Onyango, D. O. & Stark, J. M. Regulation of Single-Strand Annealing and its Role in Genome Maintenance. Trends Genet 32, 566–575, doi:10.1016/j.tig.2016.06.007 (2016).

13 Wu, X. & Malkova, A. Break-induced replication mechanisms in yeast and mammals. Curr Opin Genet Dev 71, 163–170, doi:10.1016/j.gde.2021.08.002 (2021).

14 Kramara, J., Osia, B. & Malkova, A. Break-Induced Replication: The Where, The Why, and The How. Trends Genet 34, 518–531, doi:10.1016/j.tig.2018.04.002 (2018).

15 Epum, E. A. & Haber, J. E. DNA replication: the recombination connection. Trends Cell Biol 32, 45–57, doi:10.1016/j.tcb.2021.07.005 (2022).

16 Bhowmick, R., Hickson, I. D. & Liu, Y. Completing genome replication outside of S phase. Mol Cell 83, 3596–3607, doi:10.1016/j.molcel.2023.08.023 (2023).

17 Malkova, A., Naylor, M. L., Yamaguchi, M., Ira, G. & Haber, J. E. RAD51-dependent break-induced replication differs in kinetics and checkpoint responses from RAD51-mediated gene conversion. Mol Cell Biol 25, 933–944, doi:10.1128/MCB.25.3.933-944.2005 (2005).

18 Davis, A. P. & Symington, L. S. RAD51-dependent break-induced replication in yeast. Mol Cell Biol 24, 2344–2351 (2004).

19 Li, S. et al. PIF1 helicase promotes break-induced replication in mammalian cells. EMBO J, e104509, doi:10.15252/embj.2020104509 (2021).

20 Lydeard, J. R., Jain, S., Yamaguchi, M. & Haber, J. E. Break-induced replication and telomerase-independent telomere maintenance require Pol32. Nature 448, 820–823, doi:10.1038/nature06047 (2007).

21 Donnianni, R. A. & Symington, L. S. Break-induced replication occurs by conservative DNA synthesis. Proc Natl Acad Sci U S A 110, 13475–13480, doi:10.1073/pnas.1309800110 (2013).

22 Saini, N. et al. Migrating bubble during break-induced replication drives conservative DNA synthesis. Nature 502, 389–392, doi:10.1038/nature12584 (2013).

23 Wilson, M. A. et al. Pif1 helicase and Poldelta promote recombination-coupled DNA synthesis via bubble migration. Nature 502, 393–396, doi:10.1038/nature12585 (2013).

24 Deem, A. et al. Break-induced replication is highly inaccurate. PLoS Biol 9, e1000594, doi:10.1371/journal.pbio.1000594 (2011).

25 Sakofsky, C. J. et al. Break-induced replication is a source of mutation clusters underlying kataegis. Cell reports 7, 1640–1648, doi:10.1016/j.celrep.2014.04.053 (2014).

26 Sakofsky, C. J., Ayyar, S. & Malkova, A. Break-induced replication and genome stability. Biomolecules 2, 483–504, doi:10.3390/biom2040483 (2012).

27 Smith, C. E., Llorente, B. & Symington, L. S. Template switching during break-induced replication. Nature 447, 102–105, doi:10.1038/nature05723 (2007).

28 Li, Y. et al. Patterns of somatic structural variation in human cancer genomes. Nature 578, 112–121, doi:10.1038/s41586-019-1913-9 (2020).

29 Hadi, K. et al. Distinct Classes of Complex Structural Variation Uncovered across Thousands of Cancer Genome Graphs. Cell 183, 197–210 e132, doi:10.1016/j.cell.2020.08.006 (2020).

30 Setiaputra, D. & Durocher, D. Shieldin - the protector of DNA ends. EMBO Rep 20, doi:10.15252/embr.201847560 (2019).

31 de Lange, T. Shelterin-Mediated Telomere Protection. Annu Rev Genet 52, 223–247, doi:10.1146/annurev-genet-032918-021921 (2018).

32 Panier, S. & Boulton, S. J. Double-strand break repair: 53BP1 comes into focus. Nat Rev Mol Cell Biol 15, 7–18, doi:10.1038/nrm3719 (2014).

33 Bunting, S. F. et al. 53BP1 inhibits homologous recombination in Brca1-deficient cells by blocking resection of DNA breaks. Cell 141, 243–254, doi:S0092-8674(10)00285-0 [pii] 10.1016/j.cell.2010.03.012 (2010).

34 Ward, I. M. et al. 53BP1 is required for class switch recombination. J Cell Biol 165, 459–464, doi:10.1083/jcb.200403021 (2004).

35 Manis, J. P. et al. 53BP1 links DNA damage-response pathways to immunoglobulin heavy chain class-switch recombination. Nat Immunol 5, 481–487, doi:10.1038/ni1067 (2004).

36 Dimitrova, N., Chen, Y. C., Spector, D. L. & de Lange, T. 53BP1 promotes non-homologous end joining of telomeres by increasing chromatin mobility. Nature 456, 524–528, doi:10.1038/nature07433 (2008).

37 Chapman, J. R., Sossick, A. J., Boulton, S. J. & Jackson, S. P. BRCA1-associated exclusion of 53BP1 from DNA damage sites underlies temporal control of DNA repair. J Cell Sci 125, 3529–3534, doi:10.1242/jcs.105353 (2012).

38 Zimmermann, M., Lottersberger, F., Buonomo, S. B., Sfeir, A. & de Lange, T. 53BP1 regulates DSB repair using Rif1 to control 5’ end resection. Science 339, 700–704, doi:10.1126/science.1231573 (2013).

39 Escribano-Diaz, C. et al. A cell cycle-dependent regulatory circuit composed of 53BP1-RIF1 and BRCA1-CtIP controls DNA repair pathway choice. Mol Cell 49, 872–883, doi:10.1016/j.molcel.2013.01.001 (2013).

40 Chapman, J. R. et al. RIF1 is essential for 53BP1-dependent nonhomologous end joining and suppression of DNA double-strand break resection. Mol Cell 49, 858–871, doi:10.1016/j.molcel.2013.01.002 (2013).

41 Jaspers, J. E. et al. Loss of 53BP1 causes PARP inhibitor resistance in Brca1-mutated mouse mammary tumors. Cancer Discov 3, 68–81, doi:10.1158/2159-8290.CD-12-0049 (2013).

42 Cao, L. et al. A selective requirement for 53BP1 in the biological response to genomic instability induced by Brca1 deficiency. Mol Cell 35, 534–541, doi:S1097-2765(09)00548-6 [pii] 10.1016/j.molcel.2009.06.037 (2009).

43 Noordermeer, S. M. et al. The shieldin complex mediates 53BP1-dependent DNA repair. Nature 560, 117–121, doi:10.1038/s41586-018-0340-7 (2018).

44 Dev, H. et al. Shieldin complex promotes DNA end-joining and counters homologous recombination in BRCA1-null cells. Nat Cell Biol 20, 954–965, doi:10.1038/s41556-018-0140-1 (2018).

45 Gupta, R. et al. DNA Repair Network Analysis Reveals Shieldin as a Key Regulator of NHEJ and PARP Inhibitor Sensitivity. Cell 173, 972–988 e923, doi:10.1016/j.cell.2018.03.050 (2018).

46 Ghezraoui, H. et al. 53BP1 cooperation with the REV7-shieldin complex underpins DNA structure-specific NHEJ. Nature 560, 122–127, doi:10.1038/s41586-018-0362-1 (2018).

47 Mirman, Z. et al. 53BP1-RIF1-shieldin counteracts DSB resection through CST- and Polalpha-dependent fill-in. Nature 560, 112–116, doi:10.1038/s41586-018-0324-7 (2018).

48 Mirman, Z., Sasi, N. K., King, A., Chapman, J. R. & de Lange, T. 53BP1-shieldin-dependent DSB processing in BRCA1-deficient cells requires CST-Polalpha-primase fill-in synthesis. Nat Cell Biol 24, 51–61, doi:10.1038/s41556-021-00812-9 (2022).

49 Callen, E. et al. 53BP1 mediates productive and mutagenic DNA repair through distinct phosphoprotein interactions. Cell 153, 1266–1280, doi:10.1016/j.cell.2013.05.023 (2013).

50 Xu, G. et al. REV7 counteracts DNA double-strand break resection and affects PARP inhibition. Nature 521, 541–544, doi:10.1038/nature14328 (2015).

51 Boersma, V. et al. MAD2L2 controls DNA repair at telomeres and DNA breaks by inhibiting 5’ end resection. Nature 521, 537–540, doi:10.1038/nature14216 (2015).

52 Bouwman, P. et al. 53BP1 loss rescues BRCA1 deficiency and is associated with triple-negative and BRCA-mutated breast cancers. Nat Struct Mol Biol 17, 688–695, doi:nsmb.1831 [pii] 10.1038/nsmb.1831 (2010).

53 Ochs, F. et al. 53BP1 fosters fidelity of homology-directed DNA repair. Nat Struct Mol Biol 23, 714–721, doi:10.1038/nsmb.3251 (2016).

54 Zimmermann, M. & de Lange, T. 53BP1: pro choice in DNA repair. Trends Cell Biol 24, 108–117, doi:10.1016/j.tcb.2013.09.003 (2014).

55 Sakofsky, C. J. et al. Translesion Polymerases Drive Microhomology-Mediated Break-Induced Replication Leading to Complex Chromosomal Rearrangements. Mol Cell 60, 860–872, doi:10.1016/j.molcel.2015.10.041 (2015).

56 Nacson, J. et al. BRCA1 Mutation-Specific Responses to 53BP1 Loss-Induced Homologous Recombination and PARP Inhibitor Resistance. Cell reports 24, 3513–3527 e3517, doi:10.1016/j.celrep.2018.08.086 (2018).

57 Zhang, F. et al. PALB2 links BRCA1 and BRCA2 in the DNA-damage response. Curr Biol 19, 524–529, doi:10.1016/j.cub.2009.02.018 (2009).

58 Sy, S. M., Huen, M. S. & Chen, J. PALB2 is an integral component of the BRCA complex required for homologous recombination repair. Proc Natl Acad Sci U S A 106, 7155–7160, doi:10.1073/pnas.0811159106 (2009).

59 Wang, H. et al. The concerted roles of FANCM and Rad52 in the protection of common fragile sites. Nat Commun 9, 2791, doi:10.1038/s41467-018-05066-y (2018).

60 Nakamura, K. et al. Genetic dissection of vertebrate 53BP1: a major role in non-homologous end joining of DNA double strand breaks. DNA Repair (Amst) 5, 741–749, doi:10.1016/j.dnarep.2006.03.008 (2006).

61 Callen, E. et al. 53BP1 Enforces Distinct Pre- and Post-resection Blocks on Homologous Recombination. Mol Cell 77, 26–38 e27, doi:10.1016/j.molcel.2019.09.024 (2020).

62 Thakar, T. et al. Ubiquitinated-PCNA protects replication forks from DNA2-mediated degradation by regulating Okazaki fragment maturation and chromatin assembly. Nat Commun 11, 2147, doi:10.1038/s41467-020-16096-w (2020).

63 Paiano, J. et al. Role of 53BP1 in end protection and DNA synthesis at DNA breaks. Genes Dev 35, 1356–1367, doi:10.1101/gad.348667.121 (2021).

64 Han, T. et al. The antitumor toxin CD437 is a direct inhibitor of DNA polymerase alpha. Nat Chem Biol 12, 511–515, doi:10.1038/nchembio.2082 (2016).

65 Buzovetsky, O. et al. Role of the Pif1-PCNA Complex in Pol delta-Dependent Strand Displacement DNA Synthesis and Break-Induced Replication. Cell reports 21, 1707–1714, doi:10.1016/j.celrep.2017.10.079 (2017).

66 Pastushok, L., Hanna, M. & Xiao, W. Constitutive fusion of ubiquitin to PCNA provides DNA damage tolerance independent of translesion polymerase activities. Nucleic Acids Res 38, 5047–5058, doi:10.1093/nar/gkq239 (2010).

67 Dungrawala, H. et al. The Replication Checkpoint Prevents Two Types of Fork Collapse without Regulating Replisome Stability. Mol Cell 59, 998–1010, doi:10.1016/j.molcel.2015.07.030 (2015).

68 Liu, W., Krishnamoorthy, A., Zhao, R. & Cortez, D. Two replication fork remodeling pathways generate nuclease substrates for distinct fork protection factors. Sci Adv 6, doi:10.1126/sciadv.abc3598 (2020).

69 Her, J., Ray, C., Altshuler, J., Zheng, H. & Bunting, S. F. 53BP1 Mediates ATR-Chk1 Signaling and Protects Replication Forks under Conditions of Replication Stress. Mol Cell Biol 38, doi:10.1128/MCB.00472-17 (2018).

70 Schmid, J. A. et al. Histone Ubiquitination by the DNA Damage Response Is Required for Efficient DNA Replication in Unperturbed S Phase. Mol Cell 71, 897–910 e898, doi:10.1016/j.molcel.2018.07.011 (2018).

71 Lo, C. S. Y. et al. SMARCAD1-mediated active replication fork stability maintains genome integrity. Sci Adv 7, doi:10.1126/sciadv.abe7804 (2021).

72 Densham, R. M. et al. Human BRCA1-BARD1 ubiquitin ligase activity counteracts chromatin barriers to DNA resection. Nat Struct Mol Biol 23, 647–655, doi:10.1038/nsmb.3236 (2016).

73 Costelloe, T. et al. The yeast Fun30 and human SMARCAD1 chromatin remodellers promote DNA end resection. Nature 489, 581–584, doi:10.1038/nature11353 (2012).

74 Chakraborty, S. et al. SMARCAD1 Phosphorylation and Ubiquitination Are Required for Resection during DNA Double-Strand Break Repair. iScience 2, 123–135, doi:10.1016/j.isci.2018.03.016 (2018).

75 DelloRusso, C. et al. Functional characterization of a novel BRCA1-null ovarian cancer cell line in response to ionizing radiation. Molecular cancer research : MCR 5, 35–45, doi:10.1158/1541-7786.MCR-06-0234 (2007).

76 Yazinski, S. A. et al. ATR inhibition disrupts rewired homologous recombination and fork protection pathways in PARP inhibitor-resistant BRCA-deficient cancer cells. Genes Dev 31, 318–332, doi:10.1101/gad.290957.116 (2017).

77 Drane, P. et al. TIRR regulates 53BP1 by masking its histone methyl-lysine binding function. Nature 543, 211–216, doi:10.1038/nature21358 (2017).

78 Botuyan, M. V. et al. Mechanism of 53BP1 activity regulation by RNA-binding TIRR and a designer protein. Nat Struct Mol Biol 25, 591–600, doi:10.1038/s41594-018-0083-z (2018).

79 Wang, J. et al. Molecular basis for the inhibition of the methyl-lysine binding function of 53BP1 by TIRR. Nat Commun 9, 2689, doi:10.1038/s41467-018-05174-9 (2018).

80 Dai, Y., Zhang, A., Shan, S., Gong, Z. & Zhou, Z. Structural basis for recognition of 53BP1 tandem Tudor domain by TIRR. Nat Commun 9, 2123, doi:10.1038/s41467-018-04557-2 (2018).

81 Cejka, P. & Symington, L. S. DNA End Resection: Mechanism and Control. Annu Rev Genet 55, 285–307, doi:10.1146/annurev-genet-071719-020312 (2021).

82 Clouaire, T. et al. Comprehensive Mapping of Histone Modifications at DNA Double-Strand Breaks Deciphers Repair Pathway Chromatin Signatures. Mol Cell 72, 250–262 e256, doi:10.1016/j.molcel.2018.08.020 (2018).

83 An, L. et al. RNF169 limits 53BP1 deposition at DSBs to stimulate single-strand annealing repair. Proc Natl Acad Sci U S A 115, E8286–E8295, doi:10.1073/pnas.1804823115 (2018).

84 Findlay, S. et al. SHLD2/FAM35A co-operates with REV7 to coordinate DNA double-strand break repair pathway choice. EMBO J 37, doi:10.15252/embj.2018100158 (2018).

85 Choe, K. N. & Moldovan, G. L. Forging Ahead through Darkness: PCNA, Still the Principal Conductor at the Replication Fork. Mol Cell 65, 380–392, doi:10.1016/j.molcel.2016.12.020 (2017).

86 Guilliam, T. A. et al. Human PrimPol is a highly error-prone polymerase regulated by single-stranded DNA binding proteins. Nucleic Acids Res 43, 1056–1068, doi:10.1093/nar/gku1321 (2015).

87 Guilliam, T. A. et al. Molecular basis for PrimPol recruitment to replication forks by RPA. Nat Commun 8, 15222, doi:10.1038/ncomms15222 (2017).

88 Baris, Y., Taylor, M. R. G., Aria, V. & Yeeles, J. T. P. Fast and efficient DNA replication with purified human proteins. Nature 606, 204–210, doi:10.1038/s41586-022-04759-1 (2022).

89 Abe, T. et al. AND-1 fork protection function prevents fork resection and is essential for proliferation. Nat Commun 9, 3091, doi:10.1038/s41467-018-05586-7 (2018).

90 Vouzas, A. E. & Gilbert, D. M. Mammalian DNA Replication Timing. Cold Spring Harbor perspectives in biology 13, doi:10.1101/cshperspect.a040162 (2021).

91 Richards, L., Das, S. & Nordman, J. T. Rif1-Dependent Control of Replication Timing. Genes (Basel) 13, doi:10.3390/genes13030550 (2022).

92 Malone, E. G., Thompson, M. D. & Byrd, A. K. Role and Regulation of Pif1 Family Helicases at the Replication Fork. Int J Mol Sci 23, doi:10.3390/ijms23073736 (2022).

93 Pohl, T. J. & Zakian, V. A. Pif1 family DNA helicases: A helpmate to RNase H? DNA Repair (Amst) 84, 102633, doi:10.1016/j.dnarep.2019.06.004 (2019).

94 Ray Chaudhuri, A., et al. Replication fork stability confers chemoresistance in BRCA-deficient cells. Nature 535, 382–387, doi:10.1038/nature18325 (2016).

95 Byrum, A. K. et al. Mitotic regulators TPX2 and Aurora A protect DNA forks during replication stress by counteracting 53BP1 function. J Cell Biol 218, 422–432, doi:10.1083/jcb.201803003 (2019).

96 Osia, B. et al. Cancer cells are highly susceptible to accumulation of templated insertions linked to MMBIR. Nucleic Acids Res 49, 8714–8731, doi:10.1093/nar/gkab685 (2021).

97 Cortes-Ciriano, I. et al. Comprehensive analysis of chromothripsis in 2,658 human cancers using whole-genome sequencing. Nat Genet 52, 331–341, doi:10.1038/s41588-019-0576-7 (2020).

98 Bhowmick, R., Minocherhomji, S. & Hickson, I. D. RAD52 Facilitates Mitotic DNA Synthesis Following Replication Stress. Mol Cell 64, 1117–1126, doi:10.1016/j.molcel.2016.10.037 (2016).

99 Minocherhomji, S. et al. Replication stress activates DNA repair synthesis in mitosis. Nature 528, 286–290, doi:10.1038/nature16139 (2015).

100 Giunta, S., Belotserkovskaya, R. & Jackson, S. P. DNA damage signaling in response to double-strand breaks during mitosis. J Cell Biol 190, 197–207, doi:10.1083/jcb.200911156 (2010).

101 van Vugt, M. A. et al. A mitotic phosphorylation feedback network connects Cdk1, Plk1, 53BP1, and Chk2 to inactivate the G(2)/M DNA damage checkpoint. PLoS Biol 8, e1000287, doi:10.1371/journal.pbio.1000287 (2010).

102 Nelson, G., Buhmann, M. & von Zglinicki, T. DNA damage foci in mitosis are devoid of 53BP1. Cell Cycle 8, 3379–3383, doi:10.4161/cc.8.20.9857 (2009).

103 Peterson, S. E. et al. Cdk1 uncouples CtIP-dependent resection and Rad51 filament formation during M-phase double-strand break repair. J Cell Biol 194, 705–720, doi:10.1083/jcb.201103103 (2011).

104 Ayoub, N. et al. The carboxyl terminus of Brca2 links the disassembly of Rad51 complexes to mitotic entry. Curr Biol 19, 1075–1085, doi:10.1016/j.cub.2009.05.057 (2009).

105 Freire, R., van Vugt, M. A., Mamely, I. & Medema, R. H. Claspin: timing the cell cycle arrest when the genome is damaged. Cell Cycle 5, 2831–2834, doi:10.4161/cc.5.24.3559 (2006).

106 Esashi, F. et al. CDK-dependent phosphorylation of BRCA2 as a regulatory mechanism for recombinational repair. Nature 434, 598–604, doi:10.1038/nature03404 (2005).

107 Krajewska, M. et al. Forced activation of Cdk1 via wee1 inhibition impairs homologous recombination. Oncogene 32, 3001–3008, doi:10.1038/onc.2012.296 (2013).

108 Xu, Y. et al. 53BP1 and BRCA1 control pathway choice for stalled replication restart. Elife 6, doi:10.7554/eLife.30523 (2017).

109 Drane, P. & Chowdhury, D. TIRR and 53BP1-partners in arms. Cell Cycle 16, 1235–1236, doi:10.1080/15384101.2017.1337966 (2017).

110 Parnandi, N. et al. TIRR inhibits the 53BP1-p53 complex to alter cell-fate programs. Mol Cell 81, 2583–2595 e2586, doi:10.1016/j.molcel.2021.03.039 (2021).

111 Liu, S. et al. DNA repair protein RAD52 is required for protecting G-quadruplexes in mammalian cells. J Biol Chem 299, 102770, doi:10.1016/j.jbc.2022.102770 (2023).

112 Wu, X. et al. SV40 T antigen interacts with Nbs1 to disrupt DNA replication control. Genes Dev 18, 1305–1316, doi:10.1101/gad.1182804 18/11/1305 [pii] (2004).

113 Li, Y. et al. MutSbeta protects common fragile sites by facilitating homology-directed repair at DNA double-strand breaks with secondary structures. Nucleic Acids Res 52, 1120–1135, doi:10.1093/nar/gkad1112 (2024).

114 Truong, L. N. et al. Microhomology-mediated End Joining and Homologous Recombination share the initial end resection step to repair DNA double-strand breaks in mammalian cells. Proc Natl Acad Sci U S A 110, 7720–7725, doi:10.1073/pnas.1213431110 (2013).

115 He, J. et al. Rad50 zinc hook is important for the Mre11 complex to bind chromosomal DNA double-stranded breaks and initiate various DNA damage responses. J Biol Chem 287, 31747–31756, doi:10.1074/jbc.M112.384750 (2012).

116 Botvinick, E. L. & Berns, M. W. Internet-based robotic laser scissors and tweezers microscopy. Microsc Res Tech 68, 65–74, doi:10.1002/jemt.20216 (2005).

117 Kong, X. et al. Comparative analysis of different laser systems to study cellular responses to DNA damage in mammalian cells. Nucleic Acids Res 37, e68, doi:10.1093/nar/gkp221 (2009).

118 Garribba, L. et al. Inducing and Detecting Mitotic DNA Synthesis at Difficult-to-Replicate Loci. Method Enzymol 601, 45–58, doi:10.1016/bs.mie.2017.11.025 (2018).

119 Zhou, Y., Caron, P., Legube, G. & Paull, T. T. Quantitation of DNA double-strand break resection intermediates in human cells. Nucleic Acids Res 42, e19, doi:10.1093/nar/gkt1309 (2014).

