## SupplementaryFigure for "53BP1 deficiency leads to hyperrecombination using break-induced replication (BIR)"

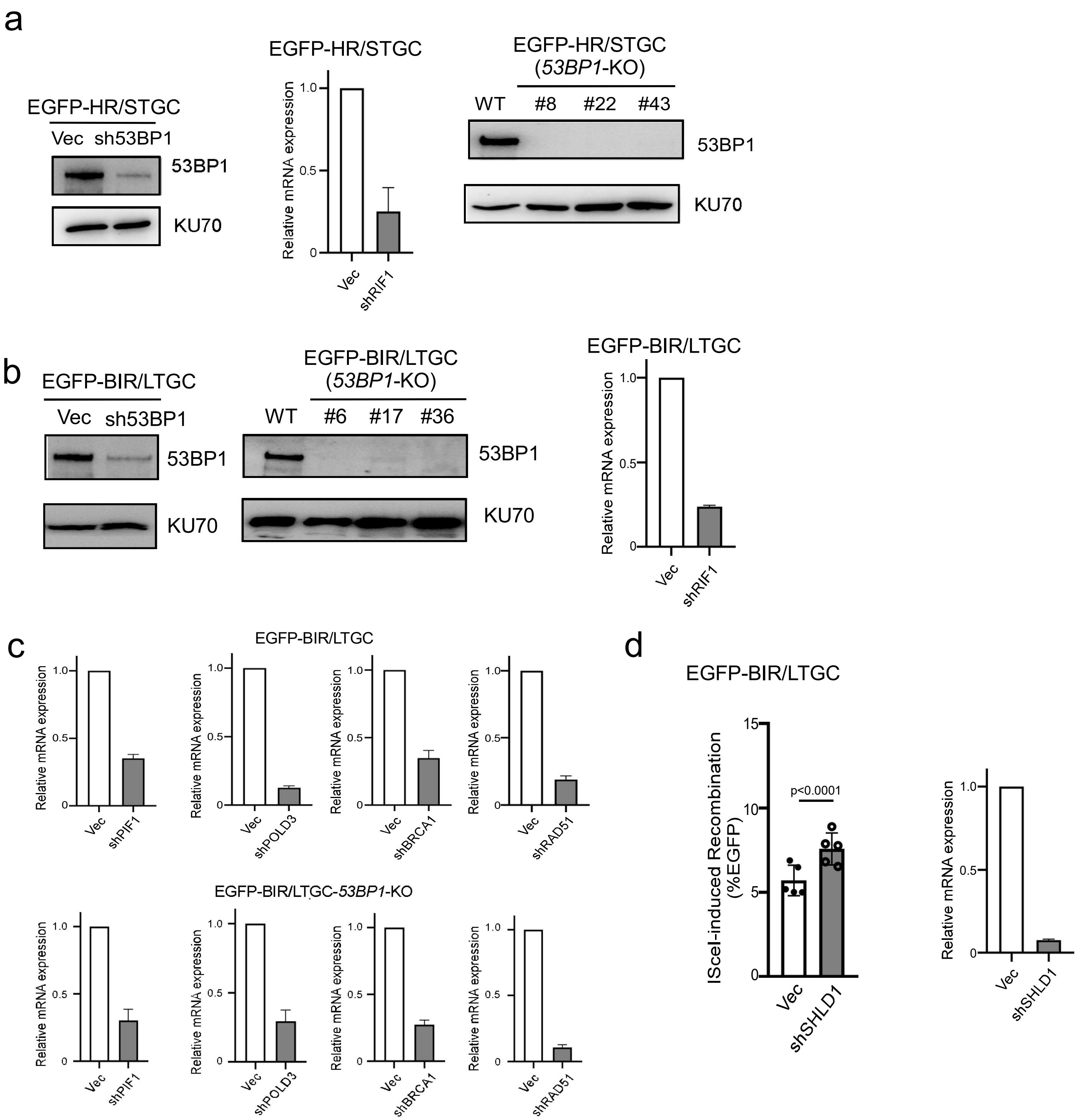


**Fig. S1. Related to Fig. 1.**

**Determining HR/STGC and BIR/LTGC frequency when the 53BP1 pathway is compromised.**

**a,** Western blot and qPCR analysis were conducted in U2OS (EGFP-HR/STGC) cells to assess the knockdown efficiency of shRNAs for 53BP1 (left) or RIF1 (middle), as well as the 53BP1 expression in *53BP1*-KO clones (right). Related to Figure 1a.

**b,** Western blot and qPCR analysis were conducted in U2OS (EGFP-BIR/LTGC) cells to assess the knockdown efficiency of shRNAs for 53BP1 (left) or RIF1 (right), as well as the 53BP1 expression in *53BP1*-KO clones (middle). Related to Figure 1c.

**c,** qPCR analysis of PIF1, POLD3, BRCA1 and RAD51 in U2OS (EGFP-BIR/LTGC) WT and *53BP1*-KO cells after lentiviral infection with corresponding shRNAs. Related to Figure 1d.

**d,** U2OS (EGFP-BIR/LTGC) cells, infected with lentiviruses encoding SHLD1 shRNA with a vector control, were assayed for BIR efficiency by determining the percentage of EGFP positive cells with FACS analysis, 5 days post-infection of I-SceI lentiviruses (left). qPCR analysis of SHLD1 expression (right) was determined to show SHLD1 shRNA knockdown efficiency. (n=5 replicates) a

| | WT-1 | WT-2 | *53BP1*-KO-17 | *53BP1*-KO-6 |
|---|---|---|---|---|
| Total green clones | **32** | **28** | **40** | **18** |
| **BIR-SDSA** (3.8 kb) | **7** (7/32:21.8%) | **5** (5/28:17.9%) | 9 (9/40: 22.5%) | 1 (1/18: 5.6%) |
| **BIR-EJ** (>0.9 kb, <3.8 kb) | **25** (25/32:78.2%) | **23** (23/28:82.1%) | 31 (31/40: 77.5%) | 17 (17/18:94.4%) |
| Average tract length | 2184.8 bp | 2135.6 bp | 2193 bp | 2070.9 bp |
| BIR-EJ with MH at EJ sites | 20 (20/25:80.0% | 15 (15/23:65.2%) | 25 (25/31:80.6%) | 12 (12/17:70.6%) |
| Size of MH | 1-6 bp (Ave: 2.5 bp) | 1-6 bp (Ave:2.7 bp) | 1-6bp (Ave:2.5 bp) | 1-6bp (Ave:2.7 bp) |
| Size of right-side deletions | 1-554 bp (Ave:39.12bp) | 1-235 bp (Ave:29.08 bp) | 1-6499bp (Ave:908.2 bp) | 1-3998bp (Ave:668.6 bp) |
| IN events in BIR-EJ | 4 (4/25:16%) | 5 (5/23:21.7%) | 5 (5/31:16.1%) | 4(4/17:23.5%) |
| IN size | 1-19 bp (Ave:6.25 bp) | 1-11 bp (Ave:7.6 bp) | 1-17bp (Ave:5.8 bp) | 2-8bp (Ave: 4.2 bp) |

b

| | WT-1 | WT-2 | *53BP1*-KO-17 | *53BP1*-KO-6 |
|---|---|---|---|---|
| BIR-EJ events with jumping | 3/25 (12%) | 4/23 (17.4%) | 7/31 (22.5%) | 4/17 (23.5%) |
| Jumping times | 1-2 times | 1-2 times | 1-2 times | 1-2 times |
| % of Jumps with MH | 4/5 (80%) | 4/6 (66.7%) | 7/9 (77.8%) | 5/5 (100%) |
| Size of MH at jumping sites | 1-3 bp (Ave:2 bp) | 1-3 bp (Ave:2.5 bp) | 1-3 bp (Ave:2 bp) | 2-5 bp (Ave:3.0 bp) |
| % of Jumps with IN | 0/5 (0%) | 1/6 (16.7%) | 2/9 (22.2%) | 0/5 (0%) |
| Size of IN at jumping sites | 0 bp (Ave: 0 bp) | 5 bp (Ave: 5 bp) | 2-17 bp (Ave:9.5 bp) | 0 bp (Ave: 0 bp) |

**Fig. S2. Related to Fig. 2**
**Analysis of BIR events after I-SceI cleavage.**
Two sets of experiments were performed to characterize the BIR events by analyzing single EGFP-positive clones derived from WT and *53BP1*-KO (EGFP-BIR/LTGC) reporter cell lines after I-SceI cleavage. A summary of the two sets of experiments on analysis of the repair products and the repair junctions (a) and the jumping/template switching events (b).

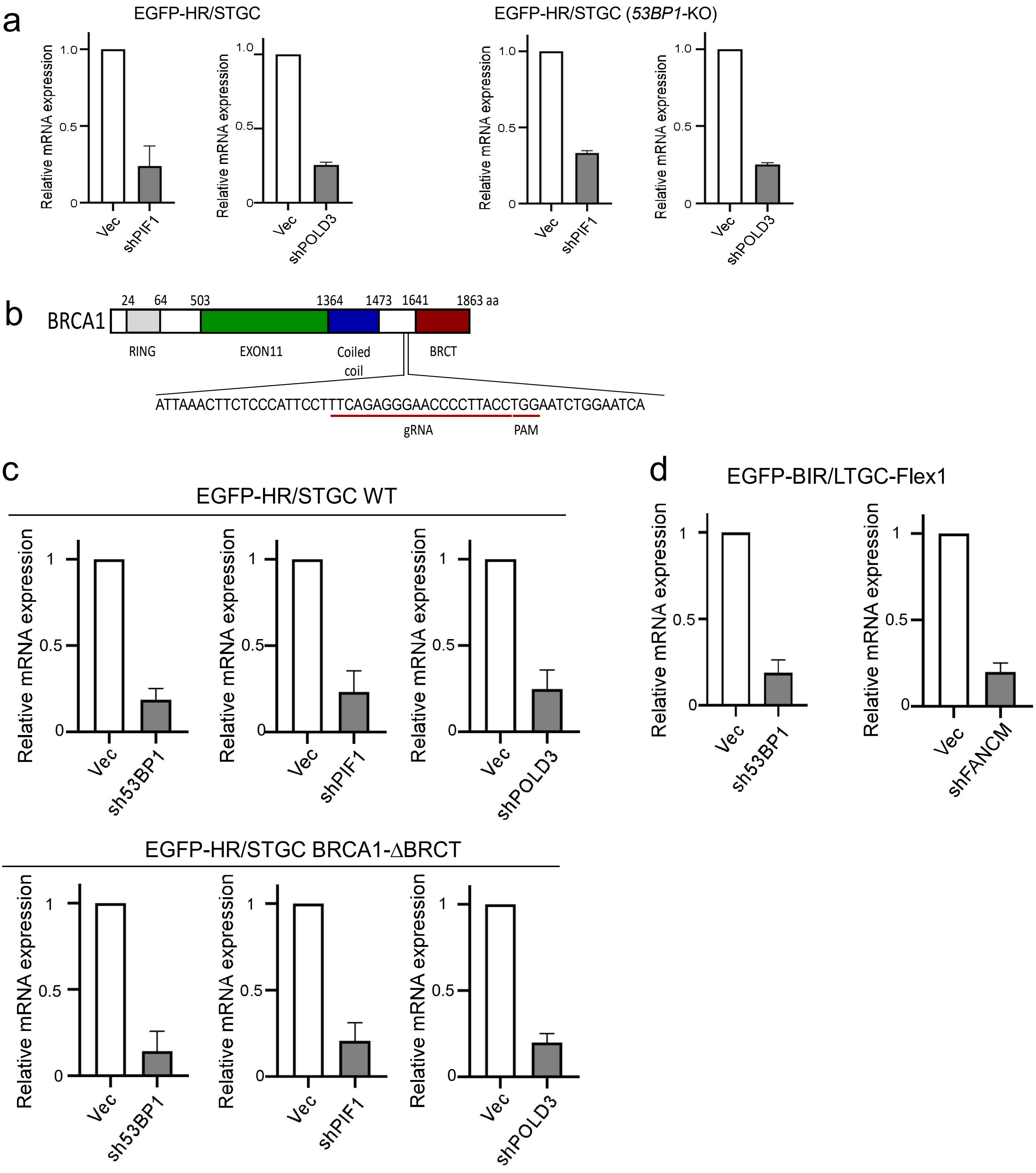


**Fig. S3. Related to Fig. 3.**

**Determining the dependence of hyperrecombination on PIF1 and POLD3 upon 53BP1 loss.**

**a,** qPCR analysis of PIF1 and POLD3 was conducted in U2OS (EGFP-HR/STGC) WT (left) and *53BP1*-KO (right) cells after lentiviral infection of PIF1 and POLD3 shRNAs. Related to Figure 3a.

**b,** Schematic drawing to show the domain structures of BRCA1 and the gRNA targeting site for generating the BRCA1-ΔBRCT allele by CRISPR/Cas9. Related to Figure 3b.

**c,** qPCR was performed to show depletion of 53BP1, PIF1 and POLD3 in U2OS (EGFP-HR/STGC) cells containing the WT allele (top) or BRCA1-ΔBRCT allele (bottom). Related to Figure 3c.

**d,** qPCR was performed to show depletion of 53BP1 (left) or FANCM (right) in U2OS (EGFP-BIR/LTGC-Flex1) cells. Related to Figure 3e.

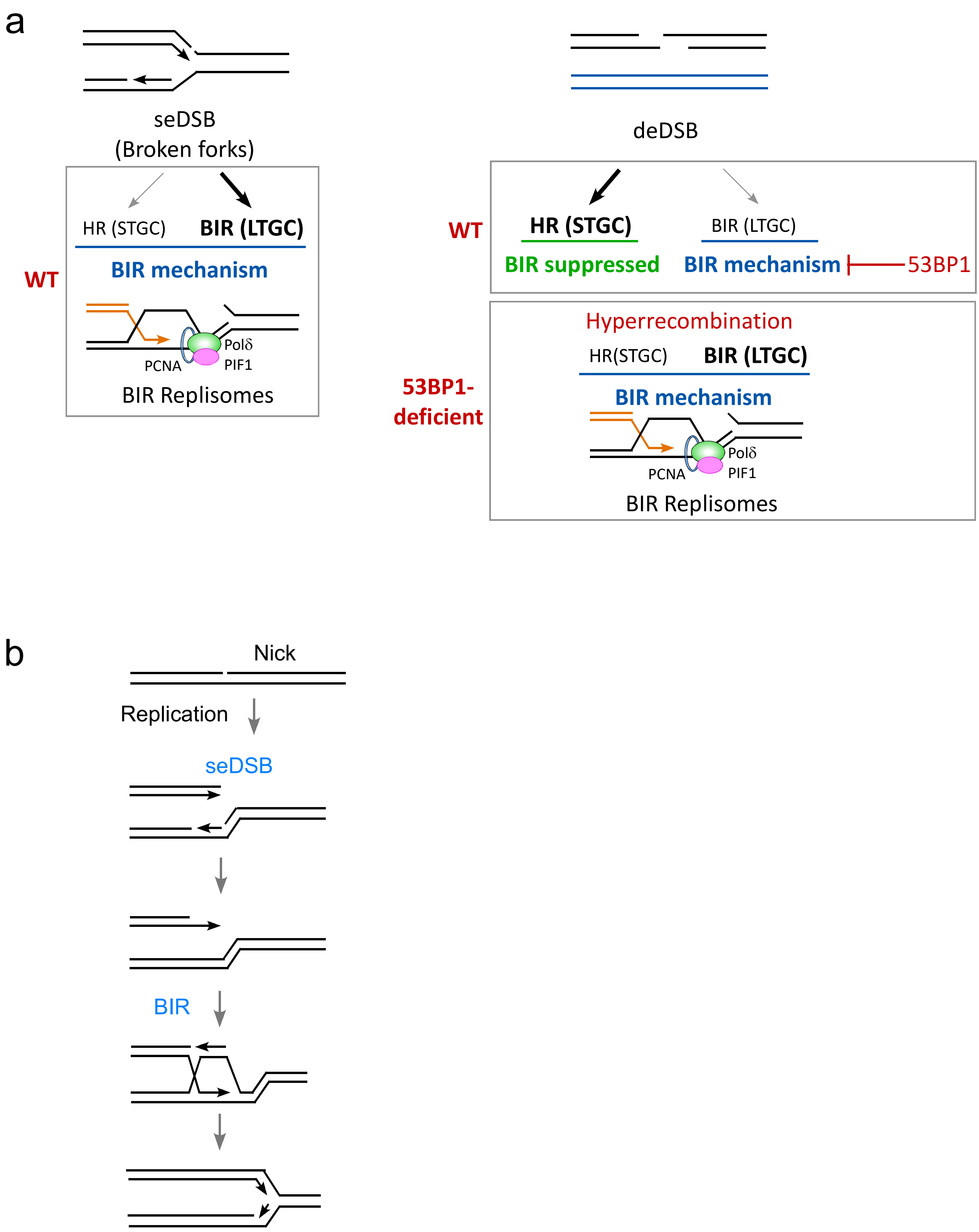


**Fig. S4.**

**Models for the use of HR/STGC and BIR/LTGC at seDSBs on broken replication forks and at deDSBs.**

**a,** Working models depicting the utilization of HR/STGC and BIR/LTGC at seDSBs on broken forks (left) and at deDSBs (right). The 53BP1 pathway is involved in restricting the use of BIR at deDSBs. See details in the main context.

**b,** A schematic drawing to show seDSB formation when replication encounters nicks on DNA and the repair of seDSBs by BIR.

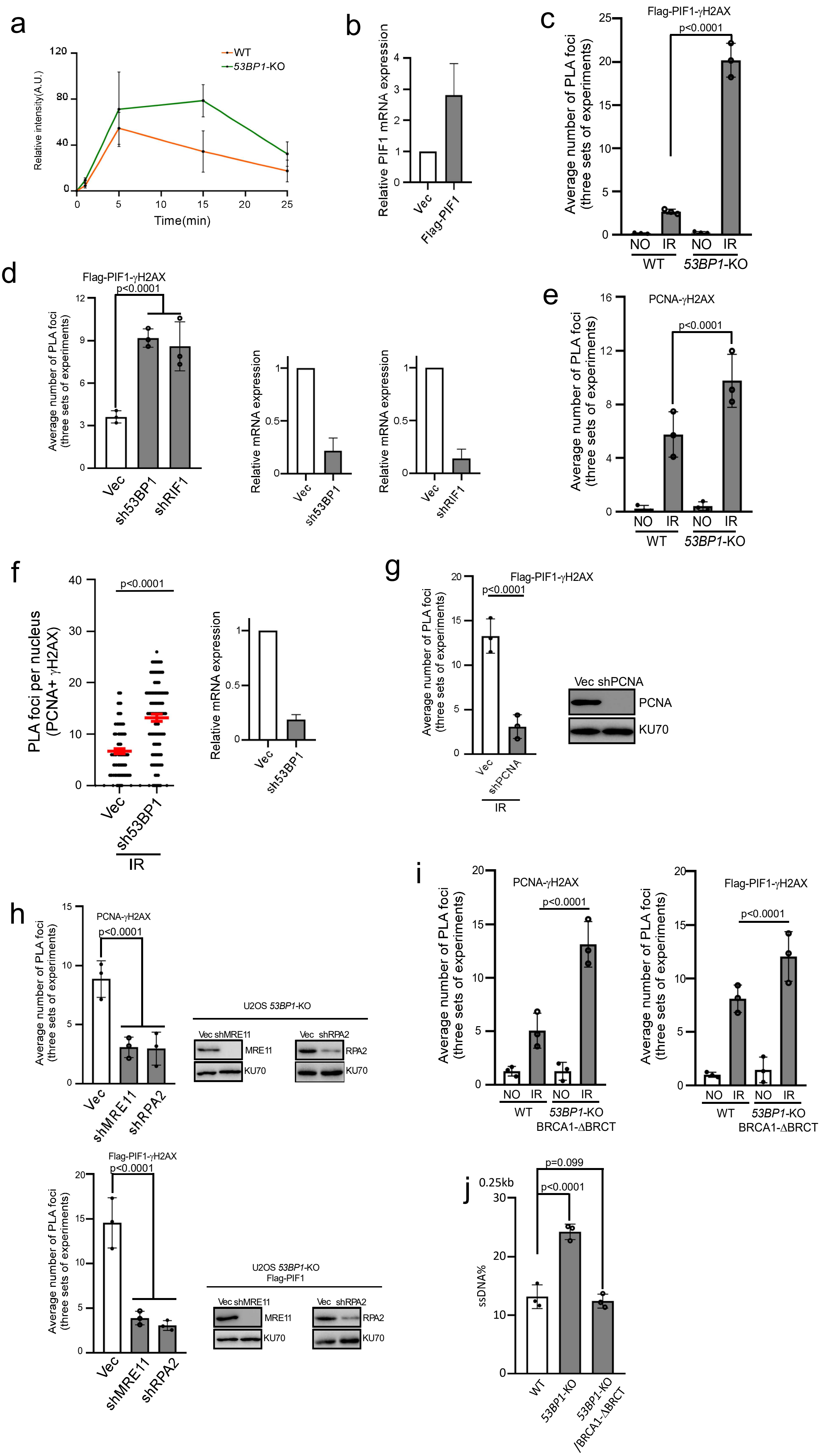
a
WT
53BP1-KO
Relative intensity(A.U.)
Time(min)
b
Relative PIF1 mRNA expression
Vec
Flag-PIF1
c
Flag-PIF1-γH2AX
p<0.0001
Average number of PLA foci (three sets of experiments)
NO IR NO IR
WT 53BP1-KO
d
Flag-PIF1-γH2AX
p<0.0001
Average number of PLA foci (three sets of experiments)
Vec sh53BP1 shRIF1
Relative mRNA expression
Vec sh53BP1
Relative mRNA expression
Vec shRIF1
e
PCNA-γH2AX
p<0.0001
Average number of PLA foci (three sets of experiments)
NO IR NO IR
WT 53BP1-KO
f
p<0.0001
PLA foci per nucleus (PCNA+ γH2AX)
Vec sh53BP1
IR
Relative mRNA expression
Vec sh53BP1
g
Flag-PIF1-γH2AX
p<0.0001
Average number of PLA foci (three sets of experiments)
Vec shPCNA
IR
Vec shPCNA
PCNA
KU70
h
PCNA-γH2AX
p<0.0001
Average number of PLA foci (three sets of experiments)
Vec shMRE11 shRPA2
U2OS 53BP1-KO
Vec shMRE11
MRE11
KU70
Vec shRPA2
RPA2
KU70
Flag-PIF1-γH2AX
p<0.0001
Average number of PLA foci (three sets of experiments)
Vec shMRE11 shRPA2
U2OS 53BP1-KO Flag-PIF1
Vec shMRE11
MRE11
KU70
Vec shRPA2
RPA2
KU70
i
PCNA-γH2AX
p<0.0001
Average number of PLA foci (three sets of experiments)
NO IR NO IR
WT 53BP1-KO
BRCA1-ΔBRCT
Flag-PIF1-γH2AX
p<0.0001
Average number of PLA foci (three sets of experiments)
NO IR NO IR
WT 53BP1-KO
BRCA1-ΔBRCT
j
0.25kb
p=0.099
p<0.0001
ssDNA%
WT
53BP1-KO
53BP1-KO /BRCA1-ΔBRCT

**Figure S5. Related to Figure 4.**
**PIF1 is accumulated at deDSBs after IR in cells deficient for the 53BP1 pathway.**
a, Quantification of the intensity of GFP-PIF1 recruited to the laser micro-irradiated sites. GFP-PIF1 images (n=5) were analyzed and quantified. The relative fluorescence intensity is plotted at each indicated time point. Error bars indicate SEM. Related to Figure 4a.
b, qPCR was performed to determine the total PIF1 expression levels in U2OS cells expressing vector (Vec) or exogenous Flag-PIF1. The expression level of the exogenous Flag-PIF1 was calculated to be ~1.8 fold as that of endogenous PIF1. Related to Figure 4b.
c, Three experiments of Flag-PIF1 and γH2AX PLA were performed in U2OS WT and *53BP1*-KO cells treated before and after or IR (4 Gy), with ~100 nuclei analyzed in each experiment. The average of three experiments and statistical significance are shown. Related to Figure 4b. (n=3 replicates)
d, Three experiments of Flag-PIF1 and γH2AX PLA were performed after depleting 53BP1 and RIF1 by shRNAs in U2OS cells expressing Flag-PIF1 after IR (4 Gy) treatment, with ~100 nuclei analyzed in each experiment (left). The average of three experiments and statistical significance are shown. qPCR was performed to show the depletion of 53BP1 and RIF1 (right). Related to Figure 4c. (n=3 replicates)
e, Three experiments of PCNA and γH2AX PLA were performed in U2OS WT and *53BP1*-KO cells treated before and after or IR (4 Gy), with ~100 nuclei analyzed in each experiment. The average of three experiments and statistical significance are shown. Related to Figure 4d. (n=3 replicates)
f, PLA of γH2AX with PCNA was performed after depleting 53BP1 in U2OS cells expressing Flag-PIF1 following IR (4 Gy) treatment (left). ~300 nuclei were analyzed in each sample. The expression of 53BP1was examined by qPCR (right). (n=300 cells)
g, Three experiments of Flag-PIF1 and γH2AX PLA were performed in U2OS cells after depleting PCNA by shRNA in U2OS cells after IR (4 Gy) treatment, with ~100 nuclei analyzed in each experiment. The average of three experiments and statistical significance are shown (left). Western blot analysis was performed to show PCNA depletion with KU70 as a loading control (right). Related to Figure 4e. (n=3 replicates)
h, Three experiments of PCNA and γH2AX PLA in *53BP1*-KO U2OS cells (top) or Flag-PIF1 and γH2AX PLA in *53BP1*-KO U2OS cells expressing Flag-PIF1 (bottom) were performed after depleting MRE11 or RPA by shRNAs after IR (4 Gy) treatment, with ~100 nuclei analyzed in each experiment. The average of three experiments and statistical significance are shown (left). Western blot analysis was performed to show MRE11 or RPA2 depletion with KU70 as a loading control. Related to Figure 4f. (n=3 replicates)
i, Three experiments of PCNA and γH2AX PLA in WT and *53BP1*-KO/BRCA1-ΔBRCT cells (left) or Flag-PIF1 and γH2AX PLA in *53BP1*-KO/BRCA1-ΔBRCT cells expressing Flag-PIF1 (right) were performed with or without treatment of IR (4 Gy), with ~100 nuclei analyzed in each experiment. The average of three experiments and statistical significance are shown. Related to Figure 4g. (n=3 replicates)
j, Genomic DNA was extracted from U2OS WT, *53BP1*-KO and *53BP*1-KO/BRCA1-ΔBRCT cells two days after I-SceI lentiviral infection. DNA end resection assay was performed as described in “Materials and Methods” at the positions 0.25 kb left to the I-SceI cleavage site. (n=3 replicates)

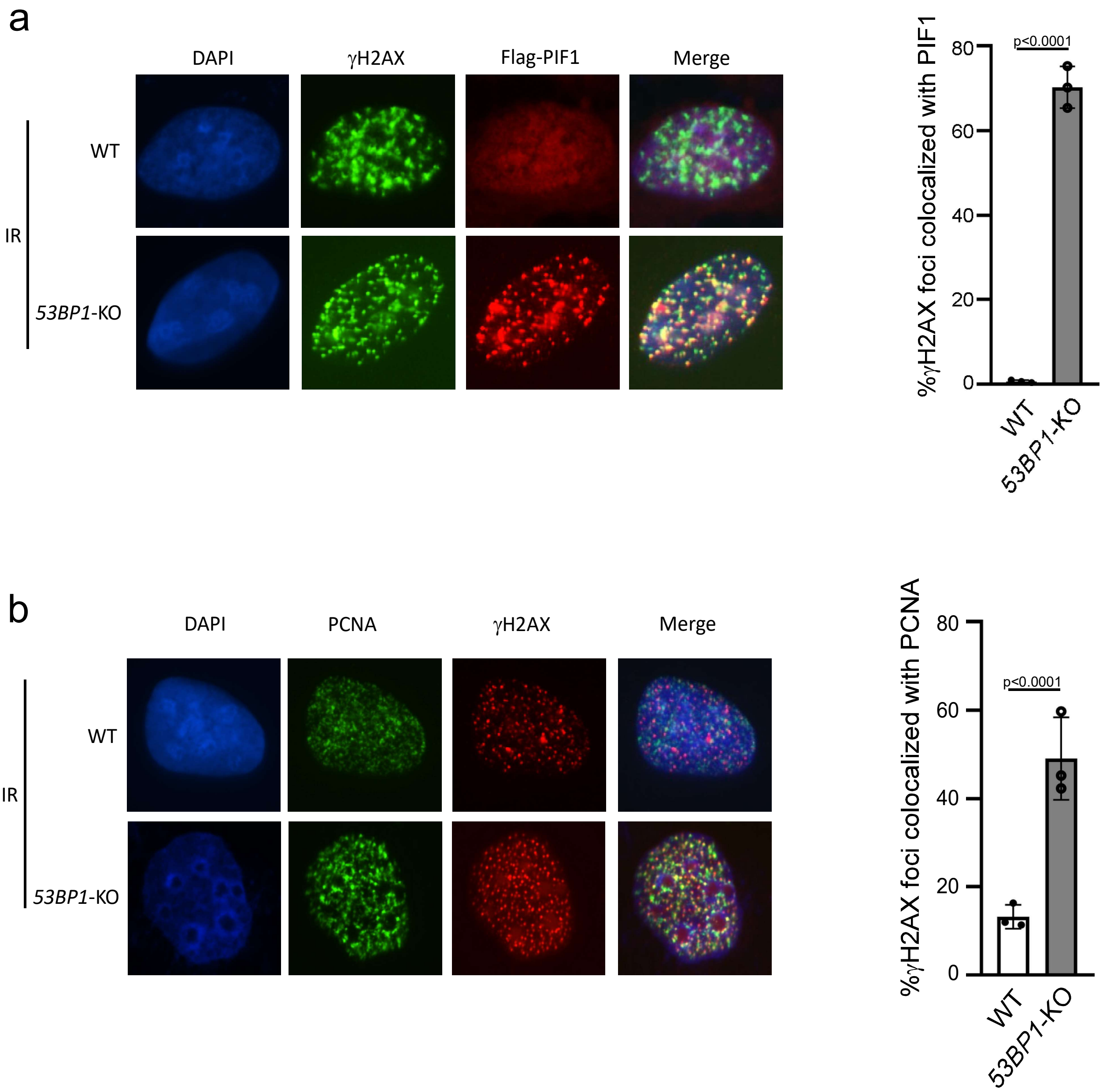


**Figure S6.**
**Colocalization of PIF1 and PCNA with γH2AX is increased in *53BP1*-KO cells after IR.**
U2OS WT and *53BP1*-KO cells stably expressing Flag-PIF1 were irradiated with 4-Gy IR, fixed at 2 hours after IR, followed by co-immunostaining of γH2AX (green) with Flag (red) (a), or γH2AX (red) with PCNA (green) (b), using DAPI for DNA staining (blue). The yellow dots represent colocalization between PIF1 and γH2AX (left). Quantification of colocalization of γH2AX with Flag-PIF1 or PCNA is shown(right). (n=3 replicates)

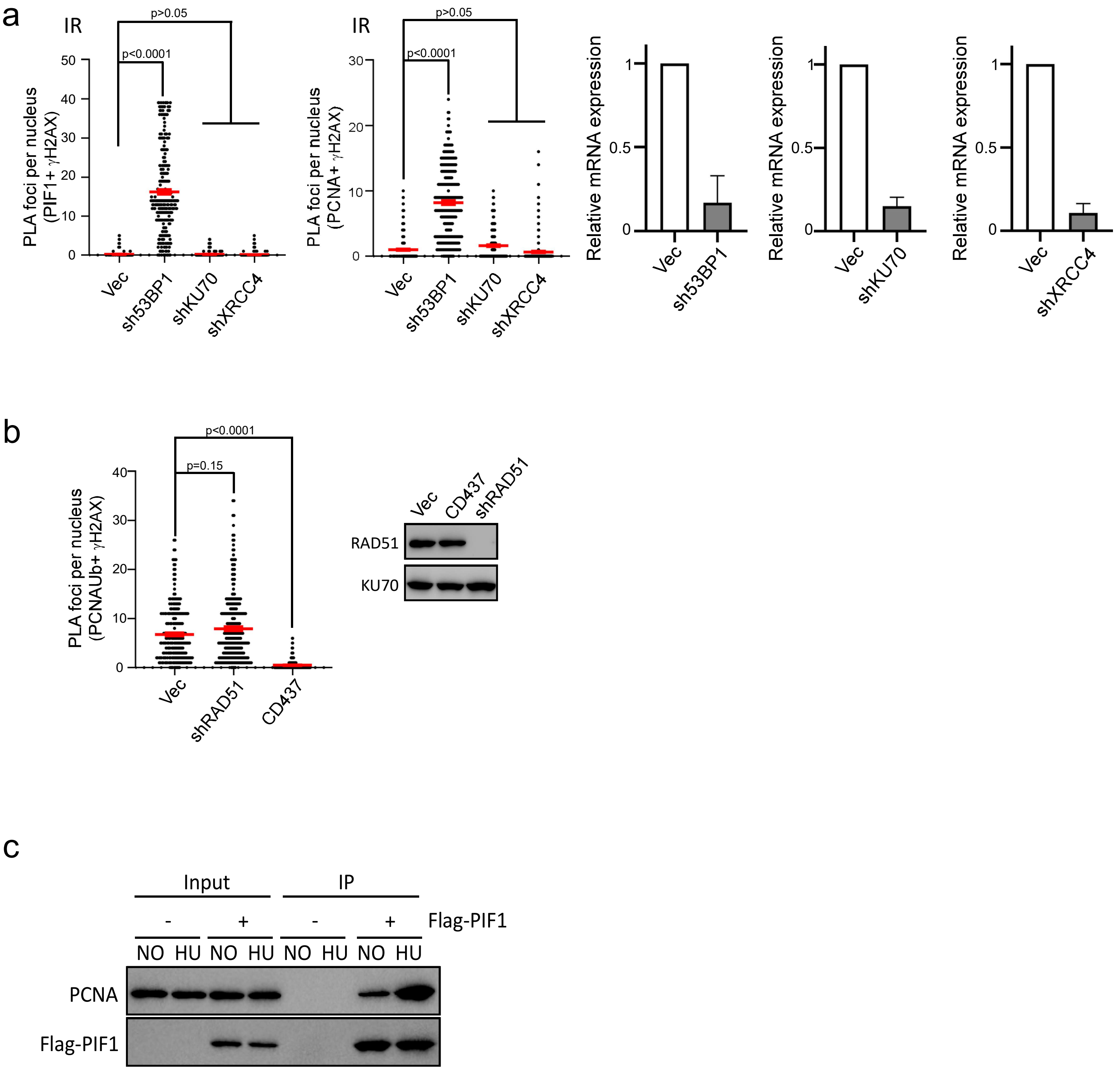


**Fig. S7.**

**Analysis of PIF1 and PCNA-Ub recruitment to DSBs and the interaction of PIF1 and PCNA.**

a, PLA of $\gamma$H2AX with Flag-PIF1 (left) or PCNA (right) after depleting 53BP1, KU70 or XRCC4 was performed in U2OS cells expressing Flag-PIF1 after IR (4Gy) treatment. ~100 nuclei were analyzed in each sample. The expression of 53BP1, KU70 and XRCC4 was examined by qPCR.(n=100 cells)

b, PLA of PCNA-Ub and $\gamma$H2AX was performed in U2OS cells after depleting RAD51 or treatment with POL$\alpha$ inhibitor CD437 (10 $\mu$M) (left). ~100 nuclei were analyzed in each sample. The expression of RAD51 was examined western blot (right).(n=100 cells)

c, Immunoprecipitation (IP) of Flag-PIF1 was performed before and after HU treatment (2 mM, 24h) in U2OS cells expressing Flag-PIF1 and the interaction of Flag-PIF1 with PCNA was revealed by anti-PCNA Western blot. Input is 5% of the lysates used for IP.

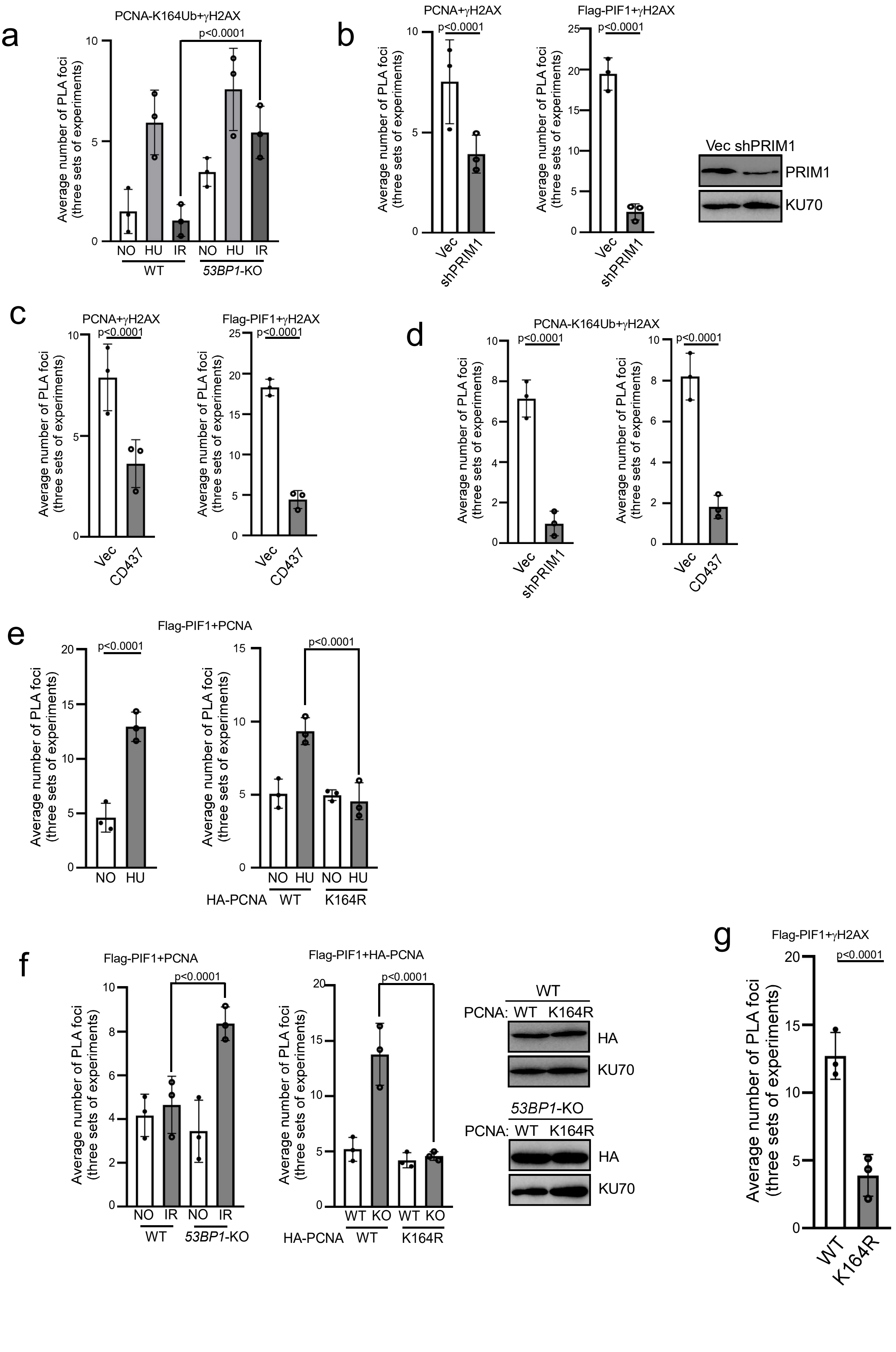
a
PCNA-K164Ub+γH2AX
p<0.0001
Average number of PLA foci (three sets of experiments)
NO HU IR NO HU IR
WT
53BP1-KO
b
PCNA+γH2AX
Flag-PIF1+γH2AX
p<0.0001
Vec
shPRIM1
Vec shPRIM1
PRIM1
KU70
c
PCNA+γH2AX
Flag-PIF1+γH2AX
CD437
d
PCNA-K164Ub+γH2AX
e
Flag-PIF1+PCNA
HA-PCNA
WT
K164R
f
Flag-PIF1+PCNA
Flag-PIF1+HA-PCNA
PCNA:
HA
g
Flag-PIF1+γH2AX

**Fig. S8. Related to Fig. 5.**
**PCNA ubiquitination and PIF1 recruitment to deDSBs are dependent on Polα-primase.**

a, Three experiments of PCNA-Ub (K164) and γH2AX PLA were performed in WT and *53BP1*-KO U2OS cells before and after treatment with HU (2 mM, 24h) or IR (4 Gy), with ~100 nuclei analyzed in each experiment. The average of three experiments and statistical significance are shown. Related to Figure 5a. (n=3 replicates)

b and c, Three experiments of PCNA and γH2AX PLA (left) or Flag-PIF1 and γH2AX PLA (middle) in U2OS *53BP1*-KO cells after depleting PRIM1 with vector as a control (b) or in the presence of CD437 (10 μM) (c) were performed after IR (4 Gy) treatment, with ~100 nuclei analyzed in each experiment. The average of three experiments and statistical significance are shown. Depletion of PRMI1 was examined by Western blot analysis using KU70 as a control in b. Related to Figure 5b and 5c. (n=3 replicates)

d, Three experiments of PCNA-Ub (K164A) and γH2AX PLA in U2OS *53BP1*-KO cells expressing PRIM1 shRNA (left) or treated with CD437 (10 μM) were performed after IR (4 Gy) treatment, with ~100 nuclei analyzed in each experiment. The average of three experiments and statistical significance are shown. Related to Figure 5d. (n=3 replicates)

e, Three PLA experiments of stably expressing Flag-PIF1 with endogenous PCNA (left), or with expressed HA-PCNA-WT and HA-PCNA-K164R were performed in U2OS cells with or without HU treatment (1 mM, 24h), with ~100 nuclei analyzed in each experiment. The average of three experiments and statistical significance are shown. Related to Figure 5e. (n=3 replicates)

f, Three PLA experiments were performed to assay for the interactions of stably expressing Flag-PIF1 with endogenous PCNA in U2OS WT and *53BP1*-KO cells (left) or with expressed HA-PCNA-WT and HA-PCNA-K164R (middle) with or without (4 Gy), with ~100 nuclei analyzed in each experiment. The average of three experiments and statistical significance are shown. Western blot analysis was performed to show the expression of HA-PCNA-WT and HA-PCNA-K164R in U2OS *53BP1*-KO (right) cells with KU70 as a loading control. Related to Figure 5f. (n=3 replicates)

g, Three experiments of Flag-PIF1 and γH2AX PLA were performed after IR treatment (4 Gy) in U2OS cells expressing Flag-PIF1 along with HA-PCNA-WT or K164R expression, with endogenous PCNA depleted by shRNA. ~100 nuclei were analyzed in each experiment. Related to Figure 5g. (n=3 replicates)

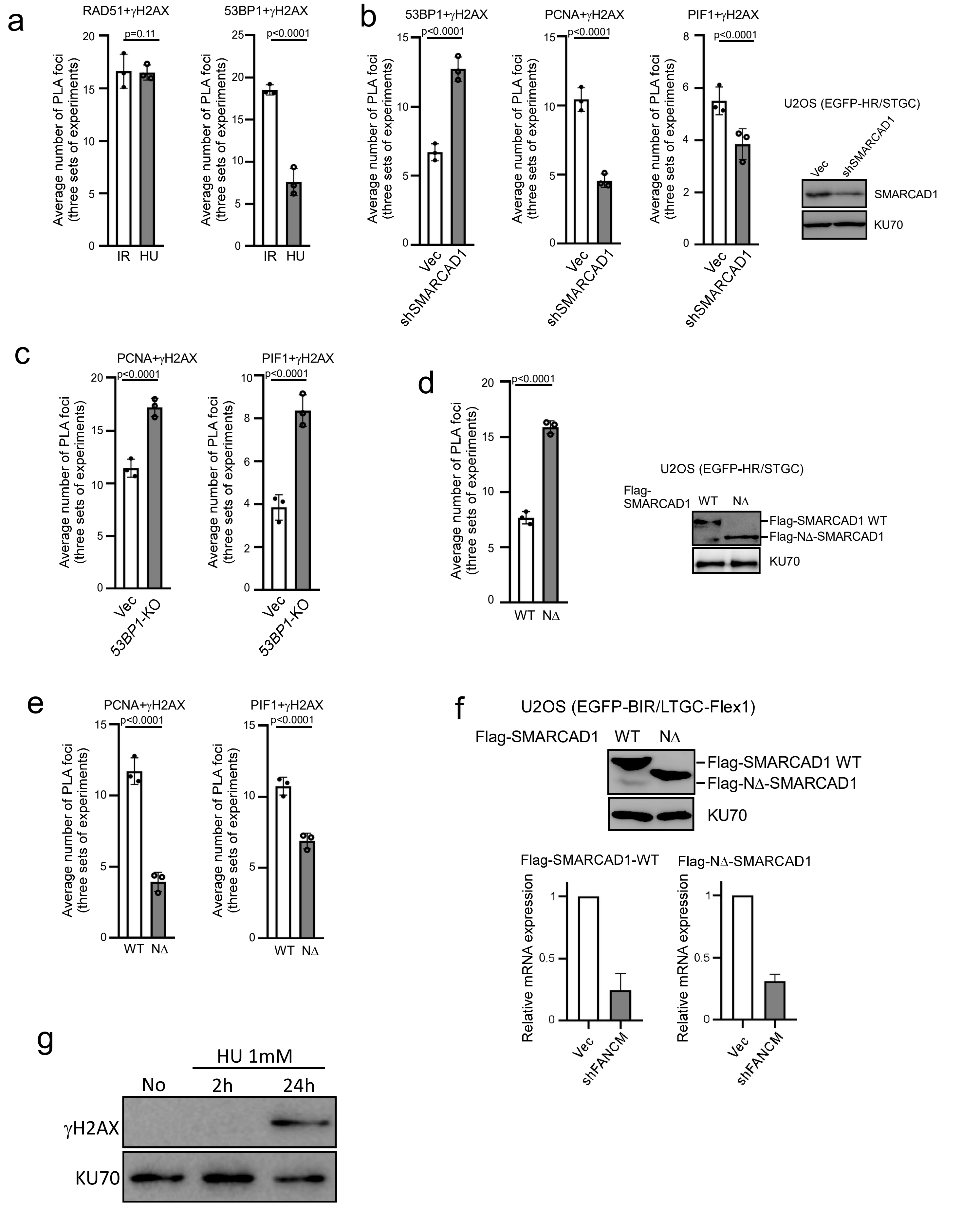
a
RAD51+γH2AX
53BP1+γH2AX
p=0.11
p<0.0001
Average number of PLA foci (three sets of experiments)
IR HU
b
53BP1+γH2AX
PCNA+γH2AX
PIF1+γH2AX
Vec
shSMARCAD1
U2OS (EGFP-HR/STGC)
SMARCAD1
KU70
c
PCNA+γH2AX
PIF1+γH2AX
Vec
53BP1-KO
d
U2OS (EGFP-HR/STGC)
Flag-SMARCAD1 WT NΔ
Flag-SMARCAD1 WT
Flag-NΔ-SMARCAD1
KU70
WT NΔ
e
PCNA+γH2AX
PIF1+γH2AX
f
U2OS (EGFP-BIR/LTGC-Flex1)
Flag-SMARCAD1 WT NΔ
Flag-SMARCAD1 WT
Flag-NΔ-SMARCAD1
KU70
Flag-SMARCAD1-WT
Flag-NΔ-SMARCAD1
Relative mRNA expression
Vec
shFANCM
g
HU 1mM
No 2h 24h
γH2AX
KU70

**Fig. S9. Related to Fig. 6.**
**SMARCAD1 displaces 53BP1 at seDSBs on broken forks.**

a, Three PLA experiments of γH2AX and RAD51 (left) or γH2AX and 53BP1 (right) were performed in U2OS cells treated with IR (4 Gy) or HU (1 mM, 24h), with ~100 nuclei analyzed in each experiment. The average of three experiments and statistical significance are shown. Related to Figure 6a. (n=3 replicates)

b, Three PLA experiments of γH2AX and 53BP1, γH2AX and PCNA or γH2AX and Flag-PIF1 were performed after HU treatment (1 mM, 24h) in U2OS cells expressing Flag-PIF1 with depletion of SMARCAD1 by shRNAs (left). ~100 nuclei were analyzed in each experiment. The average of three experiments and statistical significance are shown. Western blot analysis was performed to show SMACRAD1 depletion in U2OS (EGFP-HR/STGC) cells using KU70 as a loading control (right). Related to Figure 6b. (n=3 replicates)

c, Three experiments of γH2AX and PCNA PLA (left), γH2AX and Flag-PIF1 (right) were performed after HU treatment (1 mM, 24h) in U2OS WT and *53BP1*-KO cells, with ~100 nuclei analyzed in each experiment. The average of three experiments and statistical significance are shown. Related to Figure 6c. (n=3 replicates)

d, Three experiments of 53BP1 and Flag-SMACRAD1-WT or Flag-NΔ-SMACRAD1 were performed in U2OS (EGFP-HR/STGC) cells with endogenous SMACRAD1 depleted by shRNA after HU treatment (1 mM, 24h) (left). ~100 nuclei were analyzed in each experiment. The average of three experiments and statistical significance are shown. Western blot analysis was performed to show the expression of Flag-SMACRAD1-WT or Flag-NΔ-SMACRAD1 using KU70 as a loading control (right). Related to figure 6d. (n=3 replicates)

e, Three experiments of PCNA and γH2AX PLA (left) and Flag-PIF1 and γH2AX PLA (right) were performed after HU treatment (1 mM, 24h) in U2OS cells expressing Flag-SMACRAD1-WT or Flag-NΔ-SMACRAD1 with endogenous SMACRAD1 depleted by shRNA. ~100 nuclei were analyzed in each experiment. The average of three experiments and statistical significance are shown. Related to figure 6e. (n=3 replicates)

f, Western blot analysis was performed to show the expression of Flag-SMACRAD1-WT or Flag-NΔ-SMACRAD1 in U2OS (EGFP-BIR/LTGC-Flex1) cells with KU70 as a loading control (top). qPCR was performed to show depletion of FANCM in these cells (bottom). Related to Figure 6g.

g. γH2AX Western blot analysis was conducted before and after treating U2OS cells with HU (1 mM, 2h and 24h) with KU70 as a loading control. Replication fork is stalled upon HU (1 mM, 2h) treatment and broken after HU (1 mM, 24h) treatment.

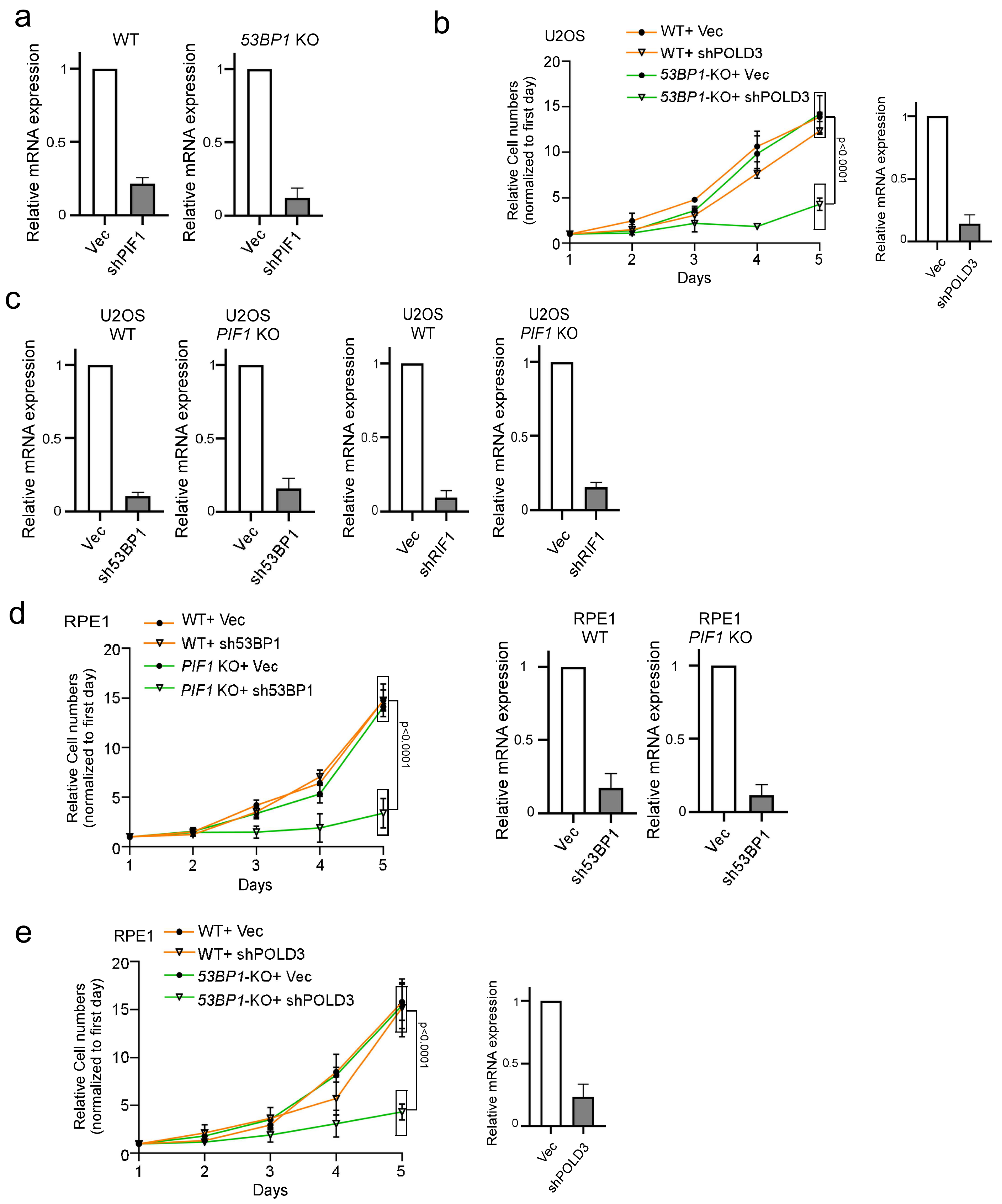


**Figure S10. Related to Figure 7.**
**53BP1-deficient cells rely on PIF1 for survival.**

a, qPCR was performed to show PIF1 depletion in U2OS WT or *53BP1*-KO cells. Related to Figure 7a.

b, The growth curves of U2OS WT and *53BP1*-KO cells were plotted after infection with lentiviruses expressing POLD3 shRNA with a vector control (left). qPCR was performed to show depletion (right).(n=3 replicates)

c, qPCR was performed to show depletion of 53BP1 or RIF1 in U2OS WT or *PIF1*-KO cells. Related to Figure 7b.

d, The growth curves of RPE1 WT and *PIF1*-KO cells were plotted after infection with lentiviruses expressing 53BP1 shRNA with a vector control (left). qPCR was performed to show depletion of 53BP1 in RPE WT or *PIF1*-KO cells (right).(n=3 replicates)

e, The growth curves of RPE1 WT and *53BP1*-KO cells were plotted after infection with lentiviruses expressing POLD3 shRNA with a vector control (left). qPCR was performed to show depletion of POLD3 in RPE WT or *53BP1*-KO cells (right).(n=3 replicates)

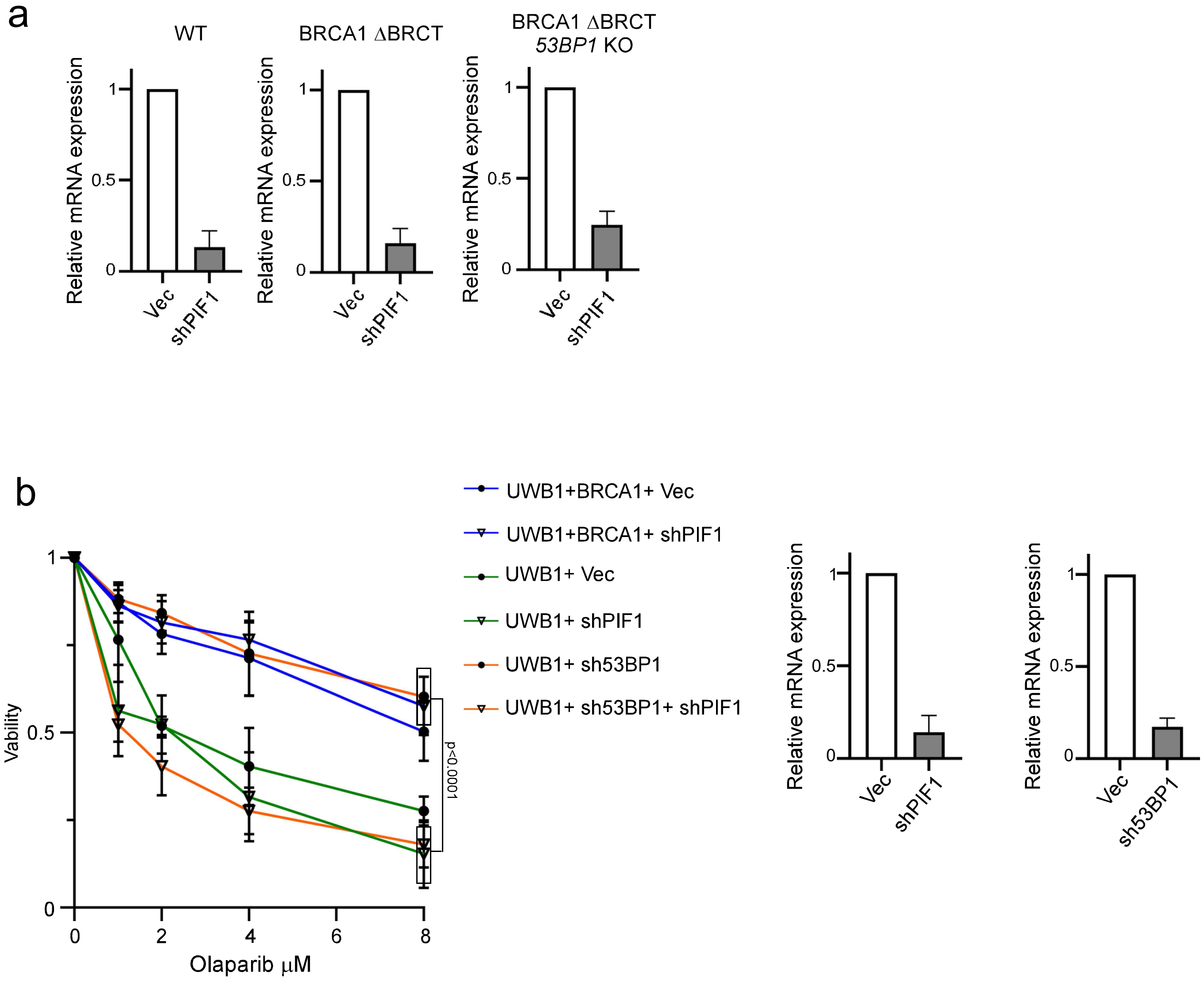


**Figure S11. Related to Figure 7.**
**Depletion of PIF1 overcomes the resistance to Olaparib due to 53BP1 inactivation in BRCA1-deficient cells.**

a, qPCR was performed to show PIF1 depletion in U2OS WT, BRCA1-ΔBRCT or *53BP1*-KO/BRCA1-ΔBRCT cells. Related to Figure 7c.
b, BRCA1-deficient UWB1 cells and BRCA1-reconstituted UWB1 (+BRCA1) cells were infected with lentiviruses encoding shRNAs for PIF1 or/and 53BP1 with a vector control, and cell viability was determined after treatment with the indicated concentrations of Olaparib for 72h (left, n=3 replicates). qPCR was performed to show depletion of PIF1 and 53BP1 (right).

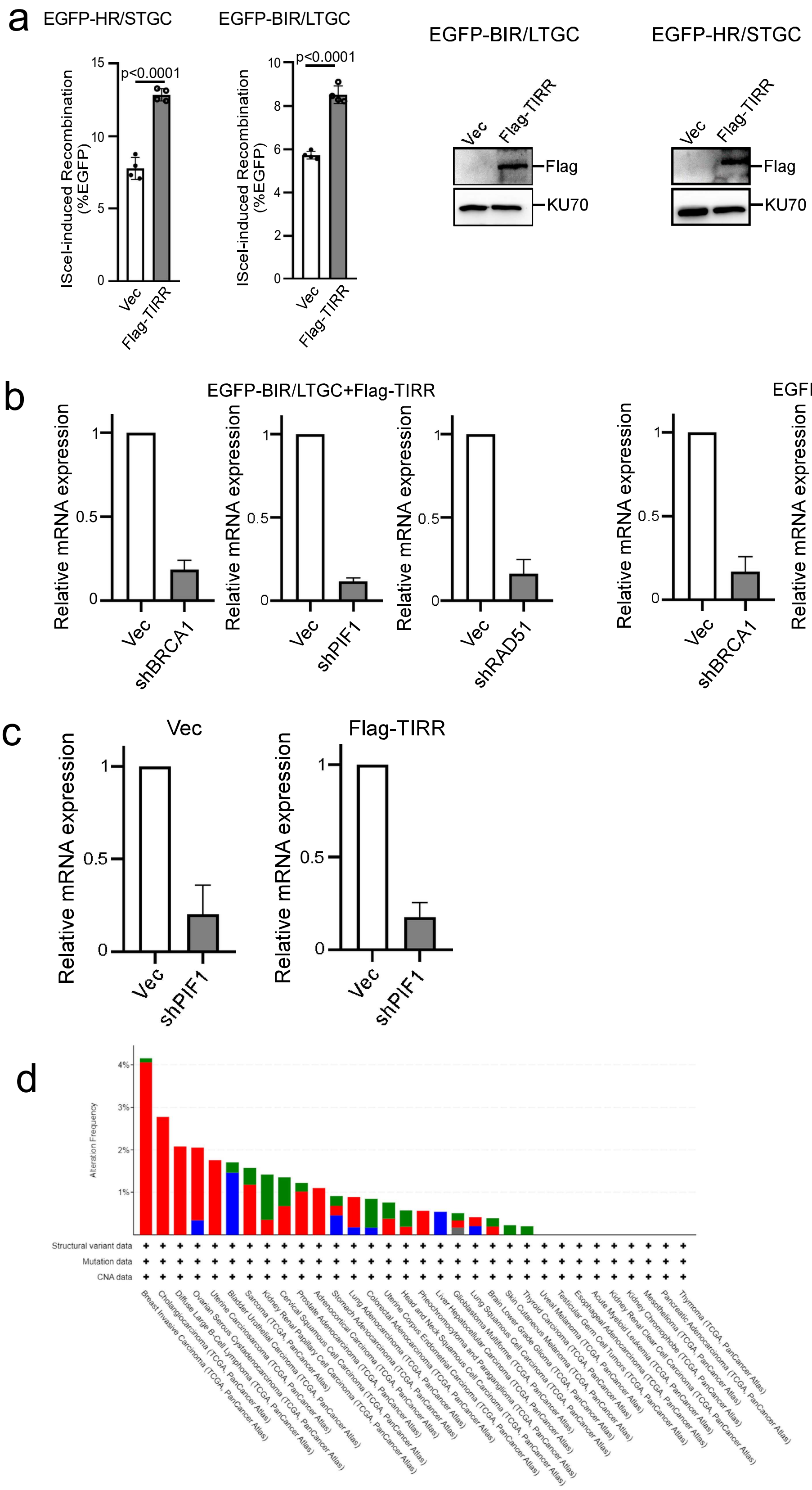


**Fig. S12. Related to Fig. 7.**
**Inactivation of PIF1 decreases viability of TIRR-overexpression cells.**
a, U2OS (EGFP-BIR/LTGC) cells and U2OS (EGFP-HR) cells were infected with lentiviruses encoding Flag-TIRR, and BIR or HR frequency was determined by the percentage of EGFP positive cells with FACS analysis, 5 days post infection of I-SceI lentiviruses (left). Western blot analysis was performed to show Flag-TIRR expression using KU70 as a loading control (right). (n=4 replicates)
b, qPCR was performed to show depletion of BRCA1, PIF1 and RAD51 in U2OS (EGFP-BIR/LTGC) (left) and U2OS (EGFP-HR/STGC) (right) cells expressing Flag-TIRR. Related to Figure 7d.
c, qPCR was performed to show depletion of PIF1 in U2OS cells with or without expressing Flag-TIRR. Related to Figure 7e.
d, Analysis of TIRR alternations in cancers using the cancer genomics data sets through cBioPortal (https://www.cbioportal.org/). Amplification (red), deep deletion (blue), mutation (green) or multiple alterations (grey) are indicated.

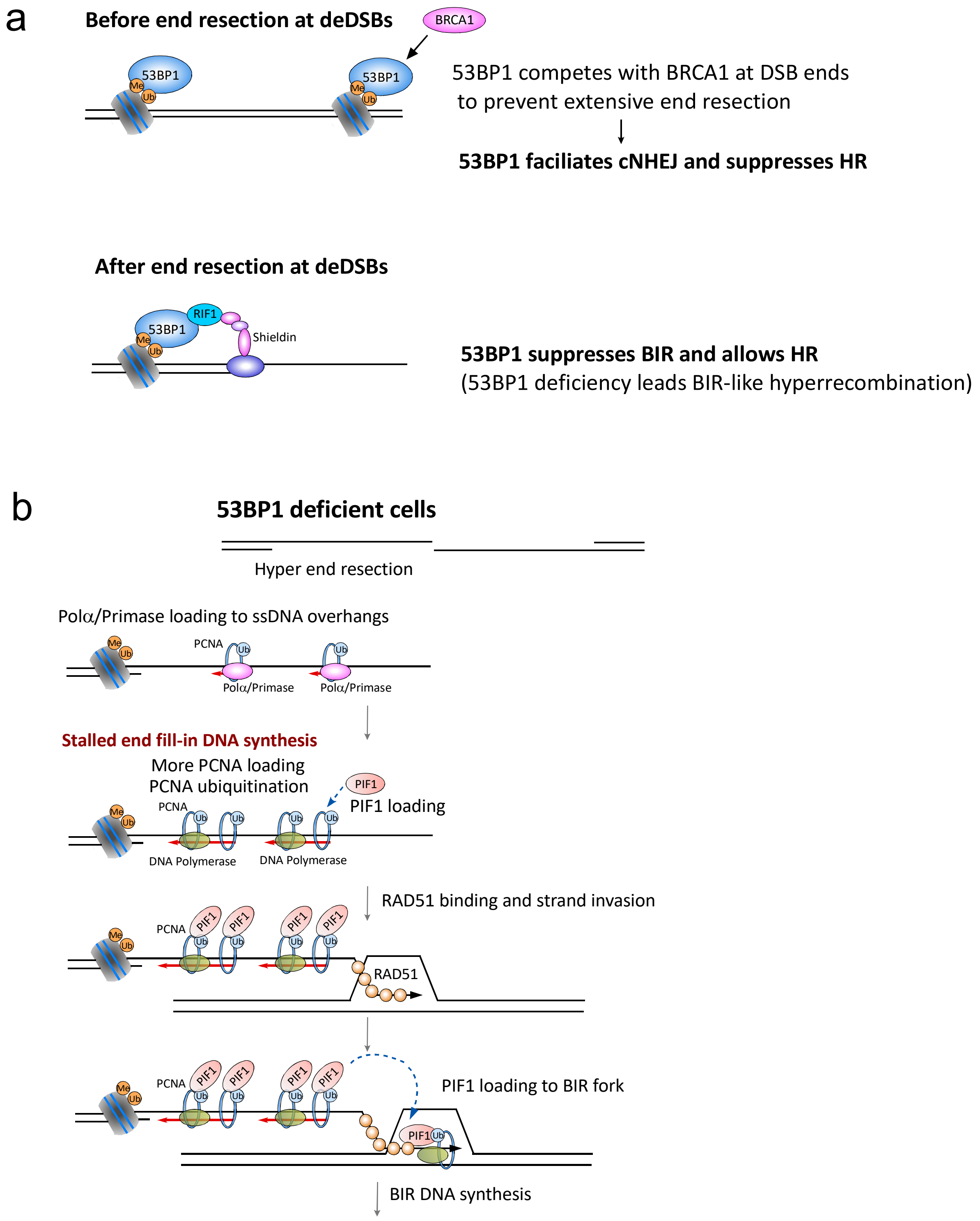


**Fig. S13**

**The role of 53BP1 in DSB repair pathway selection of NHEJ versus HR before end resection and HR versus BIR after end resection.**

a, Schematic drawings illustrating competition for DSB ends by 53BP1 and BRCA1 before end resection at deDSBs to favor cNHEJ over HR (top), and the control of the 53BP1 pathway to facilitate HR over BIR after end resection at deDSBs (below).

b, Illustration of PCNA ubiquitination and PIF1 recruitment due to fork stalling during end fill-in DNA synthesis on ssDNA overhangs at DSBs in 53BP1-deficient cells. We propose that after RAD51-mediated stand invasion, enriched PIF1 in association with ubiquitinated PCNA on ssDNA overhangs would be loaded to the 3' invading strand to promote BIR replisome assembly and BIR activation.

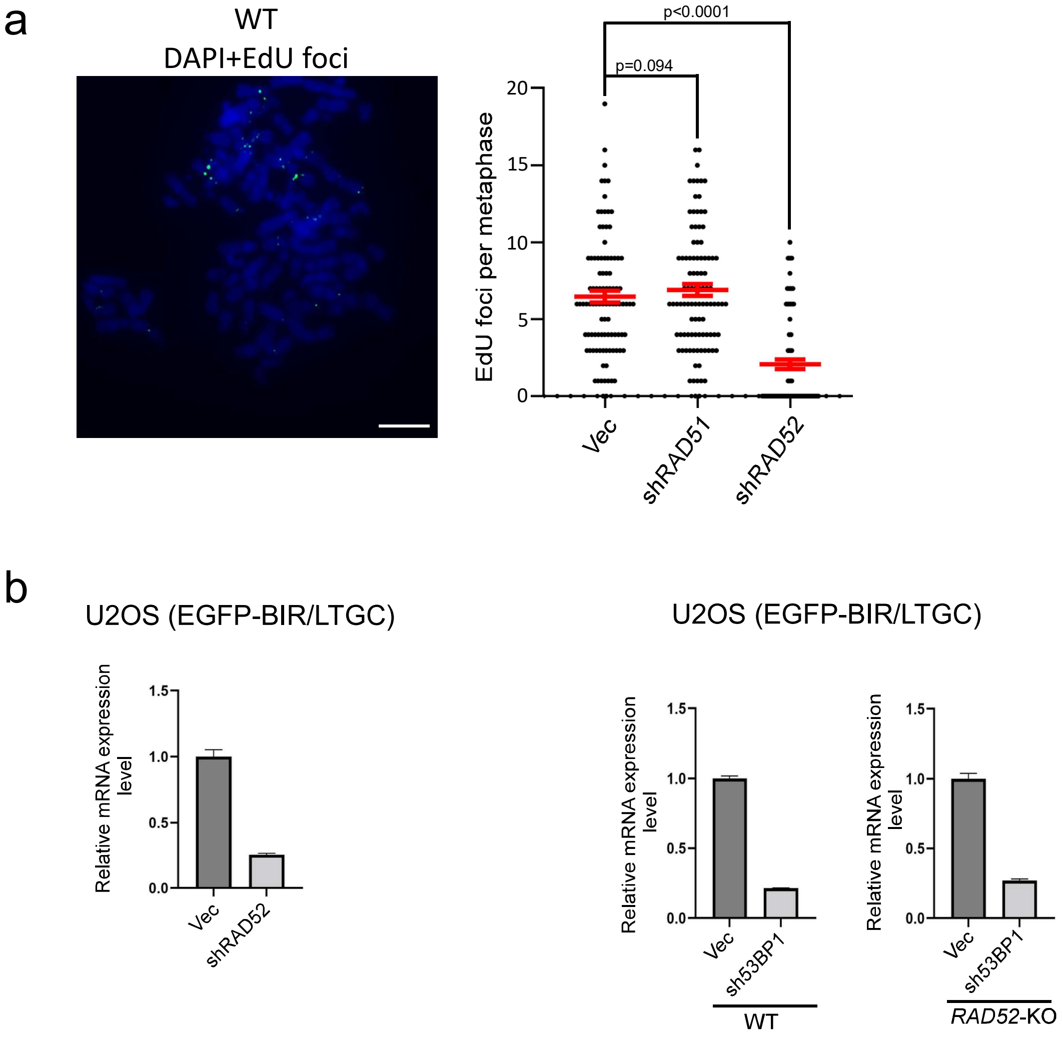


**Figure S14.**

**MiDAS and BIR in cycling cells show different dependencies on RAD51 and RAD52.**

a, MiDAS analysis was performed in U2OS cells after depleting RAD51 and RAD52 with a vector control. A representative MiDAS image illustrating EdU incorporation on metaphase chromosomes is shown (left) and EdU foci on each metaphase spread are quantified (right). ~100 metaphase spreads were analyzed in each sample. Scale bar = 10 μm.(n= 100 metaphases)

b, qPCR was performed to show RAD52 depletion in U2OS (EGFP-BIR/LTGC) cells (left) and 53BP1 depletion in U2OS (EGFP-BIR/LTGC) WT and *RAD52*-KO cells (right). Related to Figure 1e.

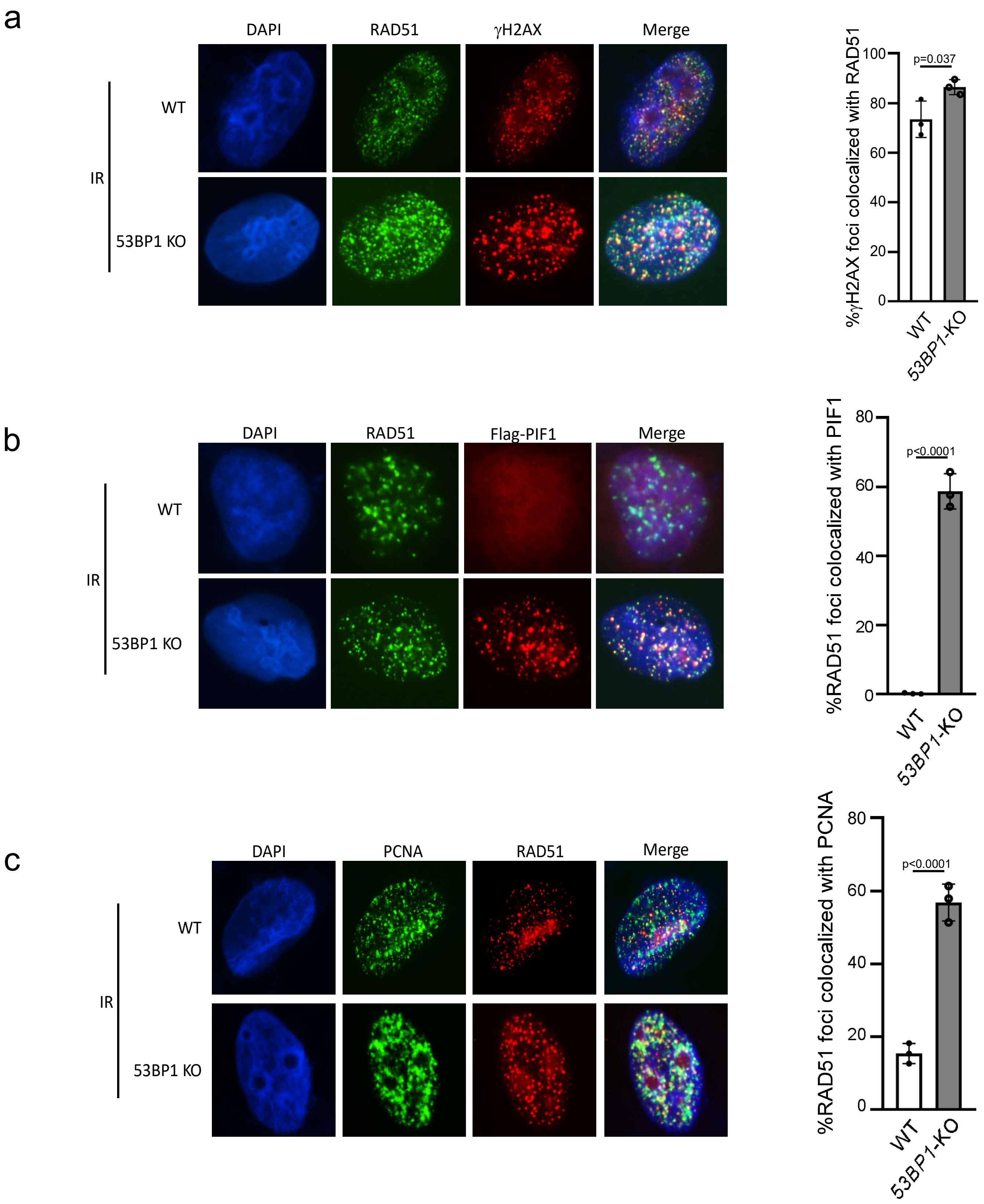


**Fig. S15**

**Colocalization of RAD51 with PIF1 and PCNA is increased in 53BP1-KO cells after IR.**

U2OS WT and *53BP1*-KO cells stably expressing Flag-PIF1 were irradiated with 4-Gy IR, fixed at 2hours after irradiation, followed by co-immunostaining of RAD51 (green in a and b, red in c) with γH2AX (red) (a), Flag (Flag-PIF1, red) (b) or PCNA (green) (c), using DAPI for DNA staining (blue). The yellow dots represent colocalization between PIF1 and γH2AX (left). Quantification of colocalization of RAD51 with γH2AX, Flag-PIF1 or PCNA is shown (right). (n=3 replicates)

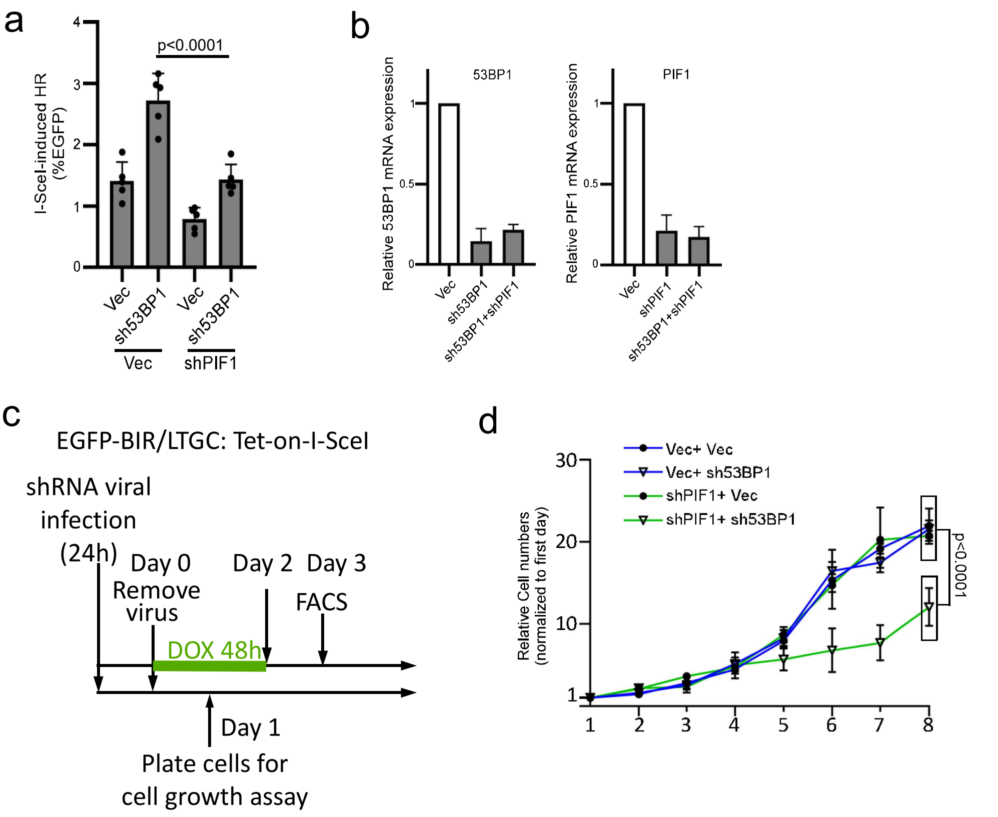


**Fig. S16**

**PIF1 is required for increased BIR in 53BP1-deficient cells.**

**a,** U2OS (EGFP-BIR/LTGC-tet-on I-SceI) cells were infected with lentiviruses encoding shRNAs for 53BP1 and PIF1 with a vector control, followed by inducing I-SceI expression with doxycycline (DOX, 2 μg/ml) at Day 0 (see **c**) for 48 hours. FACS analysis was performed 1 days after removal of DOX (Day 3).(n=5 replicates)

**b,** qPCR analysis was performed to determine the shRNA knockdown efficiency of 53BP1 and PIF1 in U2OS (EGFP-BIR/LTGC-tet-on I-SceI) cells .

**c,** The timeline of the key steps for the BIR assay using the Tet-on-I-SceI system and the cell growth assay.

**d**, The growth curves of U2OS (EGFP-BIR/LTGC-tet-on I-SceI) cells were plotted after infection with lentiviruses expressing shRNAs for 53BP1 and PIF1, with a vector control (Vec). The cell plating date is indicated in **c**.(n=3 replicates)
